# Evidence of Chemical Wave-Electric Field Interaction in Bacterial Cells

**DOI:** 10.64898/2026.09.05.749092

**Authors:** Jie-Pan Shen, Chia-Fu Chou

**Affiliations:** Institute of Physics, Academia Sinica, Taipei 11529, Taiwan; Research Centre for Applied Sciences, Academia Sinica, Taipei 11529, Taiwan

## Abstract

Charge neutrality is widely assumed in living cells, yet this approximation breaks down in micron-scale bacteria where charge imbalance and spatial confinement are significant. Using Poisson–Nernst–Planck modeling, we show that unequal cation–anion effectiveness and bounded geometry generate extended intracellular diffuse layers and steady electric fields. We demonstrate that such fields couple directly to intracellular chemical waves, focusing on the Min-protein oscillator of *Escherichia coli*. Electric-field–driven transport skews the dispersion-mode structure, induces mode crossings, and selectively amplifies Turing and Hopf–Turing instabilities over intermediate length scales, constraining the permitted ω–k spectrum and setting optimal wavelengths and modal growth-rate velocities. Experiments in wild-type, anucleate, and antibiotic-treated cells, together with simulations of nucleoid-dependent charge density and field strength, quantitatively validate these predictions and explain observed pattern asymmetries and frequency modulations. Crucially, asymmetric wave–field coupling promotes quasi-periodicity through controlled mode competition, enhancing robustness to noise, cell-size variation, and growth. These findings identify intracellular electric fields as active regulators of biochemical patterning and suggest a general role for wave–field interactions in cellular self-organization.

## Background

Chemical waves arise from nonlinear chemical reactions and diffusion processes in both chemical and biological systems (*1–5*). These waves emerge when the concentration of reactants or products changes in a manner that propagates through space and time, self-organizing into patterns that often resemble traveling waves. A notable example of chemical waves is the Belousov-Zhabotinsky (BZ) reaction, a classic oscillatory chemical system (*1*). When an electric field (E-field) is applied to this system, it gives rise to various wave phenomena, and the interactions between chemical waves and E-field have been extensively studied both theoretically (*6–9*) and experimentally (*10, 11*). However, such interactions have not been observed in biological systems because the insulating property of cell membrane avoids external E-field and bulk electroneutrality prohibits intracellular E-field in biological media. Intriguingly, it has been shown that the dissipation of membrane potential in bacteria disrupts the chemical waves self-organized by the Min-protein system — a spatiotemporal coordinator in cell division (*12*), raising the question of whether a sufficient E-field could be generated within the cellular context to influence the Min-protein oscillator in bacteria.

Electroneutrality is a basic assumption in classical electrolyte theory, shaping the key concepts in electrochemistry and electrophysiology (*13*). It suggests that charge separation or excess charge does not occur within the ionic solution, ensuring that neutrality is maintained throughout. Coulomb forces between ions in an electrolyte solution work to preserve macroscopic electroneutrality, as any spatial charge imbalances in the conventional bulk are immediately neutralized by ion movement to minimize the electrochemical (ECM) potential. At charged interfaces, the electrolyte compensates for surface charges by forming electric double layers (EDLs), within which are not electroneutral. In electrostatic equilibrium, these double layers prevent potential differences between the interfaces and prevent the formation of a macroscopic E-field in the conventional bulk (Fig. 1A). However, this physical understanding may be challenged when the assumptions underlying electroneutrality are invalid, especially considering that an E-field in an electrolyte solution can only exist due to net charge separation, creating an inherent paradox in the principle (*14*). For dynamic problems, such as ion channel-mediated current around the cell membrane (*15, 16*) and non-stationary liquid junction potential during the continued dialysis (*17*), emergence of excess charge at the interface generates a local E-field and induces extension of diffuse layer away from the interface. Alternatively, for steady-state problems such as electrochemical cells with liquid electrolytes, 1-2 nm range of EDL around the cell membrane and electroneutral bulk are derived by Poisson–Boltzmann (PB) equation based on the Gouy–Chapman theory when electroneutrality assumption is maintained at infinity and cytosolic anionic/cationic charges are equal in the conventional bulk (*13*). Whereas, the validity of both conditions may be questioned for micron-sized bacterial cells.

**Fig. 1.**
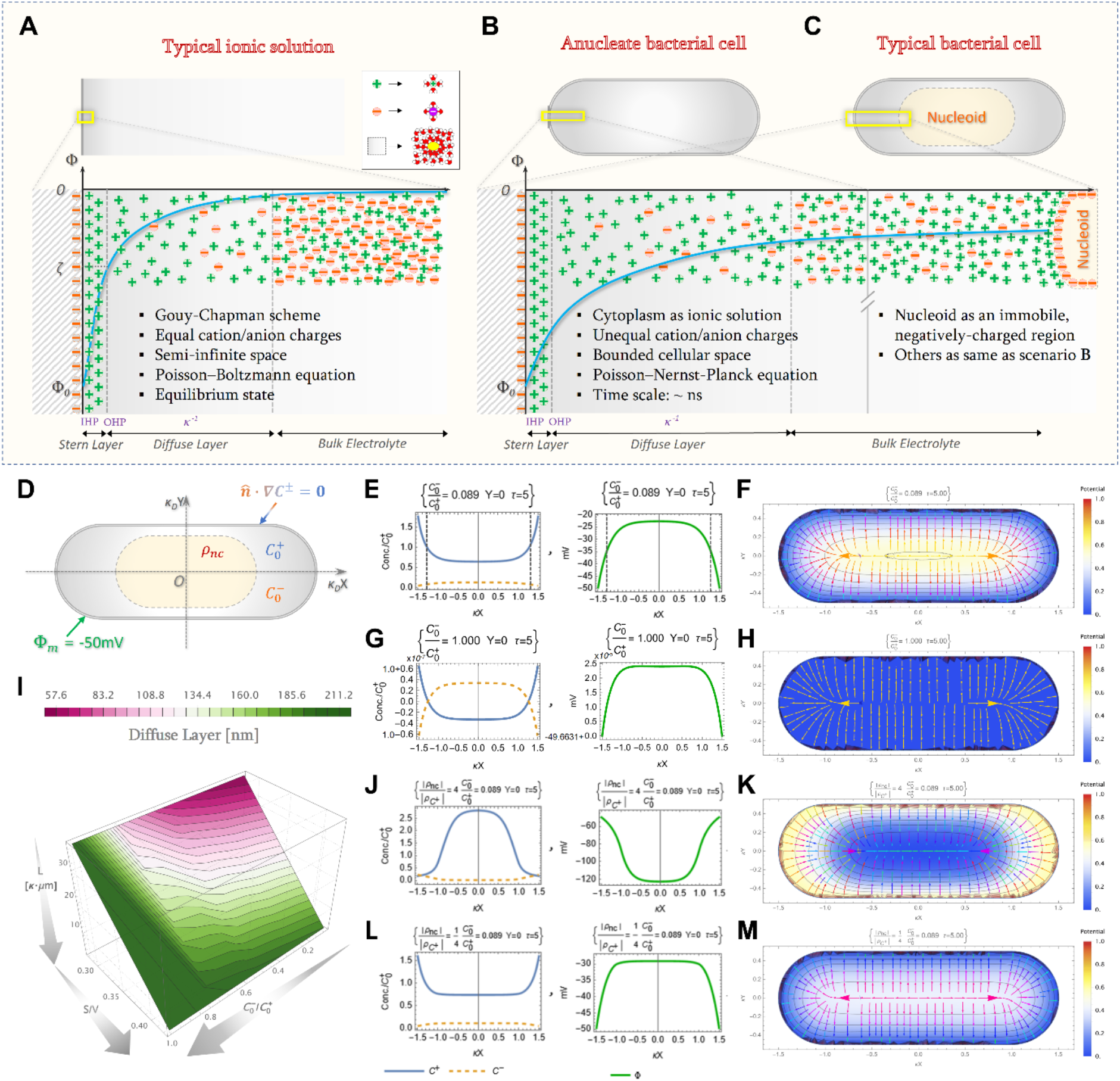
The extension of diffuse layer in bounded cellular space with differential cation/anion ratios and the configuration of electric field altered by the charge density of nucleoid. (A-C) Schematics for the physical scenarios of (A) typical ionic solution, (B) anucleate bacterial cell and (C) normal bacterial cell, leading to the region beyond the electric double layer conceptualized as the conventional (A), and cytosolic bulk (B, C), respectively. (D) Schematics for 2-dimensional projected rod-shaped cell model. The light orange region at the cell center represents the nucleoid. The Dirichlet and Neumann boundary value conditions are shown and set at cell boundary. (E and G) The longitudinal concentration profiles [left panels] and electric potential [right panels] for the anion/cation charge ratio = 0.089 (E) and 1.000 (G). The longitudinal concentration profiles of anions/cations (left panel), which determines the electric potential profile (right panel), show an exponential decay from both boundaries to reach a basin in the central region, indicated by two vertical black dash lines in (E) where the 1/e drop relative to the maximum profile at the boundary defines the cytosolic bulk. (F and H) The electric field (streamline arrows) and potential (contours) for (E) and (G). (I) The extension of diffuse layer in relation to surface-to-volume ratio (S/V), cell length (L) and cation/anion ratio *(C^−^_0_/C^+^_0_)*. (J and L) The longitudinal concentration profiles [left panels] and electric potential [right panels] for the charge density ratio between the nucleoid and the total cations = 4 (J) and 1/4 (L) when the anion/cation charge ratio = 0.089. (K and M) The electric field (streamline arrows) and potential (contours) for (J) and (L). IHP: Inner Helmholtz plane. OHP: Outer Helmholtz plane. Φ: surface potential. ζ: zeta-potential. *κ*^-1^: length of diffuse layer; equivalent of Debye length (*κD*^-1^ = λD) in (A). C^+^_0_: total cation concentration. C^−^_0_: total anion concentration. ρnc: charge density of the nucleoid. *n*$: the unit vector normal to cell boundary.

Notably, the concentrations of key intracellular electrolytes of *Escherichia coli* during its growth phase have been experimentally determined (see Supplementary Information), indicating the intracellular concentration of K^+^ constantly surpasses that of other cations, as well as that of anions like Cl^-^, Pi and metabolites. When considering ion hydration, for thick, highly structured hydration shells formed through extensive hydrogen bonding, as in glutamate, strongly attenuate intrinsic electric fields compared to ions with thinner hydration layers such as K⁺. Therefore, not only the total effective anionic/cationic concentration, but also the total effective charges carried by anions/cations are substantially imbalanced in bacterial cytosol, leading to a significant deviation from bulk electroneutrality as this charge-imbalanced electrolyte solution is enclosed in a finite, subcellular space invalidating unlimited supplies of counterions to balance the excess charge in the cytosolic bulk, wherein excess charge may exist due to imbalanced charges of anions/cations in the region away from the cell membrane (Fig. 1B). Further, those negatively-charged macromolecules allocated to the nucleoid, such as DNA, RNA, and assorted metabolites (nucleotides and nucleoid-associated proteins), are much less mobile compared to electrolytic background, at least to the time scale for electrolyte redistribution to achieve equilibrium, acting like an immobile negatively charged region over the central part of bacterial cell (Fig. 1C). Taken these contexts together, the physical conditions to support the electroneutrality assumption are not evidently justified in the cytosolic bulk of micron-sized bacteria, implicating the possibility of intracellular E-field generated by excess charge in the cytosolic bulk and asymmetric charge distribution in EDL. The latter may also bring an extended diffuse layer around the cell membrane much longer than typical Debye length (*16*).

To better address these discrepancies, we employed the Poisson-Nernst-Planck (PNP) equations which, when solved numerically, can provide a more accurate representation of ionic behavior in biological systems without relying on a priori assumption of electroneutrality (*18, 19*). Furthermore, rather than becoming distracted by the multitude of interpretations proposed for biomolecular mobility and segregation in bacterial cells (*20–23*), the chemical wave– electric field interaction offers another experimentally discernible analysis that can provide further evidences linked to the influence of intracellular E-field on biological processes. The Min-protein oscillator, consisting of MinC/D/E proteins, is a well-known chemical wave system (*24*) involved in bacterial cell division. This system self-organizes into traveling waves and oscillations. As over 70% of the proteins carry negative charge at physiological pH (*25*), the interaction between these proteins and the intracellular E-field can influence the wave properties of the Min-protein system, including frequency modulation of the oscillation. In this study, we present both analytical and numerical results that explore (1) the intracellular E-field configuration in rod-shaped bacteria, (2) the modal structure of the dispersion relation in response to the wave-field interaction, and (3) experimental verification of the dispersion relation and the frequency modulation of Min-protein oscillations, attributed to changes in nucleoid morphology and associated charge density. These findings offer valuable insights into the interaction between intracellular E-field and biochemical processes in bacteria.

## Results

### The extension of diffuse layer and the configuration of electric field in bacteria

Based on the PNP equations, the electrochemical models for 2-dimentional projected geometries of rod-shaped bacteria in various sizes (Fig. 1D and details in Supplementary text) are constructed to better understand the spatial profile of main electrolytes and the associated configuration of E-field in bacteria. The numerical results show the electrochemical gradients along the longitudinal axis of rod-shaped bacteria in various sizes are built up by imbalanced charge distribution with differential anion/cation charge ratios wherein cation concentration is set to 231.25 mM (Figs. S1-S7). Considering a 2.727 μm (≈ 3000· λ_D_ nm, where λ_D_ = *κ_D_*^-1^ ≈ 0.91 nm as the Debye length for 231.25 mM of ionic solution) long cell with anion/cation charge ratio ranging from 0.089 to 1.000, the longitudinal length of chemical gradient across from the cell pole to the cytosolic bulk is up to 0.218 μm (Fig. 1E, 1G and S1), suggesting that the diffuse layer extends much longer than that derived from the PB-based approximation in typical ionic solution. While in the cell context with *equally* effective anionic/cationic charges, the PNP-based approximation turns out the cytosolic bulk almost electroneutral (∼2.5·10^-5^ mV) with respect to the cell boundary (Fig. 1G-H), denoting a close confluence of the PNP and PB-based approximations in typical ionic solution. Whereas, the buildup of pronounced ECM potential is up to 28 mV ascribed to excess charge between cations and anions in the cytosolic bulk as anion/cation charge ratio = 0.089 (Fig. 1E-F and S2). Accordingly, our numerical results indicate that the extension of diffuse layer depends on the degree of charge separations in the cytosolic bulk caused by *unequally* effective anionic/cationic charges bounded in a micron-sized cell.

To further abstract the key factor leading to the extended diffuse layer in response to imbalanced charge distribution, changes in cellular confinement sizing from 2.727 µm to 34.125 µm and anion/cation charge ratio from 0.089 to 1.000 are implemented in the numerical experiments. Interestingly, the diffuse layer, except for balanced anionic/cationic charge, is found reduced from 174.72∼188.37 to 44.59∼91.91 nm for anion/cation charge ratio 0.089∼ 0.772 when cellular confinement increases from 2.727 to 34.125 µm (Fig. 1I and S8). Besides, for a constant cellular confinement and cation concentration, the diffuse layer increases when anion/cation charge ratio increases from 0.089 to 1.000. The effects of cellular confinement (or cell size) and anion/cation charge ratio upon the extended diffuse layer can be reduced to another key factor, the surface/volume (S/V) ratio, to bring out a general picture in determining the diffuse layer. Since the membrane surface is negatively charged, part of cations in the cytosol are attracted to the membrane for neutralizing the surface charges. The leftover cations in the cytosol then redistribute to balance anions to reduce the ECM potential due to excess cations. This ionic redistribution results in a maximal cytosolic bulk, by which the minimized charge density can be achieved for minimizing excess charge per volume, leading to the minimal electric potential and thereof minimal deviation of electroneutrality. Therefore, the transition region between the membrane and the cytosolic bulk is formed and known as the diffuse layer. It is apparent when the cell size reduces, the S/V ratio increases. In this scenario, a smaller proportion of cations is left to form the cytosolic bulk after neutralizing surface charges on the membrane, and a smaller cytosolic bulk and a more extended diffuse layer will be formed. Alternatively, when the S/V ratio (or cell size) and the cation concentration are fixed, more cations in the cell solution are available to balance anions if anion/cation charge ratio is smaller, leading to a larger cytosolic bulk and a smaller diffuse layer.

The configuration of E-field, in contrast to being outwardly directed to the cell boundary in anucleate cells (Fig. 1B, F and H) due to the presence of excess cations in the cytosol, appears to get acclimated by the regional, negatively-charged nucleoid in a cell (Fig. 1C, J-M). For example, the E-field becomes inwardly directed from the cell boundary when the charge density ratio |ρ_nc_⁄ρ_C_+ | between the nucleoid (*nc*) and cytosolic cations (*C^+^*) equals to 4 (Fig. 1J-K), while outwardly directed when |ρ_nc_⁄ρ_C_+ | = 1⁄4 (Fig. L-M). The nucleoid charge density in a cell is not a constant partly because the nucleoid size and the associated molecular amounts change in different stages of cell cycle. Since the ratio of nucleoid size over the cell size, known as nucleocytoplasmic ratio (NC ratio), is almost constant (*26*) such that the nucleoid size, as well as its charge density, varies as cells under vegetative growth. Given a single copy of chromosome in a 3.789 µm long cell, |ρ_nc_⁄ρ_C_+ | is about 1/11 (or 1/14). Whereas, up to 8 copies of chromosomes due to DNA duplications has been reported in different growth stages (*27*), and as such the nucleoid charge density, as well as the strength and direction of E-field, are altered accordingly. Our numerical experiments indicates that, apart from inward/outward direction of E-field, subtle variations in field configuration emerge around the cell poles as changes in nucleoid size alter the ratio |ρ_nc_⁄ρ_C_+| during cell growth (Figs. S9-11), suggesting a potential role in biomolecular transport, and conceivably wave-field interactions. As electrostatic mechanisms have been shown to play important roles in charge-dependent protein mobility, segregation of subcellular biomolecules into specific regions and formation of protein-ribosome coacervate compartment in bacteria (*20–23, 28*), the potential interactions between chemical waves and intracellular E-fields in living cells remain largely unexplored and warrant further investigation.

### The novel wave phenomena in bacterial Min-protein oscillator

Dispersion relations have been used in the BZ system to reveal how E-fields shape wave phase dynamics and give rise to new wave behaviors (*8, 9*). In bacterial Min-protein system, MinD and MinE are commonly described as an activator–inhibitor pair within a reaction–diffusion framework, with their roles defined by membrane binding (Fig. S12) (*24*). Building on this framework, we first attempt to apply a similar dispersion-based analysis to investigate how intracellular E-field influences Min-protein wave behavior through cytosolic nucleotide exchange—the only Min-protein reaction occurring in the cytosol—while neglecting membrane-associated reactions. The wave mode under study can be approximated by *ω(k) ~ ω_c_(1 − μ(E^⇀^ • k^⇀^) + Δk^2^)*, where the frequency *ω*(*k*) of Min-protein waves relates to the wavenumber *k* by a parabolic function. The linear term is introduced here by the wave-field interaction and the quadratic term is caused by the reaction/diffusion of free-diffusing reactants (*8*). The spatial averaged term μ(E^⇀^ • k^⇀^) in the wave-field interaction is originated from the coupling of Min-protein waves with wavevector k^⇀^ propagating across a nonuniform, intracellular E-field E^⇀^ in a cell. Parameters μ and Δ relate to electric mobility and phase diffusion owing to the reactions over a limit-cycle period, 2*π*⁄*ω_c_* (see details in Section C in SI). Since the fastest wave mode is constrained by the maximal and minimal dispersion branches, the frequency of any Min-protein oscillator cannot exceed these bounds. With nucleoid charge density varying across cellular phases (see Appendix in SI) and a longitudinally symmetric intracellular field of the rod-shaped cell, the field strength may adjust accordingly and its mean value is estimated ca. 13.82 mV/μm for 2.727 μm long cell (|ρ_nc_⁄ρ_C_+| = 1⁄16 in Fig. S11), placing the permitted frequencies of Min-protein oscillator— corresponding to specific cell lengths *π*⁄*k*—within the regime defined by *1 ± μ(E^⇀^ • k^⇀^) + Δk^2^* space (Fig. 2A), where ± sign captures the difference in how waves interact as they traverse through each half of the intracellular electric potential well. In the field-free limit, wave modes are restricted to the dispersion 1 + Δ*k*^2^ in *ω* − *k* space, whereas experimental measurements of Min-protein oscillation frequency and cell length in growth-phase *E. coli* cells (open circles in Fig. 2A; *n* = 2450) extend across the broader regime predicted by theory and simulations (mean field strength for various cell lengths in Fig. 2B; other parameters estimated in Section C in SI), suggesting a possible role for intracellular E-field in Min-protein dynamics.

**Fig. 2.**
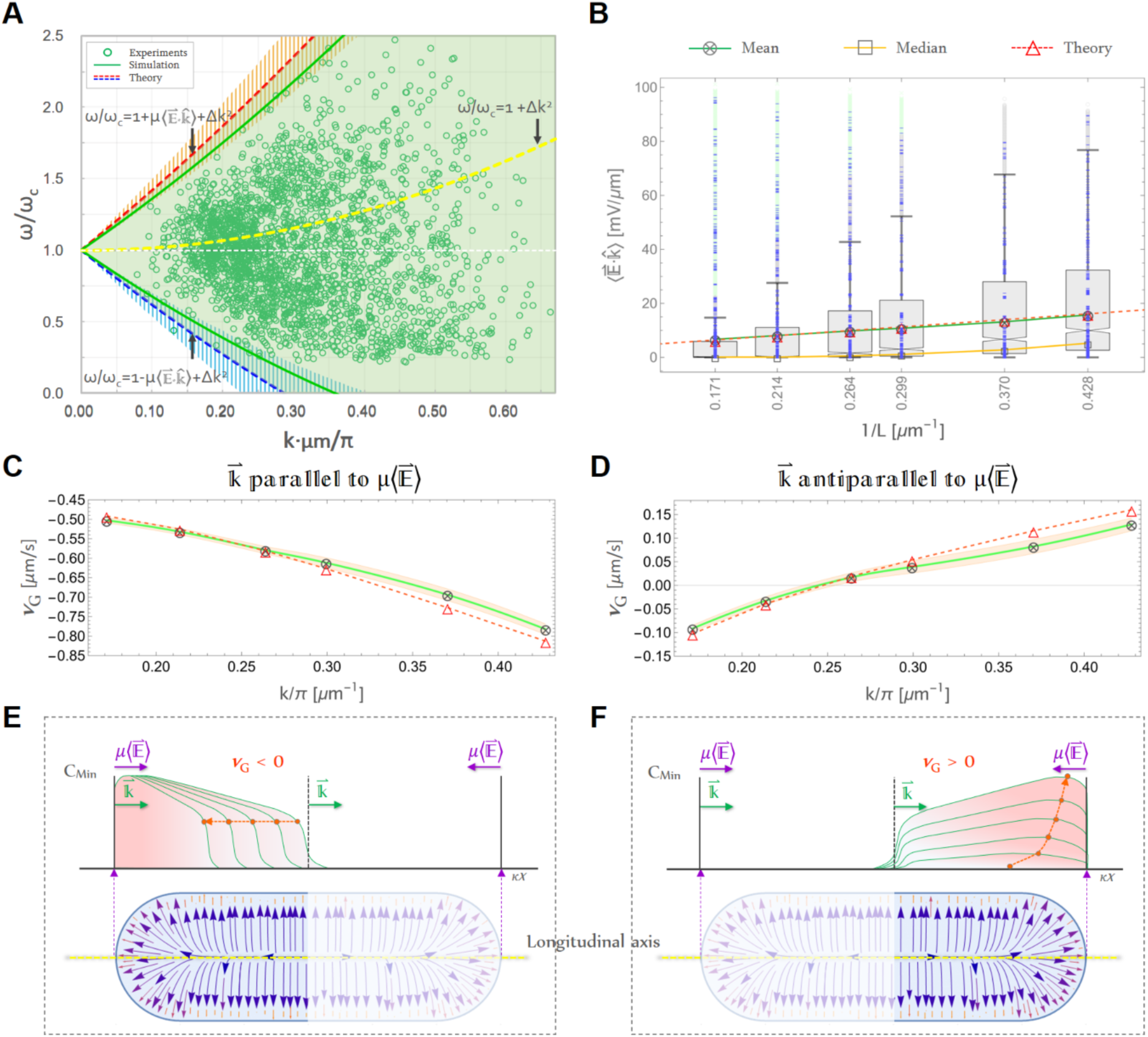
The E-field driven dispersion branches for cytosolic reaction wave mode and bacterial Min-protein dynamics. (**A**) The dispersion relation in the ω-k plane. ωc: mean oscillation frequency. Red/blue dash line: the upper/lower dispersion branches computed by theoretical prediction. Red/blue hatch area: the Std. Dev. interval of the upper/lower dispersion branches computed by theoretical prediction based on the values of integral electric mobility μ over a period Tc obtained from zone electrophoresis for large-scale proteomics (*39*) (see Section C in SI). Green line: the upper/lower dispersion branches computed by numerical simulation. Green open circles: experimental data. Yellow dash line: theoretical dispersion relation in the absence of E-field. (**B**) The diagram of mean E-field strength in relation to the inverse of cell length *L=π*⁄*k*. (**C-D**) The diagrams of growth-rate velocity when (**C**) *k^⇀^* parallel to *k^⇀^* and (**D**) *k^⇀^* antiparallel to *μ(E^⇀^)*. (**E-F**): schematics for the retrograde/growth process of Min-protein pattern in relation to growth-rate velocity ***vG***, illustrated in (**C-D**). Wavevector ***k^*** moves from the left to right pole and the response of negatively-charged Min proteins to the mean E-field, *μ(E^⇀^)*. The orange dots and the connected arrow lines are drawn to indicate the phase movement for a specific point on the wave profile in the longitudinal direction.

Next, to clarify how the intracellular E-field shapes the modal structure of the dispersion relation for bacterial Min-protein oscillator, linear stability analysis must incorporate both free-diffusing and membrane-associated reactions, capturing wave-mode evolution under E-field– driven transport. Intracellular E-field–driven transport breaks wave symmetry along the longitudinal direction. Given the demonstrated charge-dependent localization of biomolecules to specific intracellular regions in bacteria (*23*) suggests the possibility that intracellular E-field generated by the extended diffuse layer surrounding the cell membrane may contribute to the electrostatic organization of charged biomolecules, leading to their confinement within distinct spatial distributions or the establishment of equilibrium. Consequently, direct E-field–driven transport approaches an equilibrium, and small concentration perturbations experience effectively imaginary transport, giving rise to oscillatory, coupled behaviors—instabilities or sustained waves—while real transport is largely damped. The transport term is thus defined by *(E^⇀^•M∇C_j_) ≡ (iβ^⇀^•∇C_j_)*, with β = (β^⇀^ • x^~^), where *M* is electric mobility, x^ is unit vector along the gradient direction and *i* is imaginary number (see Section D in SI for details). Notably, waves propagate parallel to the field when *β* > 0 and antiparallel when *β* < 0. As such, the linear stability analysis of the bacterial Min-protein oscillator is formulated using reaction–diffusion equations incorporating an effective imaginary transport term, employing the ansatz *C_j_ = C^*^_j_ + δC_j_ • e^ik’x’+λt’^*, where *C^*^_j_* denotes equilibrium state and primed variables are dimensionless. For analytical tractability, a rectangular geometry is adopted in place of a rod-shaped cell. Min-protein oscillations in microfluidics-resculpted rectangular cells display wave dynamics comparable to those in rod-like cells (*29*), indicating that this geometry captures relevant biophysical features. Exploiting longitudinal symmetry, Min-protein waves are modeled in a 2D rectangular domain bounded by a reactive membrane, with Min proteins diffusing freely in the interior and interacting upon membrane binding.

Dispersion-based analysis (see Section D in SI for details) reveals five eigenmodes, each displaying distinct wave patterns and associated growth/decay or oscillatory behavior. As shown by the dispersion diagram for a 2D rectangular cell (L×H = 4 μm×1 μm) with *β* = −11.83, 0 and 11.83, the real part of λ(k′) (Re[λ(k′)]; Fig. 3A–B) transitions from negative (*β* ≤ 0) to positive (*β* > 0), while the imaginary part (Im[λ(k′)]; Fig. 3C-E) exhibits one pair of counter-propagating modes for *β* ≤ 0 and two pairs for *β* > 0, with symmetry about Im[λ(k′)] = 0 in each pair (Fig. 3H & J). For a given *β*, the envelopes of the real components of the five eigenmodes delineate a parabolic-like region (Fig. 3A). Remarkably, in the absence of intracellular E-field (*β* = 0), Re[λ(k′)] is negative for all wave modes, meaning that even waves with Im[λ(k′)] ≠ 0 cannot spontaneously arise without external stimulation, as they eventually decay. Such external stimulation, however, is unlikely under realistic cellular conditions. While mathematically valid, these waves are physically unrealizable. Only when waves propagate parallel to the E-field (*β* > 0) does Re[λ(k′)] become positive (Fig. 3A), allowing wave growth and self-emergence. This requirement for E-field-driven transport underscores the essential role of the intracellular E-field in bacterial Min-protein oscillator. It is therefore instructive to examine the wave modes for *β* > 0: Modes 2–3 (green and blue curves in Fig. 3G-H) and Modes 4–5 (cyan and magenta curves in Fig. 3I-J) each constitute a pair of counter-propagating modes. Mode 2-3 predominantly exhibits a Turing pattern (Re[λ(k′)] > 0 and Im[λ(k′)] = 0) across multiple wavenumber bands (Fig. 3K-L), whereas Mode 4-5 is dominated by a Hopf-Turing pattern (Re[λ(k′)] > 0 and Im[λ(k′)] ≠ 0) over specific wavenumber band (Fig. 3M-N).

**Fig. 3.**
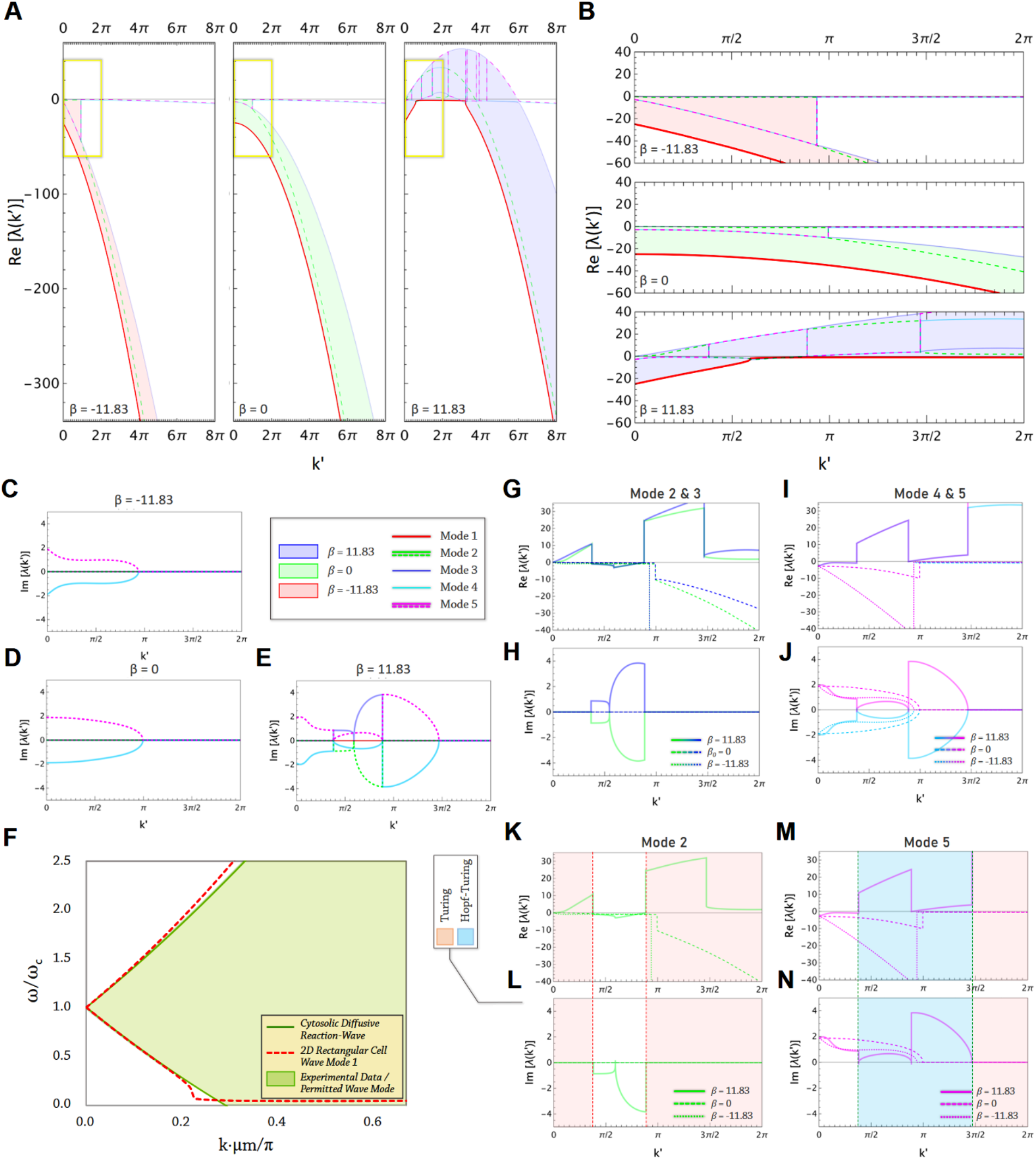
The dispersion-mode structure of bacterial Min-protein oscillator in 2D rectangular cells skewed by intracellular E-field-driven transport. (**A**) The complete diagram for the real parts of dispersion mode structures and envelops with β = 11.83, 0 and −11.83 (light blue, green and red region) respectively. (**B**) The zoom-in diagram for the real parts of dispersion mode structures and envelops correspond to the yellow boxed region in (**A**). The x-y enlarged scales are not equal. (**C-E**) The diagrams for the imaginary part of dispersion mode structures with β = 11.83 (**C**), 0 (**D**) and −11.83 (**E**) respectively. (**F**) The convergence of Mode 1 with the cytosolic reaction wave dispersion indicates that Mode 1 arises from diffusion-coupled nucleotide exchange in the cytosol and biased by E-field-driven transport. (**G-J**) The dispersion diagrams for Wave Mode 2&3 (**G-H**) and Wave Mode 4&5 (**I-J**). The real parts for all modes are in the upper panel (**G** and **I**) and the imaginary parts for all modes are in the lower panels (**H** and **J**). (**K-N**) Indications of wavenumber bands for Turing mode (Re[λ(k’)] > 0 and Im[λ(k’)] = 0) and Hopf-Turing mode (Re[λ(k’)] > 0 and Im[λ(k’)] ≠ 0) in Mode 2 and 5. (**K** and **M**) Real parts; (**L** and **N**) Imaginary parts. The color codes for Mode 1-5 are listed in the legend panel.

Unlike the other wave modes, Mode 1 (red curves in Fig. 3A, B & F) decays rather than grows, asymptotically approaching the previously approximated dispersion branches *1 ± μ(E^⇀^ • k^⇀^) + Δk^2^* of the cytosolic reaction wave (i.e., neglecting membrane-associated reactions) from the negative Re[λ(k′) side (Fig. 3F). The convergence of Mode 1 with the cytosolic reaction wave dispersion indicates that Mode 1 behaves as a decaying diffusive mode (Re[λ(k′)] < 0 and Im[λ(k′)] = 0), arising from diffusion-coupled nucleotide exchange in the cytosol and biased by E-field-driven transport (Fig. 3B), which shifts and reshapes the spatial reaction gradient. Deviations from this behavior become apparent as the wavenumber approaches the intermediate-*k* regime (*k*^′^ ≈ 3*π*⁄8; Fig. 3F). For small wavenumbers (*k*^′^ ≲ 3*π*⁄8, corresponding to *L* ≳ 10.667 *μm*), the diffusive mode typically dominates, governing dynamics at large spatiotemporal scales and remaining effectively decoupled from membrane-associated reactions. In this regime, the cytosolic reaction–wave model provides an accurate description of the system dynamics. Although classical Turing instabilities select finite wavelengths, geometric confinement creates reflective boundaries, and intracellular E-field– driven asymmetric transport enhances cytosol–membrane coupling in opposite directions as Modes 2 and 3 (denoted by Modes 2/3 hereafter) propagate along the cell. This establishes distinct reaction centers set by the diffusive length scale (≈ 8 μm, estimated by *D_MinD_* ≈ 16 μm^2^/s (*30*)), producing Turing modes with direction-dependent growth rates in cells significantly longer than this scale (green and blue curves in Fig. 3G–H).

Within the intermediate-*k* regime closer to small *k* (3*π*⁄8 ≲ *k*^′^ ≲ 8*π*⁄9, corresponding to 4.5 *μm* ≲ *L* ≲ 10.667 *μm*), cell lengths are comparable to the diffusive length scale, allowing MinE to reach the membrane in time. Cytosol–membrane coupling then suppresses diffusive and Turing modes, letting Hopf–Turing modes (Modes 4/5; Fig. 3I–J) dominate. These modes have identical growth rates and correspond to counter-propagating traveling waves whose superposition produces standing-wave patterns. At these spatial scales, set by the interplay between the diffusive Mode 1 and cytosol–membrane coupling, the diffusive length scale determines the timescale over which the inhibitor MinE reaches the membrane through its recruitment by membrane-associated MinD. As a result, MinE arrives while MinD is still accumulating, before a Turing pattern can fully develop. This timing enables MinDE complexes to assemble and dissociate on the membrane, providing the oscillatory feedback that drives a Hopf–Turing instability, which governs pattern formation in this regime.

In the higher intermediate-*k* regime (8*π*⁄9 ≲ *k*^′^ ≲ 22*π*⁄15, corresponding to 2.727 *μm* ≲ *L* ≲ 4.5 *μm*), which lies within the typical range of unit cell lengths, geometric confinement plays a dominant role. Boundary reflection effectively folds each cell half into the diffusive length scale, thereby enhancing cytosol–membrane coupling and enabling a large growth rate of the Turing instability in this regime (Modes 2/3; Fig. G-H). Under these conditions, a Turing pattern can fully develop within a cell half before sufficient MinE inhibitor accumulates at the membrane, owing to the relatively slow cytosolic diffusion of MinE. As a result, the Hopf– Turing instability (Modes 4/5; Fig. 3I–J) initially emerges at later times and competes weakly with the established Turing pattern. However, continued MinE accumulation on the membrane progressively destabilizes the Turing pattern, allowing the Hopf–Turing mode to eventually outcompete it and drive pattern disassembly. Notably, the growth rates of Modes 2 and 3 differ slightly, whereas those of Modes 4 and 5 are identical (Fig. 3G & I), implying asynchronous mode competition between cell halves and resulting in quasi-periodic dynamics. These oscillations arise from competition between stationary Turing patterns and oscillatory Hopf– Turing modes acting on comparable but non-commensurate timescales, preventing convergence to a single periodic state. Instead, geometric confinement and intracellular E-field–driven asymmetric transport sustain quasi-periodicity, imparting robustness to variations in cell size, geometry, and protein abundance during growth and division.

Finally, for cell lengths much shorter than the diffusive length scale (*k*^′^ > 22*π*⁄15 corresponding to *L* < 2.727 *μm*), Turing modes dominate, as the confined space enhances cytosol–membrane coupling, producing large direction-dependent growth rates. With MinE inhibitors distributed throughout the cell, this can trigger abrupt, collective disassembly of the Turing pattern, leading to stochastic, oscillation-like switching of spatial patterns (*31*).

### The dispersion-mode structure of bacterial Min-protein oscillator skewed by intracellular E-field-driven transport

Intracellular E-field–driven asymmetric transport separates the length scales of cytosol– membrane coupling and diffusion-coupled reaction waves, yielding skewed real parts of the dispersion modes and their envelopes (Fig. 3A–B; β = 0, β < 0, β > 0). For β ≤ 0, all dispersion branches remain negative, whereas for β > 0 some become positive in selective wavenumber bands, particularly in the intermediate-*k* regime where length scales favor cytosol–membrane coupling. When β > 0, corresponding to negatively charged Min-protein waves propagating parallel to intracellular E-field, the decay rate of the diffusive Mode 1 decreases and approaches zero over the diffusive length scale due to transport-mediated balance between nucleotide exchange and diffusion. This reduced decay enhances cytosol–membrane coupling and facilitates mode crossings near *k*^′^ ≃ 8*π*⁄9, allowing the Mode 4/5 branches to follow a dispersive trajectory that is well approximated by cytosolic reaction wave dynamics, *ω/ω_c_ ⋍ 1 + μ(E^⇀^ • k^⇀^) − Δk^2^*, in the higher intermediate-*k* regime. From this relation, the optimal wavenumber *k_max_* that maximizes the growth rate of Modes 4/5 is obtained by *бRe*(*ω*)⁄*бk* = 0, yielding *k_max_ = μ(E^⇀^•k^⇀^)−Δk^2^*, where k^ is the unit wavevector. Using our PNP-based electrochemical model of bacterial electrolytes, we simulated intracellular E-fields across a range of cell lengths (Fig. 2B). The resulting mean E-field strengths are consistent with the theoretical values required to select *k_max_*, or cell length, supporting maximal growth of Modes 4/5, corroborating the essential role of intracellular E-fields in regulating the bacterial Min-protein oscillator.

In the higher intermediate-*k* regime (around *k*′ ≃ 8*π*⁄9), mode crossings align the Mode 4/5 branches with a dispersive trajectory approximated by cytosolic reaction wave dynamics (comparing the branches in the β > 0 panel in Fig. 3A with the lower branch of Fig. 3F from the negative Re[λ(*k’*)] side), allowing the definition of a wave growth-rate velocity *v_G_ = Re(ω)/k≃−μ(E^⇀^•k^⇀^)−Δk^2^*. This velocity encapsulates the outcome of mode competition among the diffusive mode, stationary Turing modes, and oscillatory Hopf–Turing modes, reflecting the balance between reaction-driven growth or decay, E-field–induced transport, and diffusive damping. Notably, Hopf-Turing mode would eventually outcompete Turing mode because the former intrinsically destabilize the later mode, here, by MinE’s inhibition upon the accumulation of membrane-bound MinD. Consequently, *v_G_* determines whether Mode 4/5 patterns locally amplify (*v_G_* > 0) or regress (*v_G_* < 0) when these modes interact on comparable spatiotemporal scales. Theoretical *v_G_* values correspond to negatively charged Min-protein waves propagating parallel or antiparallel to the intracellular E-field (*k^⇀^* parallel or antiparallel to *μ(E^⇀^)*; Fig. 2C-F). Wave growth-rate velocities computed from simulated E-field distributions across various cell lengths (Fig. 2B) agree with these theoretical predictions. Waves propagating antiparallel to the E-field exhibit *v_G_* < 0, reflecting retrograde patterns in which the Hopf–Turing instability and subsequent diffusive cytosolic reaction waves shift toward the opposite cell half (Fig. 2E). In contrast, waves propagating parallel to the E-field when 1⁄*L* ≧ 0.238 *μm*^-1^ show *v_G_* > 0, indicating pattern growth driven by the cytosol-membrane coupling between the diffusive cytosolic reaction wave and the Turing instability within the same cell half (Fig. 2F).

Intracellular E-field–driven asymmetric transport skews the dispersion-mode structure of the bacterial Min-protein oscillator by separating the length scales of cytosol–membrane coupling and diffusion-coupled reaction wave. In the higher intermediate-*k* regime, this promotes mode crossings between diffusive, Turing, and Hopf–Turing instabilities, with the wave growth-rate velocity *v_G_* determining local pattern amplification or regression depending on wave propagation relative to the E-field. Simulations of intracellular E-fields across cell lengths agree with theoretical predictions, highlighting intracellular E-field as a key regulator of mode competition and spatial pattern formation in bacterial Min-protein dynamics.

### The frequency modulations of bacterial Min-protein oscillator via experimental manipulations of nucleoid morphology

As implicated by numerical simulations of intracellular E-fields (Fig. S11), anucleate and nucleoid-perturbed bacteria can be used to alter nucleoid morphology (*27, 32, 33*), thereby modulating charge density and E-field strength to test wave-field interactions in bacterial Min-protein oscillator. Prior experiments (*34*) report frequency modulations in anucleate strains (SH6761, Fig. 4A) and antibiotic-treated wildtype cells (RP437, Fig. 4B). Kymographs of Min-protein dynamics under different nucleoid manipulations (Fig. 4C–F) reveal distinct spatiotemporal patterns. Half-cycle averages of normalized longitudinal mCherry-MinD intensity profile show steep polar gradients in chloramphenicol-or rifampicin-treated cells relative to wildtype (Fig. 4D–E), and moderate gradients in anucleate cells (Fig. 4F). Compressive or expanded nucleoids reduce or increase mean oscillation frequency and narrow or broaden frequency distributions, respectively (Fig. 4G–I).

**Fig. 4.**
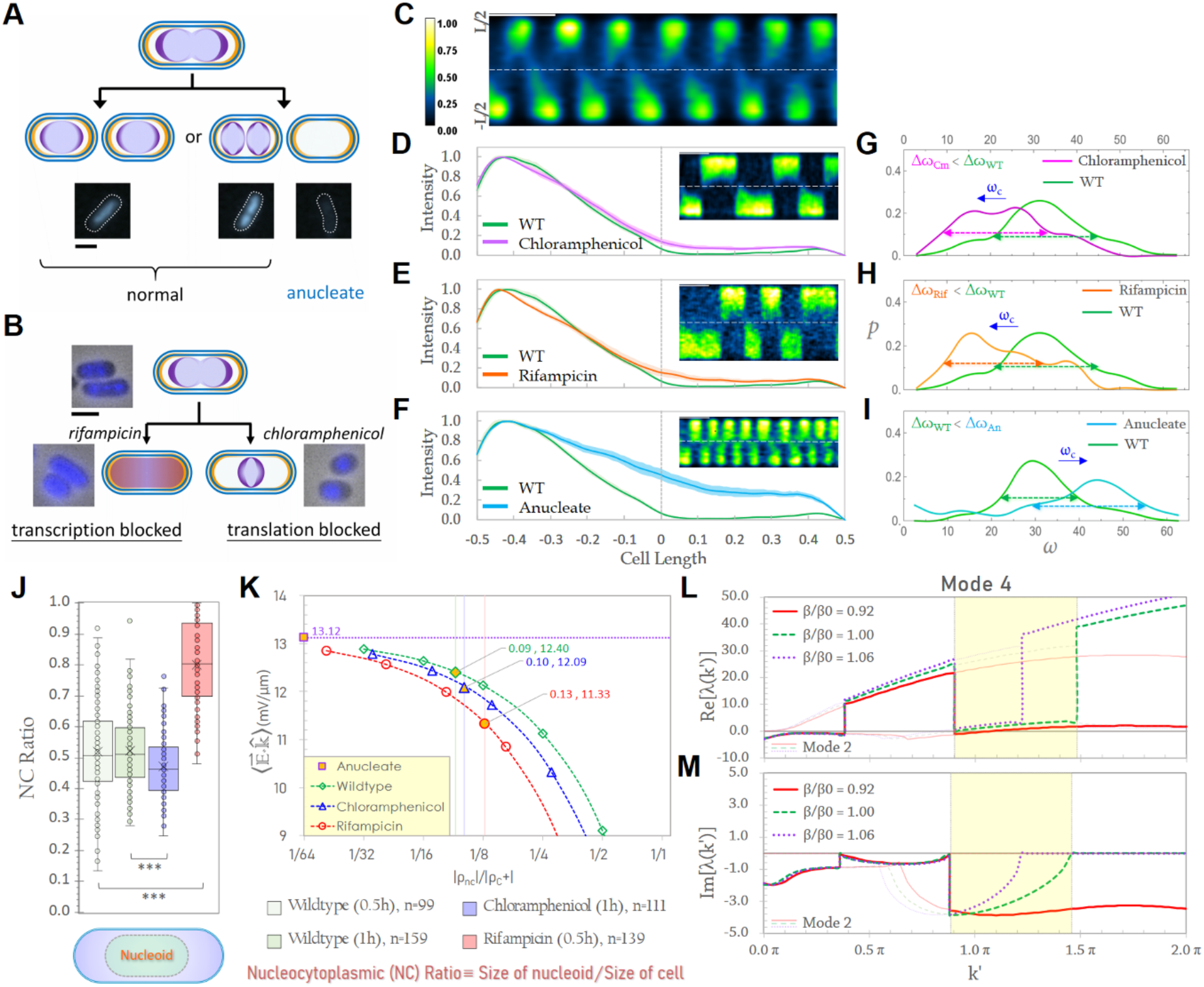
The frequency modulations of Min-protein oscillator in nucleoid-perturbed bacteria. (**A**) The concept of the nucleoid segregation in the *ΔmukB* strain, SH6067. (**B**) The concept of the nucleoid perturbation by rifampicin and chloramphenicol. The merge insets are phase-contrast images of cells (grey) and the fluorescence images of the DAPI-stained nucleoid (blue). Scale bar in (**A-B**): 1 μm. (**C**) The kymograph of MinD oscillations in wildtype bacteria. Vertical color bar: normalized fluorescence intensity. Scale bars in (**C-F**): 60 sec. Other kymographs (**D-F**) drawn to the same scale as (**C**). Longitudinal, normalized intensity profile over half-cycle average for (**D**) chloramphenicol-treated (magenta), (**E**) rifampicin-treated (orange) and (**F**) anucleate (cyan) bacteria in comparison with that of wildtype bacteria (green). The band around profile curves indicate confidence interval (n=10). Cell length is normalized to unit cell length. (**G-I**) The frequency distribution of MinD oscillations for (**G**) chloramphenicol-treated (magenta), (**H**) rifampicin-treated (orange) and (**I**) anucleate (cyan) bacteria in comparison with that of wildtype bacteria (green). The shift of mean oscillation frequency ωc for anucleate and nucleoid-perturbed bacteria in comparison with that for wildtype bacteria is labeled by blue arrow in (**G-I**). (**J**) The distributions of NC ratios for wildtype, chloramphenicol and rifampicin-treated bacteria. (**K**) The dependence of mean E- field strength upon the ratio of charge density |ρ_nc_⁄ρ_C_+| for wildtype, chloramphenicol and rifampicin-treated and anucleate bacteria. The filled diamond, triangle and circle symbols denote the mean E-field strength corresponding to the charge density ratio estimated in wildtype, chloramphenicol and rifampicin-treated bacteria. The filled square symbol denotes the mean E-field strength for anucleate bacteria. (**L-M**) The real (**L**) and imaginary (**M**) parts of Dispersion branches for Mode 2 and 4 with β/β0 = 1.06, 1.00 and 0.92, each corresponding to anucleate, wildtype and antibiotic-treated cells respectively. β0 = 11.83 (see Section D in SI). [Images in **A**-**F** are adopted from (*34*)]

To probe whether wave-field interactions underlie these frequency modulations, we simulated E-field changes induced by altered nucleoid charge densities, specifying distinct nucleoid morphologies for wildtype, chloramphenicol-treated, rifampicin-treated, and anucleate cells (Fig. 4J–K, S14). Total cellular charge is kept constant, while the ratio of nucleoid to cytosolic charge density, |*ρ_nc_*|⁄|*ρ_C_*+ |, ranged from 2^2^ to 2^-6^ to reflect morphological changes. Wildtype cells correspond to NC ratio ≈ 0.52 (*26*), while anucleate cells lack the nucleoid, leading to zero NC ratio (Fig. S14). Chloramphenicol-treated cells, with compressed nucleoids, have NC ratio ≈ 0.47; despite higher apparent charge density, residual synthesis of nucleoid associated proteins increases total nucleoid charge ≈ 1.17-fold adopted in our simulations. Rifampicin-treated cells, with expanded nucleoids, have NC ratio ≈ 0.8; although charge density decreases, the total nucleoid charge increases 4 – 5-fold due to inclusion of additional DNA, RNA, and proteins, yielding an effective 2.5-fold increase for evaluating E-field strength in simulations (see Materials and Methods and Appendix, SI for estimation of nucleoid charge density).

The mean E-field strength in each cell state is plotted versus the charge-density ratio |*ρ_nc_*|⁄|*ρ_C_*+| (Fig. 4K). In anucleate cells, the mean field at zero nucleoid charge is ∼13.12 mV/μm (filled square). Across the relevant charge-density range *(|ρ_nc_|/|ρ_C+_|≈1/32 − 1/4)*, the mean E-field is highest in anucleate cells, intermediate in wildtype (green diamond), and lowest in chloramphenicol-(blue triangle) and rifampicin-treated cells (red circle). Based on *E. coli* nucleoid composition (*35, 36*) in a 3.636 µm long cell, |*q_nc_*|⁄|*q_C_*+| ≈ 1/30 converts to |*ρ_nc_*|⁄|*ρ_C_*+| ≈1/11 for wildtype, 1/10 for chloramphenicol-treated, and 1/8 for rifampicin-treated cells. Statistical analyses of E-field distributions (Fig. S15) confirm that mean field strengths decrease in the order: anucleate > wildtype > chloramphenicol-treated > rifampicin-treated cells. Because the region of permitted wave modes in *ω* − *k* space (Fig. 2A) depends on mean E-field strength, both the size of the permitted region and the mean frequency and the width of the Min-protein oscillation distributions scale accordingly (Fig. 4G–I).

Beyond the experimental and numerical verification of altered mean intracellular E-fields arising from perturbed nucleoid morphologies and associated charge densities, further physical insight can be obtained from dispersion-based analyses of the reaction–diffusion–transport model in 2D rectangular cells. Guided by experimental estimates (Fig. 4J) and simulations (Fig. 4K), mean E-field strengths corresponding to anucleate, wildtype, and antibiotic-treated cells—parameterized as β/β₀ = 1.06, 1.00, and 0.92, respectively—were incorporated into the dispersion-based analysis, revealing pronounced differences in the dispersion of Modes 4/5 within the higher intermediate-*k* regime (yellow regions in Fig. 4L–M). For antibiotic-treated cells (β/β₀ = 0.92), reduced E-field–driven transport weakens cytosol–membrane coupling and delays MinE recruitment at these spatiotemporal scales, resulting in substantially lower growth rates of Hopf–Turing mode (red curve in Fig. 4L). This favors prolonged, asymmetric Turing patterns (light red curve in Fig. 4L) in each cell half, consistent with the unequal growth rates of Modes 2/3 under asymmetric transport (Fig. 3G). The group velocity of Modes 4/5, *v*_g_ = ∂(Im(λ(*k’*)))/∂*k’*, is smaller in magnitude for antibiotic-treated cells (the derivative of red curve in Fig. 4L), suggesting slower oscillations (cf. kymographs in Fig. 4C-E). Moreover, the phase velocity of Modes 4/5, *v_p_* = Im(λ(*k’*))/*k’*, is larger in magnitude and negative for antibiotic-treated cells (Fig. 4M), indicating a fast retrograde pattern propagation driven by the Hopf– Turing mode (cf. kymographs in Fig. 4C-E).

In contrast, anucleate cells (β/β₀ = 1.06) exhibit stronger E-field–driven transport, yielding higher growth rates of Modes 2/3 and 4/5, and slightly faster oscillations (larger *v*_g_) than wildtype cells (purple dotted curve in Fig. 4L). For shorter cells (*k*^′^ ≳ 6*π*⁄5 or *L* ≲ 3.333 *μm*), Modes 4/5 transition to Turing modes with large, asymmetric growth rates due to geometric confinement and associated enhanced MinE availability (Fig. S16), leading to rapid, stochastic switching of Turing patterns (Fig. 4F) (34). These dispersion features closely mirror the experimentally observed pattern dynamics (Fig. 4D–F), further validating the role of wave– field interactions in shaping Min-protein oscillations.

Together, these experimental, numerical, and theoretical analyses demonstrate that nucleoid morphology modulates the bacterial Min-protein oscillator by tuning intracellular E-field strength through changes in charge density. Perturbations to the nucleoid shift the region of permitted wave modes and selectively alter the growth and propagation of the Hopf–Turing and Turing instabilities, thereby controlling oscillation frequency, directionality, and pattern stability. The quantitative agreement between dispersion-based predictions and experimentally observed spatiotemporal dynamics supports a unified wave–field interaction framework, in which intracellular E-field acts as a key physical regulator linking cell-scale electrostatics to robust yet adaptable Min-protein patterning.

## Discussions

Here we propose that the extension of the diffuse layer is a critical but underappreciated feature of micron-scale biological systems. In confined cellular geometries, unequal effectiveness of cationic and anionic charges gives rise to an extended diffuse layer reaching several hundred nanometers, with the S/V ratio and nucleoid organization serving as key determinants of the resulting electrochemical (ECM) potential (Fig. 1). The emergence of a sustained intracellular E-field motivates a combined theoretical, numerical, and experimental investigation of its coupling to the bacterial Min-protein oscillator. Our results provide convergent evidence for chemical wave–electric field interactions in bacteria, manifested experimentally through both pattern alterations and frequency modulations.

Wave–field interactions affect Min-protein dynamics in two distinct but coupled mechanisms: modulation of reaction gradients in real space and alteration of wave dispersion landscape in phase space. First, the altered reaction gradients arise from direct E-field-driven transport of Min proteins (Fig. 5A.1). Under a higher E-field, the field-induced drift flux of negatively charged Min proteins at the polar zone opposes the diffusive flux from the opposite pole, promoting substantial depletion of Min proteins from the polar region and thereby weakening the local concentration gradient. Conversely, a lower E-field reduces drift, allowing Min proteins to accumulate at the polar zone and generating a more pronounced concentration gradient. Because the membrane has a finite capacity for Min-protein binding, surface reaction rates are additionally coupled to the cytosolic Min-protein concentration, resulting in heterogeneous, fractional-order reaction kinetics characteristic of Langmuir-Hinshelwood-type surface reactions (*37*) (see Supplementary Information). For example, membrane binding that is first-order when binding sites are abundant can approach zero-order kinetics when the surface becomes saturated. This transition also alters the characteristic timescale of reactant consumption: for first-order kinetics, the half-life is concentration-independent, whereas for zero-order kinetics it increases with reactant concentration. In the Min-protein system, surface reactions include not only membrane binding but also autocatalytic reactions involving membrane-bound Min proteins, for which the reaction timescale similarly increases with reactant abundance. Thus, E-field-driven transport modulates both the spatial distribution of Min proteins and the kinetics of the reactions that sustain their dynamics. Consequently, higher E-fields accelerate the oscillatory dynamics, yielding a shorter oscillation period. These findings reveal a real-space instability of the Min-protein wave, in which changes in E-field strength reshape both the reaction gradients and the underlying reaction kinetics.

**Fig. 5.**
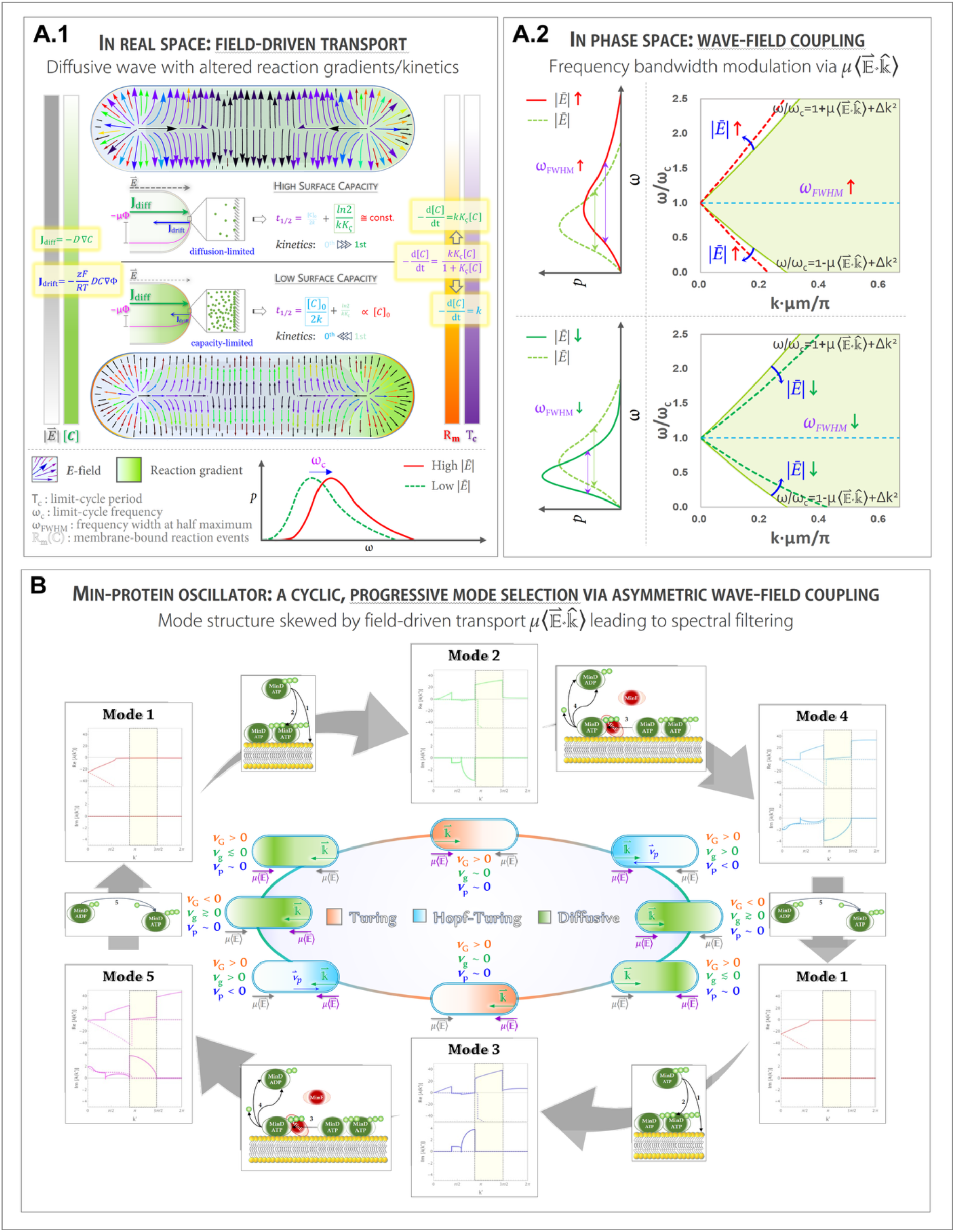
The instabilities of bacterial Min-protein oscillator due to chemical wave – electric field interactions. (**A.1**) E-field-driven transport reshapes polar Min-protein concentration gradients and membrane reaction kinetics R_m_(C) in a field-strength- dependent manner *(|E^⇀^|)*. Direct membrane binding illustrates the transition from first- order kinetics under weak concentration gradients and unsaturated binding sites (lower panel) to zero-order kinetics under pronounced polar accumulation and membrane saturation (upper panel), extending the reaction timescale and modulating the oscillation period T_c_ and frequency ω_c_. The intracellular E-field structure and corresponding Min-protein concentration gradients under high and low fields are shown at the top and bottom of the upper and lower panels, respectively. Diffusive and drift fluxes (J_diff_ and J_drift_) are given at left; *D*, *z*, *F*, *R*, *T*, *μ* and Φ denote the diffusion coefficient, valence, Faraday constant, gas constant, temperature, electric mobility and electric potential, respectively. Reaction-rate equations and their kinetic transitions are shown at right, where *k*, *K_ς_*, and t_1/2_ denote the rate constant, surface-adsorption equilibrium constant, and half-time for consumption of the initial reactant concentration [*C*]_0_, respectively. (**A.2**) E-field strength alters wave-phase coupling and dispersion: lower fields promote polar accumulation and membrane saturation, narrowing the permitted frequency bandwidth, whereas higher fields expand the accessible wave-mode and frequency ranges. Thus, the presence and strength of the intracellular E-field provide an electric cue that couples Min-protein transport, reaction kinetics, and wave propagation. (**B**) Cyclic mode selection underlying the instability in phase space mediated by asymmetric wave–field coupling. Asymmetric E-field–driven transport separates the length scales of cytosol–membrane coupling and diffusion-coupled reaction waves, skewing the dispersion mode structure into an effective spectral filter for mode selection, with spatiotemporal scales set by the diffusive length scale and cell length. For bacterial Min-protein oscillator with typical cell length, the oscillation cycle begins with Mode 1, a diffusion-coupled reaction wave propagating from one cell half to the other, which locally enhances cytosol–membrane coupling and promotes MinD membrane association (Steps 1– 2). This triggers a Turing instability (Mode 2 or 3) that grows until saturation. Owing to the delayed arrival of MinE, set by its diffusion spatiotemporal scales, MinE is subsequently recruited to the membrane to form MinDE complexes (Step 3), leading to MinD/MinE dissociation (Step 4) and destabilization of the Turing pattern, thereby activating a Hopf– Turing instability (Mode 4 or 5). Released MinD re-enters the cytosol, undergoes nucleotide exchanges during diffusion/transport (Step 5), and reforms a reaction wave that propagates into the opposite cell half, where the sequence repeats. This cyclic, progressive mode selection coordinates diffusive, Turing, and Hopf–Turing modes to sustain robust Min-protein oscillations. *μ(E^⇀^)* in purple color denotes the intracellular E-field that mediates wave–field\ interactions. *k^⇀^* is the wavevector. Here, *vG* represents the growth-rate velocity of wave mode, *vg* the group velocity and *vp* the phase velocity.

Second, E-field perturbations also act in phase space, reinforcing another instability in wave dynamics through spatially asymmetric coupling between the E-field and wave propagation (Fig. 5A.2). Because the Min-protein wavelength is comparable to the intracellular E-field gradient, waves longitudinally traversing an asymmetric field along the cell (Fig. 1M and Fig. S11) experience position-dependent coupling angles, leading to heterogeneous phase responses along the wave. As a consequence, an outwardly directed E-field (Fig. S11) biases negatively charged Min proteins toward the poles when field strength decreases due to increased nucleoid charge density (Fig. 4D–E, J–K). The resulting polar accumulation drives membrane saturation, which limits membrane-associated reaction rates, extends the reaction timescale, prolongs the oscillation period, and narrows the permitted frequency band (lower panel, Fig. 5A.2). Conversely, increasing E-field strength by reducing nucleoid charge density drives Min proteins toward the nucleoid, yielding more moderate polar gradients (Fig. 4F, J, K) and expanding both the permitted wave-mode region and frequency bandwidth (upper panel, Fig. 5A.2). The presence and strength of the intracellular E-field act as an electric cue that couples Min-protein transport, reaction kinetics, and wave propagation, thereby tuning the stability and frequency of dynamic protein patterns.

From a theoretical perspective, the dispersion-mode structure skewed by intracellular E-field– driven transport reveals a physically novel mechanism for regulating Min-protein oscillations. Asymmetric transport orchestrates cyclic and progressive mode selection, facilitating a second instability associated with cytosol–membrane coupling and diffusion-coupled reaction waves across distinct spatiotemporal scales. This process effectively implements spectral filtering through mode competition and transition (Fig. 5B). A complete oscillation cycle therefore requires a coordinated succession of cytosolic diffusive reaction waves, stationary Turing modes, and oscillatory Hopf–Turing modes (Fig. 3G–N), each linked to specific biochemical processes (Fig. 5B). Such a hierarchy enables functional robustness and flexibility in the noisy intracellular environment, allowing the Min-protein system to adapt to changes in cell size, geometry, and protein abundance during growth and division. Maintaining the system in a quasi-periodic regime—achieved here through asymmetric mode competition mediated by wave–field interactions—appears to be a general strategy for reliable intracellular oscillations.

Finally, chemical wave–electric field interactions are unlikely to be unique to bacteria. In eukaryotic cells, pronounced morphological changes can locally reduce cellular dimensions to a few hundred nanometers, amplifying electrostatic and transport effects. For example, reversible switching between fast and slow keratocyte migration is associated with large variations in intracellular diffusion and lamellipodial thickness (76–556 nm), driven by small GTPase–mediated actomyosin dynamics (*38*). These observations suggest that intracellular E-fields, wave–field coupling, and spatial organization of charged biomolecules may play broader regulatory roles across diverse biological systems, warranting further investigation.

## Data availability

Simulation data and raw images are available from the corresponding author upon request.

## Code availability

Custom-built Mathematica and MATLAB scripts are available from the corresponding author upon request.

## Contributions

JPS designed/conducted experiments, analyzed data/images, conducted theoretical analysis, built the mathematical models and wrote the codes, conducted numerical simulations, wrote the manuscript. CFC acquired the funding, supervised the project, and co-wrote the manuscript.

## Acknowledgment

We thank Prof. Yi-Ren Chang for his help on experiments for Figure S13. We appreciate Prof. Victor Sourjik for insightful discussions and kindly sharing experimental data with us. We also thank helpful discussions with Profs. Robert Austin, Erwin Frey, and Tzyy-Leng Horng, and Drs. Hong-Yan Shih, Wei-Shiang Lin, Keita Kamino and Tetsuya Hiraiwa. We acknowledge funding support from National Science and Technology Council (grants # NSTC 111-2112-M-001-029-MY3 and 114-2112-M-001-036-MY3) and Academia Sinica (AS-IA-109-M04 and AS-iMATE-115-12). We thank DNA Sequencing Core Facility of the Institute of Biomedical Sciences, Academia Sinica for DNA sequencing analysis. The core facility is funded by Academia Sinica Core Facility and Innovative Instrument Project (AS-CFII-113-A12). This work was written in part at Aspen Center for Physics, which is supported by National Science Foundation grant PHY-2210452.

## References

1. A. N. Zaikin, A. M. Zhabotinsky, Concentration wave propagation in two-dimensional liquid-phase self-oscillating system. Nature 225, 535–537 (1970).

2. A. M. Turing, The Chemical Basis of Morphogenesis. Philos. Trans. R. Soc. Lond. B Biol. Sci. 237, 37–72 (1952).

3. L. Wolpert, Positional information and the spatial pattern of cellular differentiation. J. Theor. Biol. 25, 1–47 (1969).

4. D. M. Raskin, P. A. de Boer, MinDE-dependent pole-to-pole oscillation of division inhibitor MinC in Escherichia coli. J. Bacteriol. 181, 6419–6424 (1999).

5. K. C. Huang, Y. Meir, N. S. Wingreen, Dynamic structures in Escherichia coli: spontaneous formation of MinE rings and MinD polar zones. Proc Natl Acad Sci U S A 100, 12724–12728 (2003).

6. S. Schmidt, P. Ortoleva, A new chemical wave equation for ionic systems. J. Chem. Phys. 67, 3771–3776 (1977).

7. S. Schmidt, P. Ortoleva, Electric field effects on propagating BZ waves: Predictions of an Oregonator and new pulse supporting models. J. Chem. Phys. 74, 4488–4500 (1981).

8. P. Ortoleva, Chemical wave-electrical field interaction phenomena. Physica D 26, 67–84 (1987).

9. P. Ortoleva, S. L. Schmidt, in Oscillations and Traveling Waves in Chemical Systems, R. J. Field, M. Burger, Eds. (1985), chap. 10, pp. 333–418.

10. R. Feeney, S. L. Schmidt, P. Ortoleva, Experiments on electric field-BZ chemical wave interactions: Annihilation and the crescent wave. Physica D 2, 536–544 (1981).

11. H. Ševčíková, M. Marek, Chemical front waves in an electric field. Physica D 13, 379–386 (1984).

12. H. Strahl, L. W. Hamoen, Membrane potential is important for bacterial cell division. Proc Natl Acad Sci U S A 107, 12281–12286 (2010).

13. L. P. Savtchenko, M. M. Poo, D. A. Rusakov, Electrodiffusion phenomena in neuroscience: a neglected companion. Nat. Rev. Neurosci. 18, 598–612 (2017).

14. E. J. F. Dickinson, J. G. Limon-Petersen, R. G. Compton, The electroneutrality approximation in electrochemistry. J. Solid State Electrochem. 15, 1335–1345 (2011).

15. K. R. Ward, E. J. F. Dickinson, R. G. Compton, How Far Do Membrane Potentials Extend in Space Beyond the Membrane Itself? Int. J. Electrochem. Sci. 5, 1527–1534 (2010).

16. M. Marhl, M. Brumen, R. Glaser, R. Heinrich, Diffusion layer caused by local ionic transmembrane fluxes. Pflugers Arch 431, R259–260 (1996).

17. E. J. Dickinson, L. Freitag, R. G. Compton, Dynamic theory of liquid junction potentials. J. Phys. Chem. B 114, 187–197 (2010).

18. B. Eisenberg, Interacting ions in biophysics: real is not ideal. Biophys. J. 104, 1849–1866 (2013).

19. J. J. Jasielec, Electrodiffusion Phenomena in Neuroscience and the Nernst–Planck– Poisson Equations. Electrochem 2, 197–215 (2021).

20. S. Bakshi, H. Choi, J. C. Weisshaar, The spatial biology of transcription and translation in rapidly growing Escherichia coli. Front Microbiol 6, 636 (2015).

21. P. E. Schavemaker, W. M. Smigiel, B. Poolman, Ribosome surface properties may impose limits on the nature of the cytoplasmic proteome. Elife 6, e30084 (2017).

22. W. M. Smigiel et al., Protein diffusion in Escherichia coli cytoplasm scales with the mass of the complexes and is location dependent. Sci Adv 8, eabo5387 (2022).

23. D. Valverde-Mendez et al., Macromolecular interactions and geometrical confinement determine the 3D diffusion of ribosome-sized particles in live Escherichia coli cells. Proc Natl Acad Sci U S A 122, e2406340121 (2025).

24. H. Meinhardt, Turing’s theory of morphogenesis of 1952 and the subsequent discovery of the crucial role of local self-enhancement and long-range inhibition. Interface Focus 2, 407–416 (2012).

25. E. Gianazza, P. Giorgio Righetti, Size and charge distribution of macromolecules in living systems. J. Chromatogr. A 193, 1–8 (1980).

26. W. T. Gray et al., Nucleoid Size Scaling and Intracellular Organization of Translation across Bacteria. Cell 177, 1632–1648 e1620 (2019).

27. E. Boye, A. Lobner-Olesen, Bacterial growth control studied by flow cytometry. Res. Microbiol. 142, 131–135 (1991).

28. V. Yeong, E. G. Werth, L. M. Brown, A. C. Obermeyer, Formation of Biomolecular Condensates in Bacteria by Tuning Protein Electrostatics. ACS Cent Sci 6, 2301–2310 (2020).

29. F. Wu, B. G. van Schie, J. E. Keymer, C. Dekker, Symmetry and scale orient Min protein patterns in shaped bacterial sculptures. Nat Nanotechnol 10, 719–726 (2015).

30. G. Meacci et al., Mobility of Min-proteins in Escherichia coli measured by fluorescence correlation spectroscopy. Phys Biol 3, 255–263 (2006).

31. E. Fischer-Friedrich, G. Meacci, J. Lutkenhaus, H. Chate, K. Kruse, Intra- and intercellular fluctuations in Min-protein dynamics decrease with cell length. Proc Natl Acad Sci U S A 107, 6134–6139 (2010).

32. Q. Sun, W. Margolin, Effects of perturbing nucleoid structure on nucleoid occlusion-mediated toporegulation of FtsZ ring assembly. J. Bacteriol. 186, 3951–3959 (2004).

33. Z. Binenbaum, A. H. Parola, A. Zaritsky, I. Fishov, Transcription- and translation-dependent changes in membrane dynamics in bacteria: testing the transertion model for domain formation. Mol. Microbiol. 32, 1173–1182 (1999).

34. J. P. Shen, Y. R. Chang, C. F. Chou, Frequency modulation of the Min-protein oscillator by nucleoid-associated factors in Escherichia coli. Biochem. Biophys. Res. Commun. 525, 857–862 (2020).

35. O. G. Stonington, D. E. Pettijohn, The folded genome of Escherichia coli isolated in a protein-DNA-RNA complex. Proc Natl Acad Sci U S A 68, 6–9 (1971).

36. A. Worcel, E. Burgi, On the structure of the folded chromosome of Escherichia coli. J. Mol. Biol. 71, 127–147 (1972).

37. H. S. Fogler, Elements of Chemical Reaction Engineering. (Prentice Hall, ed. 5th, 2016).

38. C. Jiang et al., Switch of cell migration modes orchestrated by changes of three-dimensional lamellipodium structure and intracellular diffusion. Nat Commun 14, 5166 (2023).

39. D. Chen et al., Predicting Electrophoretic Mobility of Proteoforms for Large-Scale Top-Down Proteomics. Anal. Chem. 92, 3503–3507 (2020).

