## Supplementary Materials for "Evidence of Chemical Wave-Electric Field Interaction in Bacterial Cells"

Contents:

Supplementary Figures S1-S16.

A. Materials and Methods (p.21)

B. Discrepancies of Electroneutrality in Real Cellular Context (p.25)

C. Models for Intracellular Electric Potential and Chemical Wave – Electric Field Interaction (p.29)

D. Dispersion Relation for Min-protein Waves in a 2D Rectangular Cell (p.41)

E. Statistical Analysis (p.54)

F. Capacity-Limited Surface Kinetics (p.55)

Appendix: Key Numbers and Parameters (p.61)

References (p.64)

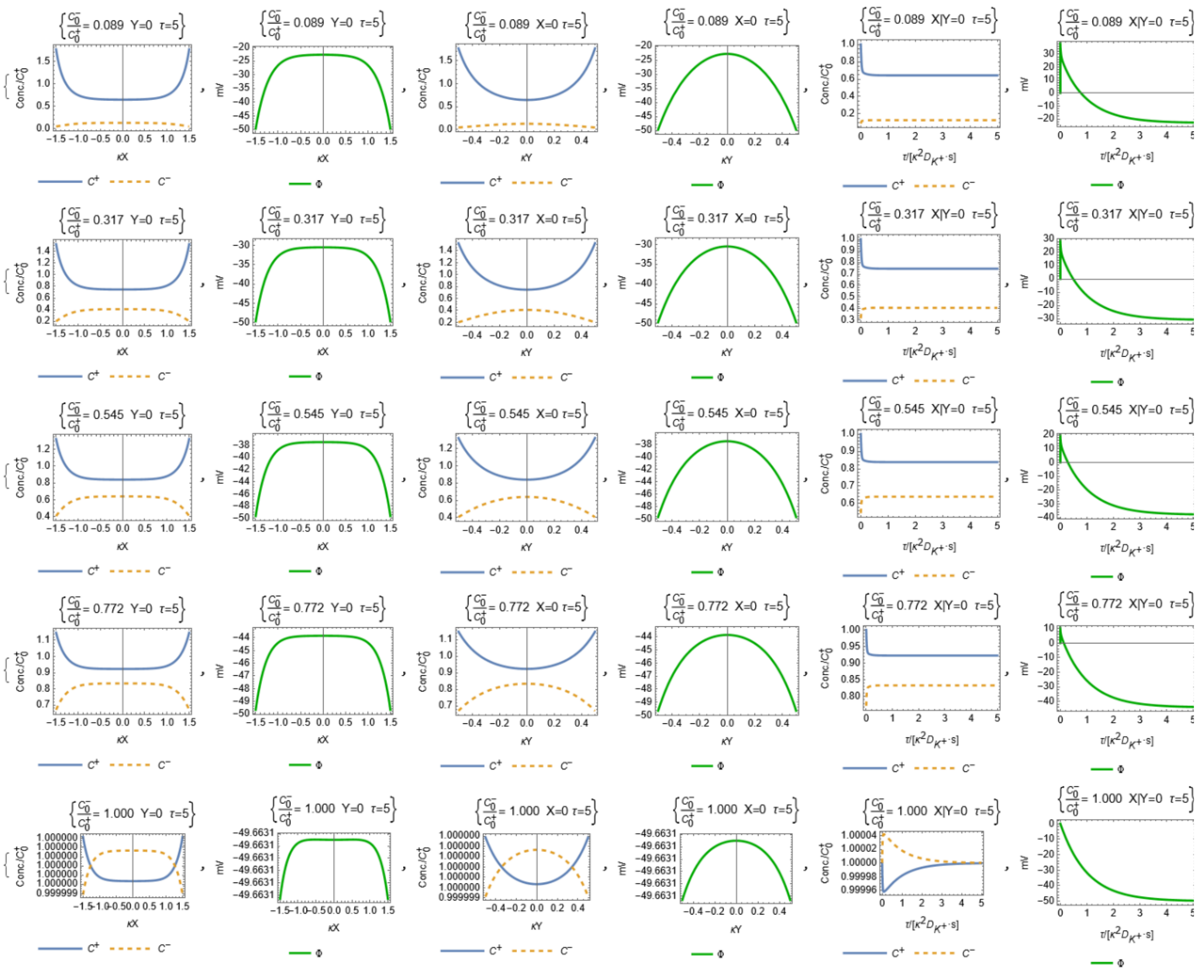


FIG. S1 The longitudinal and lateral concentration profiles [1st and 3rd column] and electrical potentials [2nd and 4th column] in the bounded, rod-shaped cell geometry [cell length/width: 3000/1000·$\kappa_D^{-1}$ nm, where $\kappa_D^{-1} \approx 0.91$ nm as the Debye length for 231.25 mM of ionic solution] are illustrated for the anion/cation ratios of 0.089, 0.317, 0545, 0.772 and 1.000 [1st to 5th row]. The time courses from τ = 1 to 5 for the longitudinal and lateral concentration profiles and electrical potential [5th and 6th column] are shown for the same range of the anion/cation ratios, where $\tau$ is the dimensionless time and relates to $t$ by $\tau = \kappa^2 D_{K^+} \cdot t$. ($\kappa$ is the inverse of Debye length $\lambda_D$ and $D_{K^+}$ is the diffusion coefficient for K$^+$). $\tau = 1$ is equivalent to $t = 0.424\, ns$. $\lambda_D$ is equal to 0.91 nm.

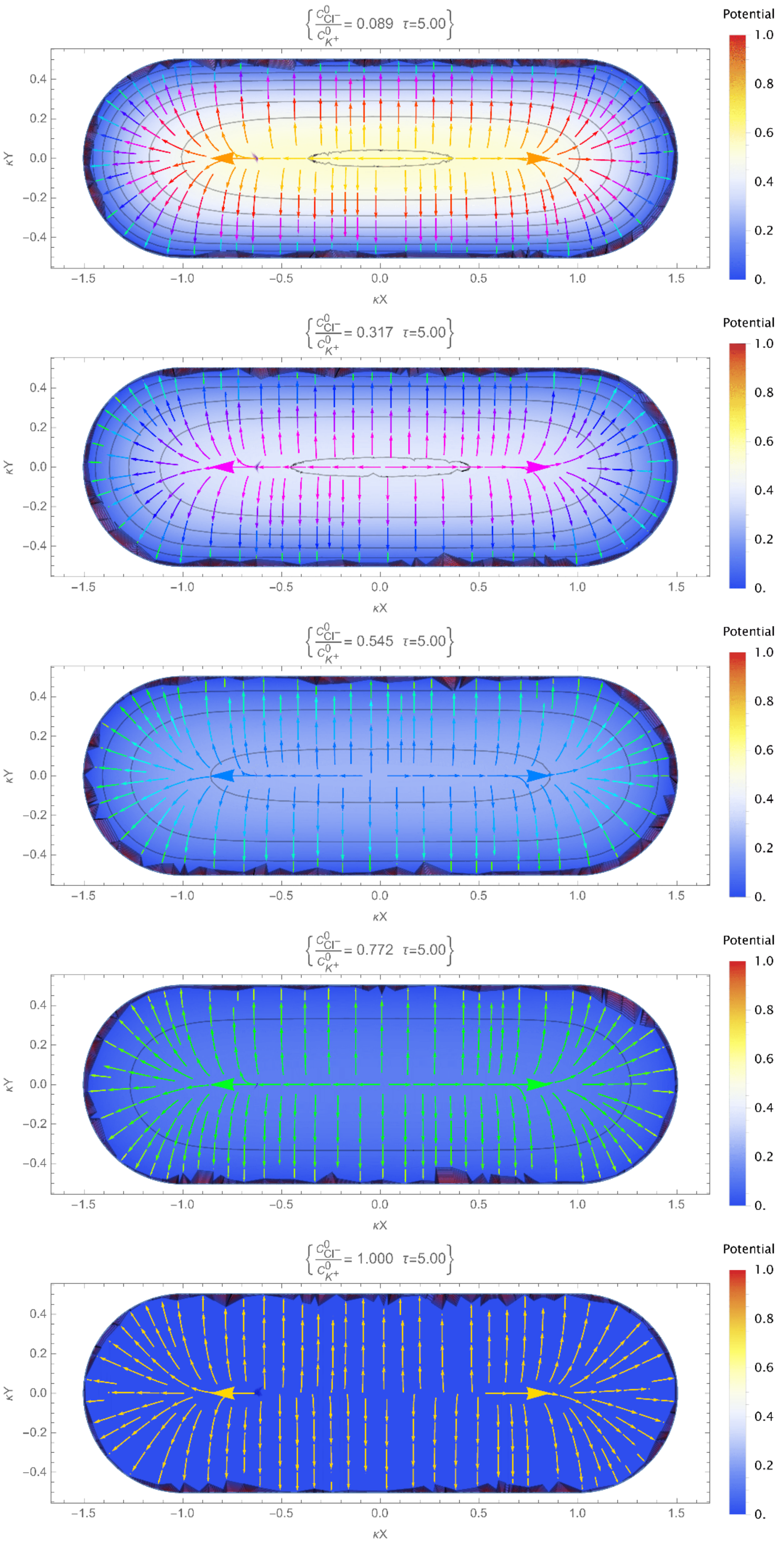

{C0Cl− / C0K+ = 0.089 τ=5.00}
{C0Cl− / C0K+ = 0.317 τ=5.00}
{C0Cl− / C0K+ = 0.545 τ=5.00}
{C0Cl− / C0K+ = 0.772 τ=5.00}
{C0Cl− / C0K+ = 1.000 τ=5.00}
Potential
1.0
0.8
0.6
0.4
0.2
0.
κY
κX
−1.5
−1.0
−0.5
0.0
0.5
1.0
1.5
0.4
0.2
0.0
−0.2
−0.4

FIG. S2 The electrical field (streamline arrows) and potential (contours) at $\tau = 5$ in the bounded, rod-shaped cell geometry [cell length/width: 3000/1000·$\kappa_D^{-1}$ nm, where $\kappa_D^{-1}$ was defined in Fig. S1] for the cation/anion ratios of 0.089, 0.317, 0545, 0.772 and 1.000 [1st to 5th row]. The potentials are normalized to the maximum and minimum levels among presented cation/anion ratios. $\kappa_D^{-1}$ is equal to 0.91 nm.

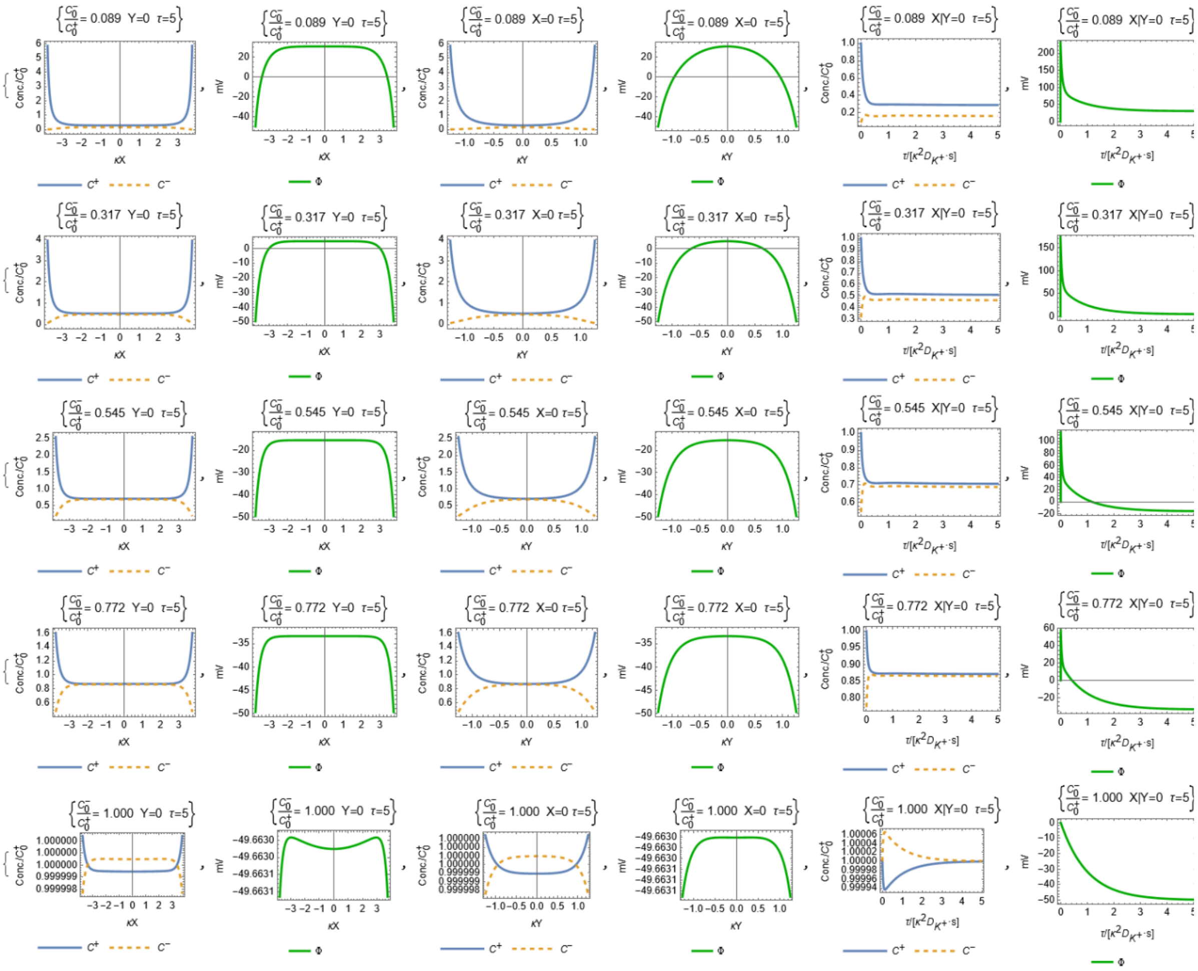


FIG. S3 The longitudinal and lateral concentration profiles [1st and 3rd column] and electrical potentials [2nd and 4th column] in the bounded, rod-shaped cell geometry [cell length/width: 7500/2500 $\cdot\kappa_D^{-1}$ nm, where $\kappa_D^{-1}$ was defined in Fig. S1] are illustrated for the anion/cation ratios of 0.089, 0.317, 0545, 0.772 and 1.000 [1st to 5th row]. The time courses from τ = 1 to 5 for the longitudinal and lateral concentration profiles and electrical potential [5th and 6th column] are shown for the same range of the anion/cation ratios, where $\tau$ is the dimensionless time and relates to $t$ by $\tau = \kappa^2 D_{K^+} \cdot t$. ($\kappa$ is the inverse of Debye length $\lambda_D$ and $D_{K^+}$ is the diffusion coefficient for $K^+$). $\tau = 1$ is equivalent to $t = 0.424\, ns$. $\lambda_D$ is equal to 0.91 nm.

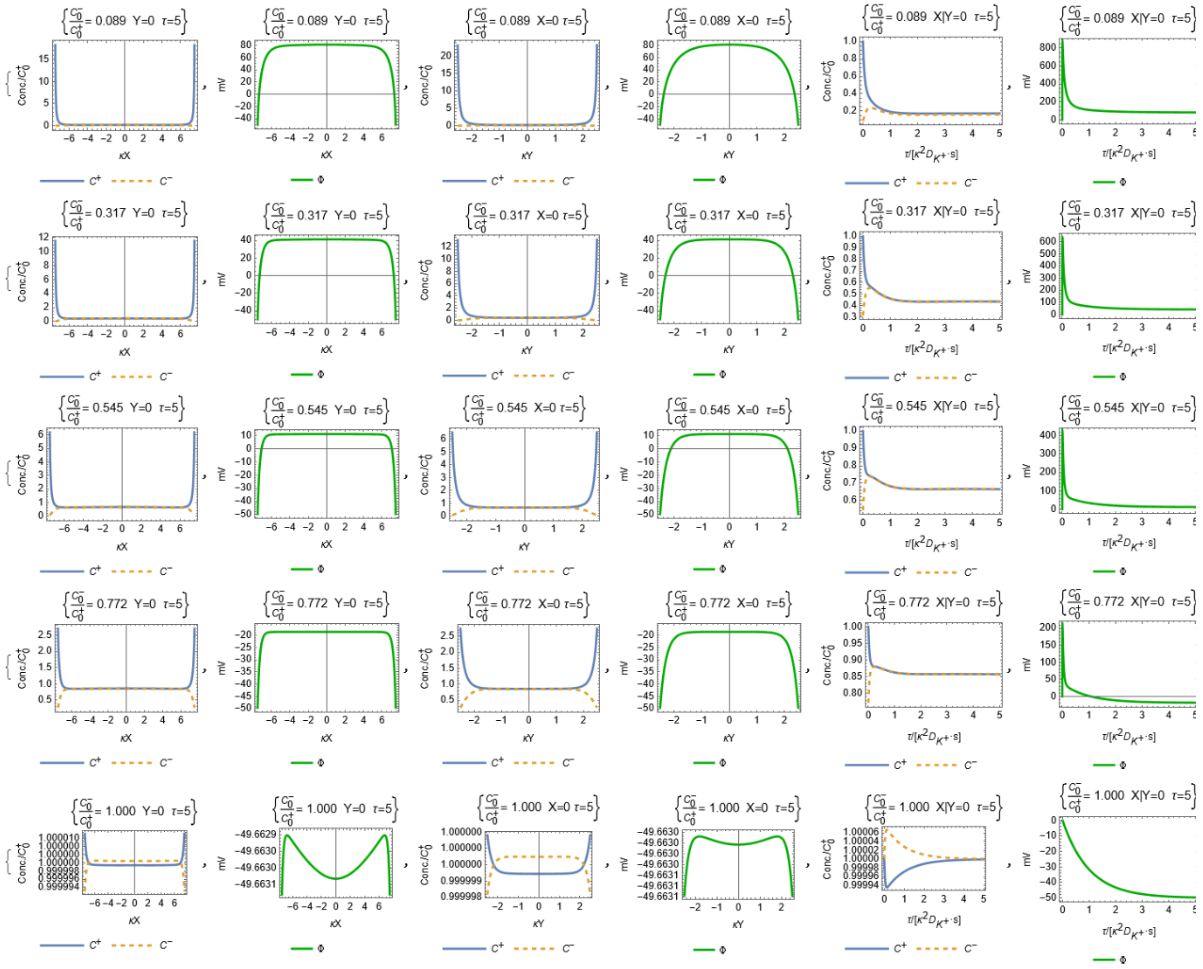


FIG. S4 The longitudinal and lateral concentration profiles [1st and 3rd column] and electrical potentials [2nd and 4th column] in the bounded, rod-shaped cell geometry [cell length/width: 15000/5000·$\kappa_D^{-1}$ nm, where $\kappa_D^{-1}$ was defined in Fig. S1] are illustrated for the anion/cation ratios of 0.089, 0.317, 0545, 0.772 and 1.000 [1st to 5th row]. The time courses from τ = 1 to 5 for the longitudinal and lateral concentration profiles and electrical potential [5th and 6th column] are shown for the same range of the anion/cation ratios, where $\tau$ is the dimensionless time and relates to $t$ by $\tau = \kappa^2 D_{K^+} \cdot t$. ($\kappa$ is the inverse of Debye length $\lambda_D$ and $D_{K^+}$ is the diffusion coefficient for $K^+$). $\tau = 1$ is equivalent to $t = 0.424\, ns$. $\lambda_D$ is equal to 0.91 nm.

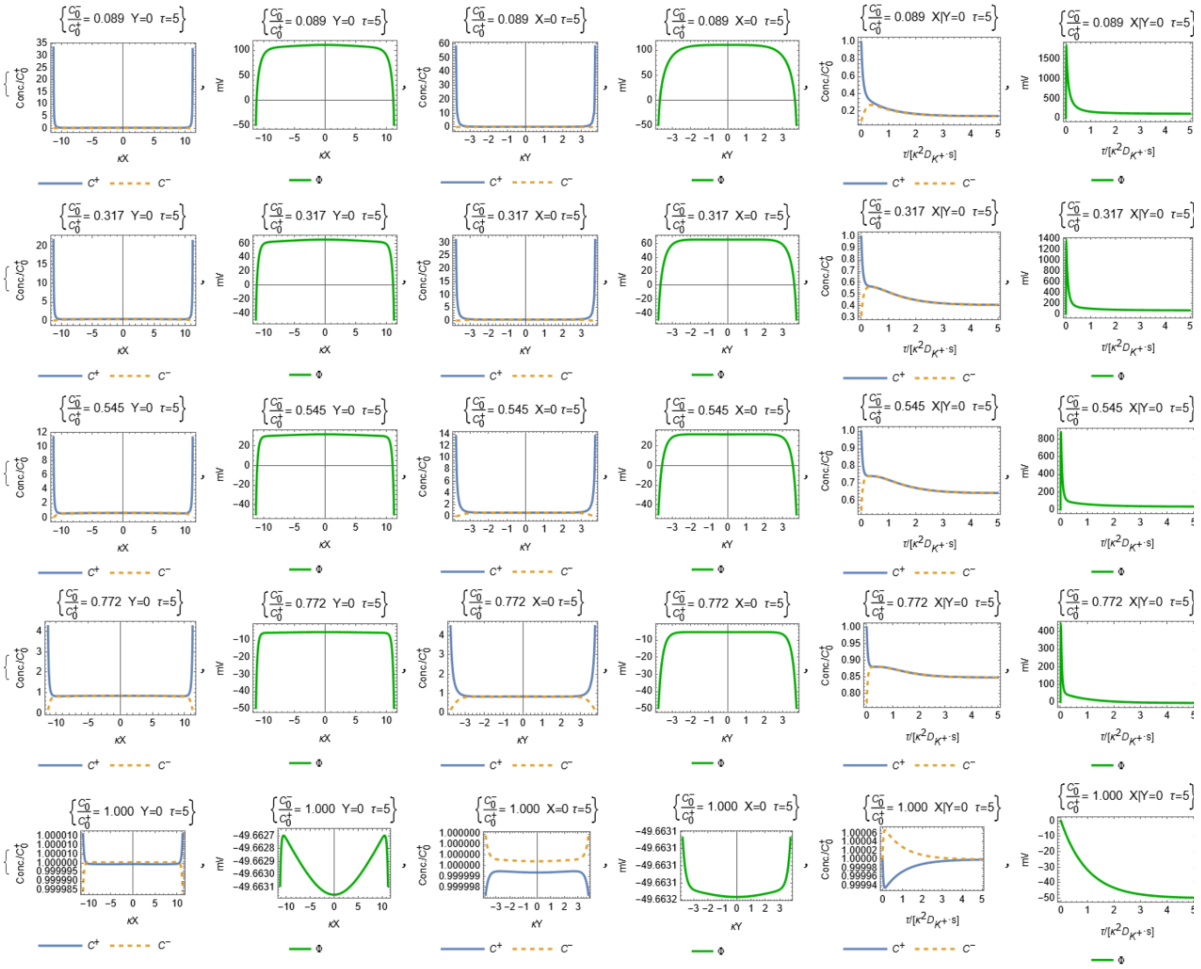


FIG. S5 The longitudinal and lateral concentration profiles [1st and 3rd column] and electrical potentials [2nd and 4th column] in the bounded, rod-shaped cell geometry [cell length/width: 225000/7500·$\kappa_D^{-1}$ nm, where $\kappa_D^{-1}$ was defined in Fig. S1] are illustrated for the anion/cation ratios of 0.089, 0.317, 0545, 0.772 and 1.000 [1st to 5th row]. The time courses from τ = 1 to 5 for the longitudinal and lateral concentration profiles and electrical potential [5th and 6th column] are shown for the same range of the anion/cation ratios, where $\tau$ is the dimensionless time and relates to $t$ by $\tau = \kappa^2 D_{K^+} \cdot t$. ($\kappa$ is the inverse of Debye length $\lambda_D$ and $D_{K^+}$ is the diffusion coefficient for $K^+$). $\tau = 1$ is equivalent to $t = 0.424\, ns$. $\kappa_D^{-1}$ nm is equal to 0.91 nm.

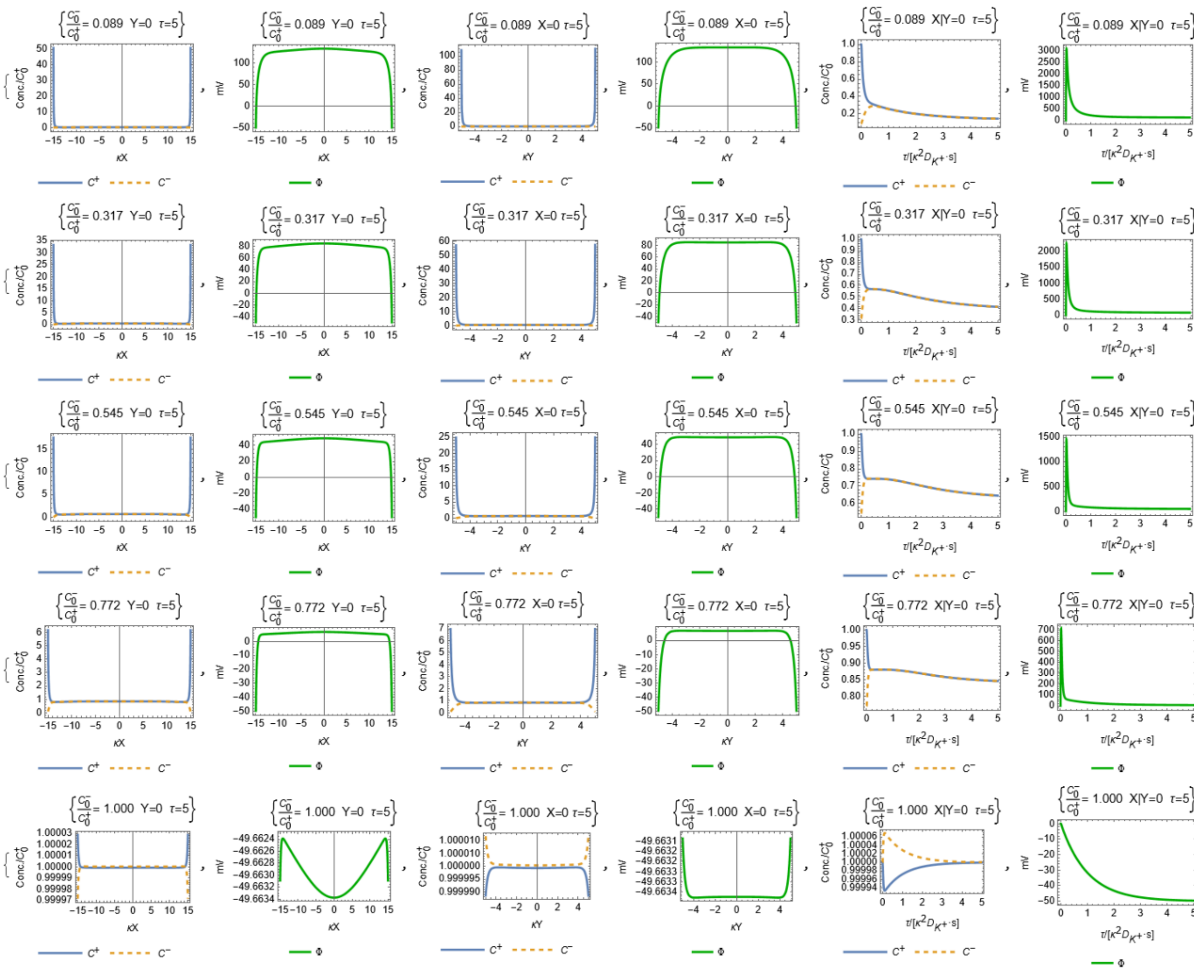


FIG. S6 The longitudinal and lateral concentration profiles [1st and 3rd column] and electrical potentials [2nd and 4th column] in the bounded, rod-shaped cell geometry [cell length/width: 30000/10000·$\kappa_D^{-1}$ nm, where $\kappa_D^{-1}$ was defined in Fig. S1] are illustrated for the anion/cation ratios of 0.089, 0.317, 0545, 0.772 and 1.000 [1st to 5th row]. The time courses from $\tau$ = 1 to 5 for the longitudinal and lateral concentration profiles and electrical potential [5th and 6th column] are shown for the same range of the anion/cation ratios, where $\tau$ is the dimensionless time and relates to $t$ by $\tau = \kappa^2 D_{K^+} \cdot t$. ($\kappa$ is the inverse of Debye length $\lambda_D$ and $D_{K^+}$ is the diffusion coefficient for $K^+$). $\tau = 1$ is equivalent to $t = 0.424\, ns$. $\kappa_D^{-1}$ is equal to 0.91 nm.

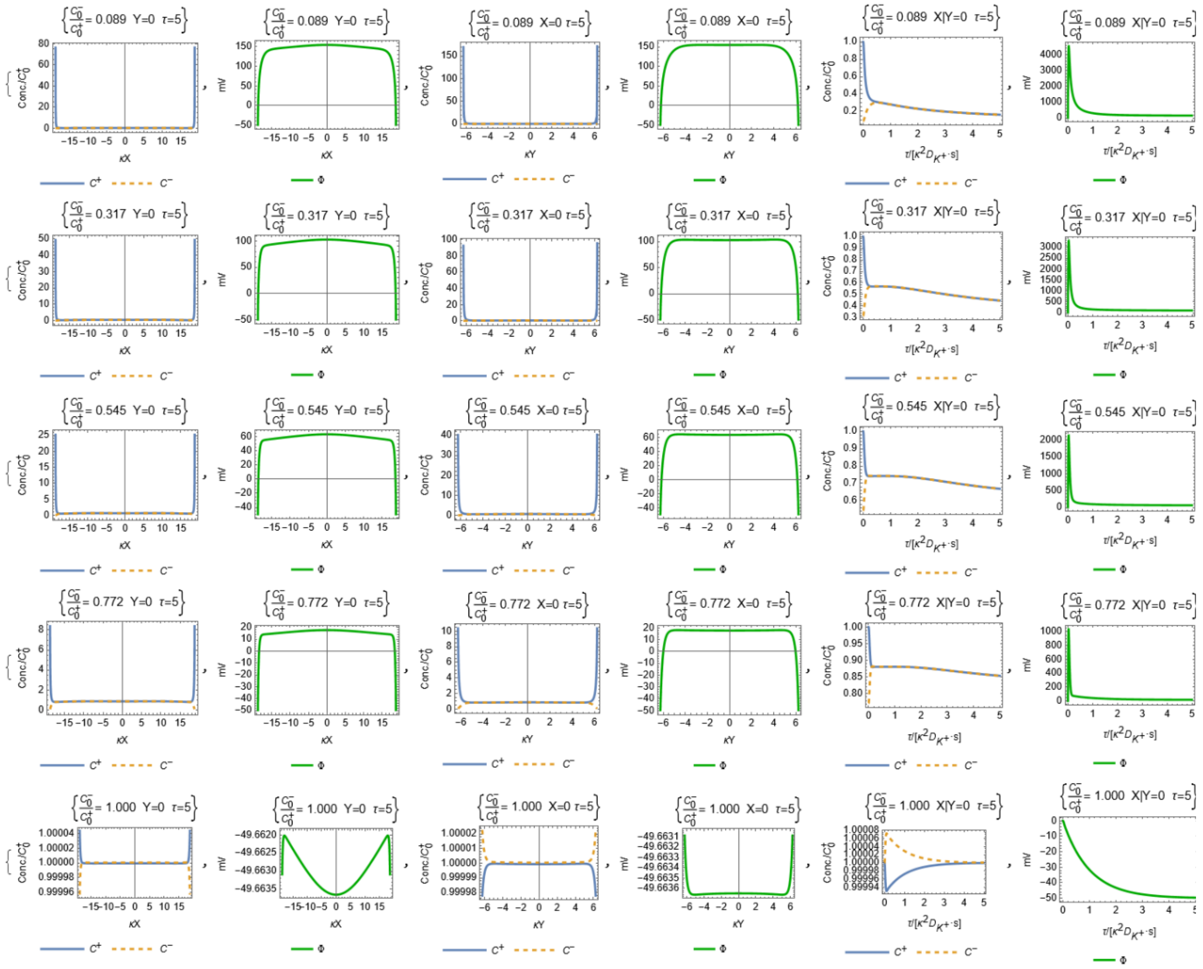

FIG. S7 The longitudinal and lateral concentration profiles [1st and 3rd column] and electrical potentials [2nd and 4th column] in the bounded, rod-shaped cell geometry [cell length/width: 37500/12500·$\kappa_D^{-1}$ nm, where $\kappa_D^{-1}$ was defined in Fig. S1] are illustrated for the anion/cation ratios of 0.089, 0.317, 0545, 0.772 and 1.000 [1st to 5th row]. The time courses from τ = 1 to 5 for the longitudinal and lateral concentration profiles and electrical potential [5th and 6th column] are shown for the same range of the anion/cation ratios, where $\tau$ is the dimensionless time and relates to $t$ by $\tau = \kappa^2 D_{K^+} \cdot t$. ($\kappa$ is the inverse of Debye length $\lambda_D$ and $D_{K^+}$ is the diffusion coefficient for $K^+$). $\tau = 1$ is equivalent to $t = 0.424\,ns$. $\kappa_D^{-1}$ nm is equal to 0.91 nm.

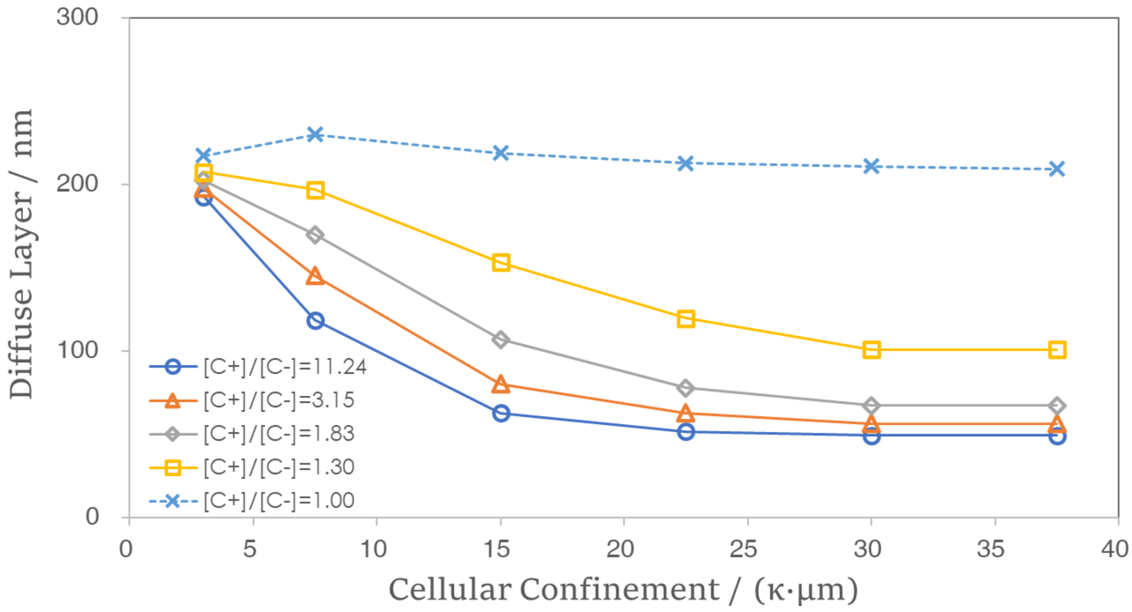


FIG. S8 The diffuse layer in relation to cellular confinement for the cation/anion ratios of 11.24, 3.15, 1.83, 1.30 and 1.00 (or the anion/cation ratios of 0.089, 0.317, 0545, 0.772 and 1.000). $\kappa_D^{-1}$ was defined in Fig. S1.

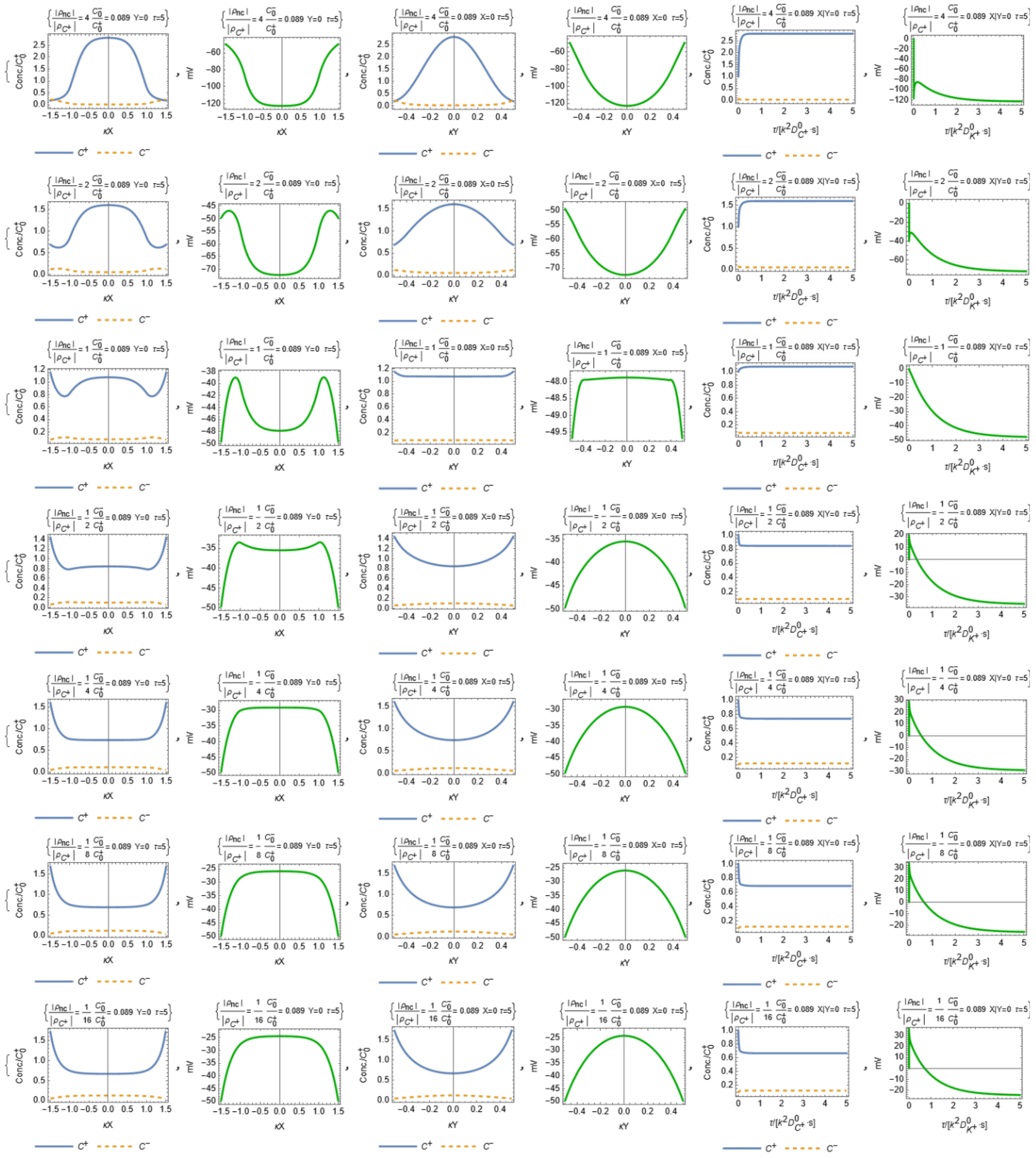


FIG. S9 With the nucleocytoplasmic (NC) ratio 0.52, the width of nucleoid is set as 700·$\kappa_D^{-1}$ nm. The longitudinal and lateral concentration profiles [1st and 3rd column] and electrical potentials [2nd and 4th column] in the bounded, rod-shaped cell geometry [cell length/width: 3000/1000·$\kappa_D^{-1}$ nm, where $\kappa_D^{-1}$ was defined in Fig. S1] including the nucleoid with various total charges are illustrated for the anion/cation ratio of 0.089. The time courses from τ = 1 to 5 for the longitudinal and lateral concentration profiles and electrical potential [5th and 6th column] are shown for the same range of the anion/cation ratios, where $\tau$ is the dimensionless time and relates to $t$ by $\tau = \kappa^2 D_{K^+} \cdot t$. ($\kappa$ is the inverse of Debye length $\lambda_D$ and $D_{K^+}$ is the diffusion coefficient for K+). $\tau = 1$ is equivalent to $t = 0.424\,ns$. $\kappa_D^{-1}$ nm is equal to 0.91 nm. The total charges of the nucleoid include $\rho_{nc}/\rho_{C^+} = \left(\frac{4}{1}, \frac{2}{1}, \frac{1}{1}, \frac{1}{2}, \frac{1}{4}, \frac{1}{8}, \frac{1}{16}\right)$ [1st to 7th row].

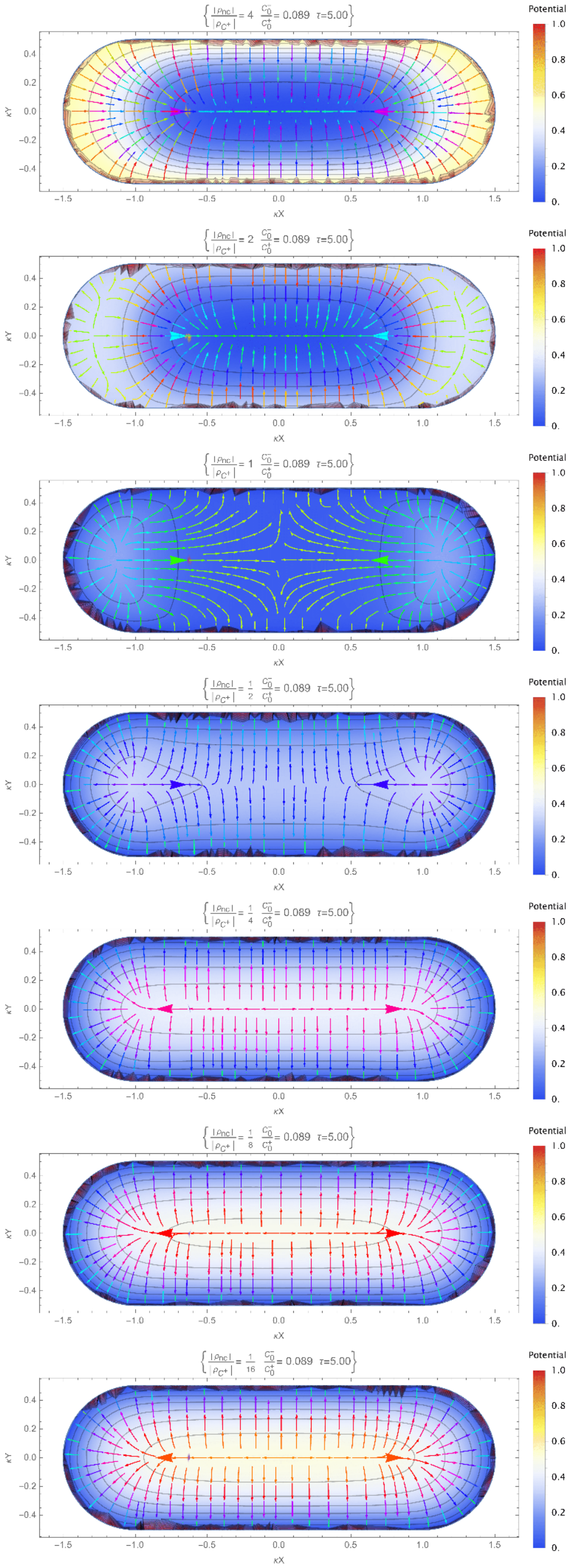

{ |ρnc| / |ρC+| = 4   C0− / C0+ = 0.089   τ=5.00 }
{ |ρnc| / |ρC+| = 2   C0− / C0+ = 0.089   τ=5.00 }
{ |ρnc| / |ρC+| = 1   C0− / C0+ = 0.089   τ=5.00 }
{ |ρnc| / |ρC+| = 1/2   C0− / C0+ = 0.089   τ=5.00 }
{ |ρnc| / |ρC+| = 1/4   C0− / C0+ = 0.089   τ=5.00 }
{ |ρnc| / |ρC+| = 1/8   C0− / C0+ = 0.089   τ=5.00 }
{ |ρnc| / |ρC+| = 1/16   C0− / C0+ = 0.089   τ=5.00 }
Potential
κX
κY

FIG. S10 With the nucleocytoplasmic ratio 0.52, the width of nucleoid is set as 700·$\kappa_D^{-1}$ nm, where $\kappa_D^{-1}$ was defined in Fig. S1. The electrical field (streamline arrows) and potential (contours) at $\tau = 5$ in the bounded, rod-shaped cell geometry [cell length/width: 3000/1000·$\kappa_D^{-1}$ nm] including the nucleoid with various total charges for the cation/anion ratios of 0.089. The potentials are normalized to the maximum and minimum levels among presented cation/anion ratios. The total charges of the nucleoid include $\rho_{nc}/\rho_{C^+} = \left(\frac{4}{1}, \frac{2}{1}, \frac{1}{1}, \frac{1}{2}, \frac{1}{4}, \frac{1}{8}, \frac{1}{16}\right)$ [1st to 7th row]. $\kappa_D^{-1}$ nm is equal to 0.91 nm.

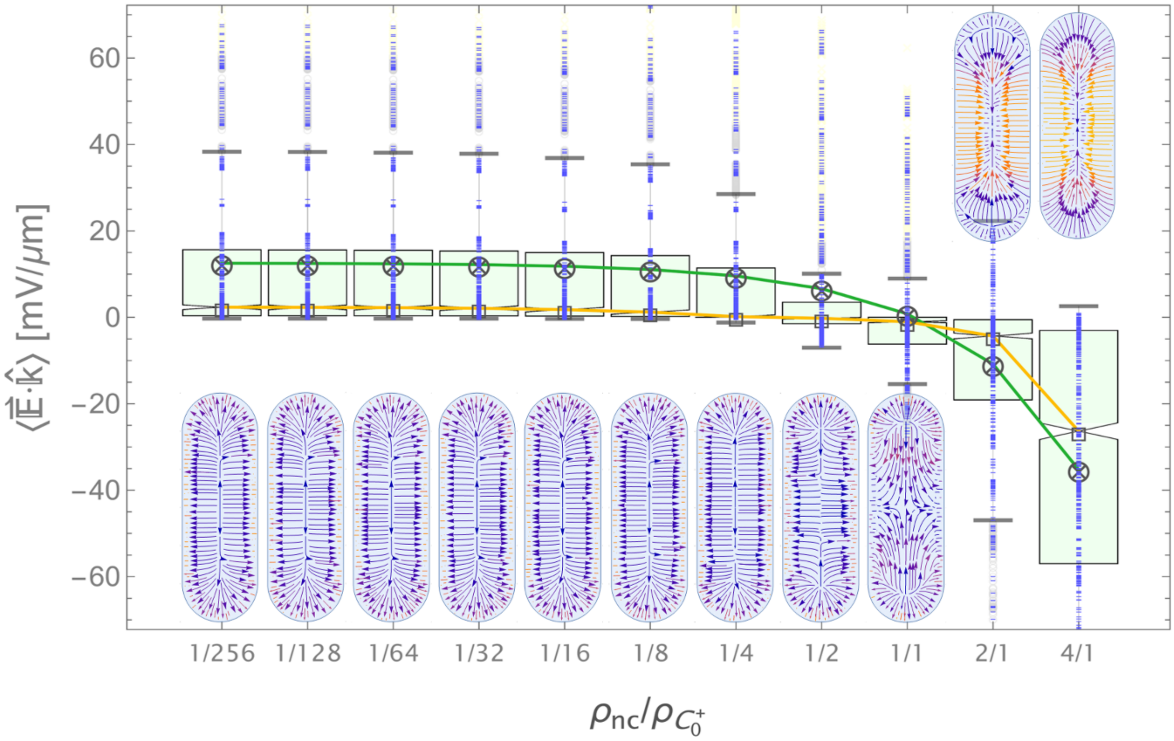


FIG. S11 The dependence of mean E-field strength upon the charge density of the nucleoid $|\rho_{nc}/\rho_{C^+}|$. In each column, box chart of the distribution of $\langle\vec{E}\cdot\hat{k}\rangle$ is plotted. Cross symbols: mean. Square symbol: median. Blue dots: data. The bottom and top of light green box indicate the 1st and 3rd quartile. Upper/lower horizontal bar: outlier bounds.

#### ▪ The Minimal Model for the Min-protein Oscillator

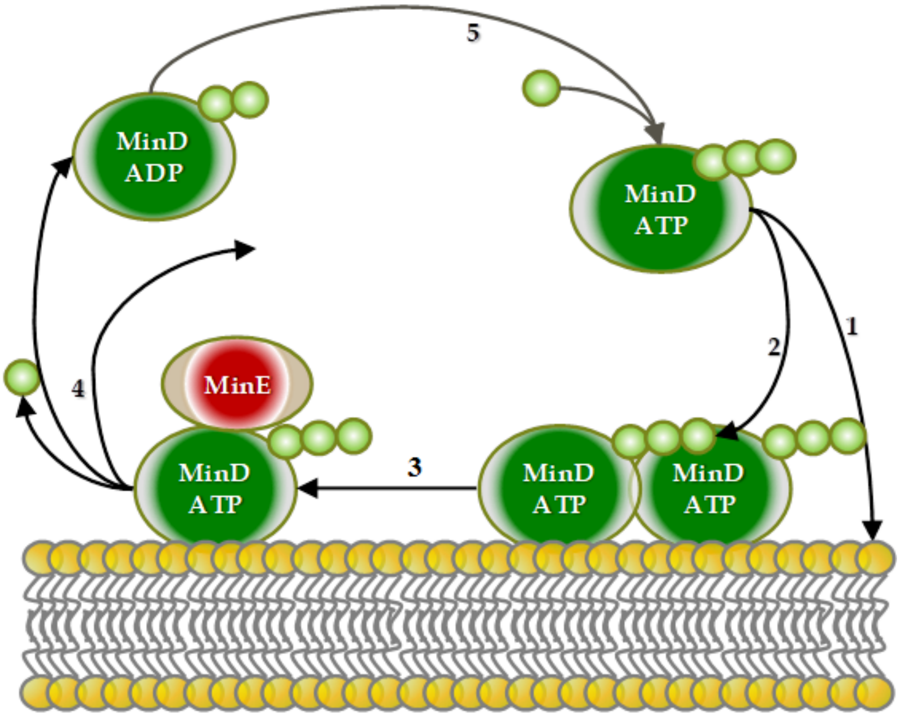


FIG. S12 The minimal model for the Min-protein oscillator. MinD-ATP: membrane-associated MinD. MinD-ATP: free-diffusing cytosolic MinD. The reaction of nucleotide exchanges between MinD-ADP and MinD-ATP is the only reaction in the cytosol. (1) MinD-ATP is hydrophobic and tends to be associated on the cell membrane. (2) Once associated on the cell membrane, more MinD-ATP and MinE are recruited to the cell membrane. (3) Membrane-associated MinE forms MinDE complex with MinD-ATP and activate the ATPase on the MinD-ATP for hydrolysis. (4) Then MinD-ADP and MinE return back to the cytosol. (5) The r nucleotide exchanges between MinD-ADP and MinD-ATP is the only reaction in the cytosol. [Figure revised from (*1*)]

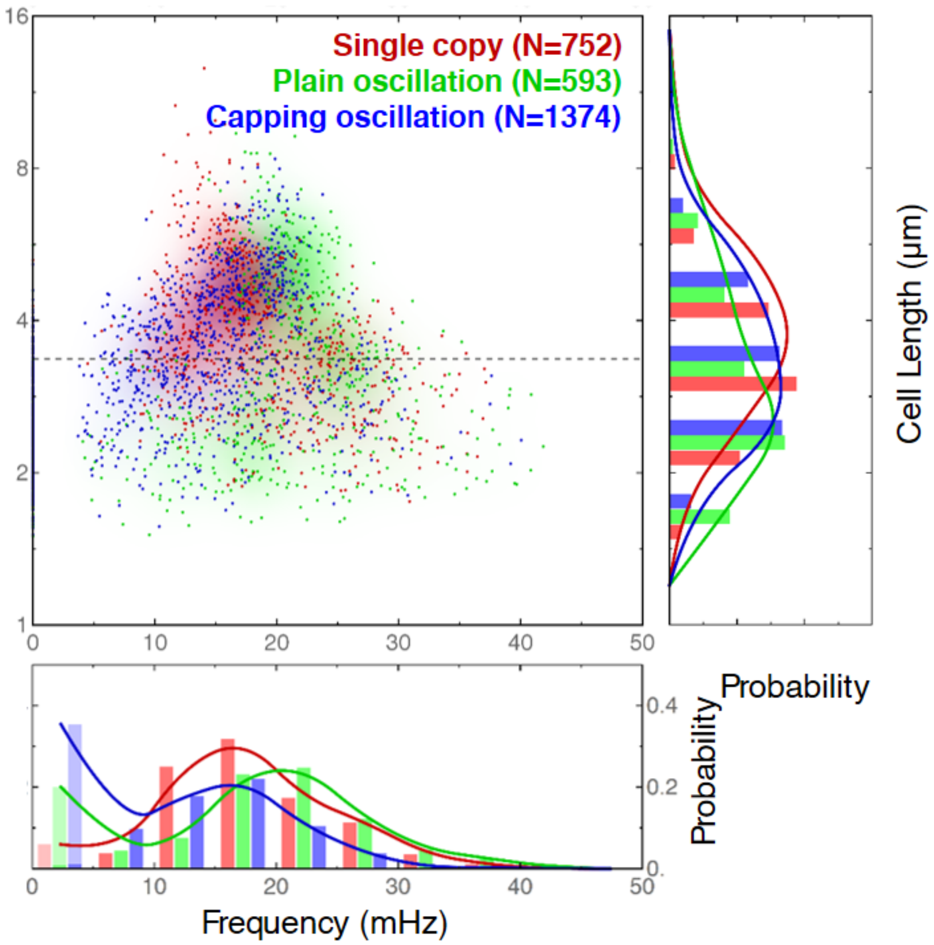


FIG. S13 The scatter plots for Min-protein oscillation frequency. Three sample groups are labeled by color: single-copy (red), plain oscillation (green) and capping oscillation (blue). Corresponding to the bottom scatter plot, histograms of oscillation frequency and cell length are displayed in bottom and right panels. The horizontal dash line in the scatter plot indicates the average of cell lengths (3.36 ± 0.02 μm) measured from all cell samples. The expression of Min proteins is under the control of single-copy $P_{minB}$ promoter; and under the control of $P_{lac}$ promoter by IPTG inductions. All expression levels of mCherry-MinD/MinE-YFP in each cell are displayed by normalization to the average of mCherry-MinD/MinE-YFP fluorescent intensities in the single-copy cell group expressing mCherry-MinD and MinE-YFP. The average levels of MinD (MinE) in the *minB*⁻ cells undergoing either plain or capping oscillations are estimated 2.05 ± 0.07 and 6.17 ± 0.12 (1.80 ± 0.07 or 5.74 ± 0.16; mean ± S.E.M.) folds higher than that in the *minB*⁻ cells hosting a single-copy operon [$P_{minB}$::mCherry-minD minE-yfp], i.e., 1.00 ± 0.67 (1.00 ± 0.49). [We thank Prof. Yi-Ren Chang for his help on experiments for this Figure.]

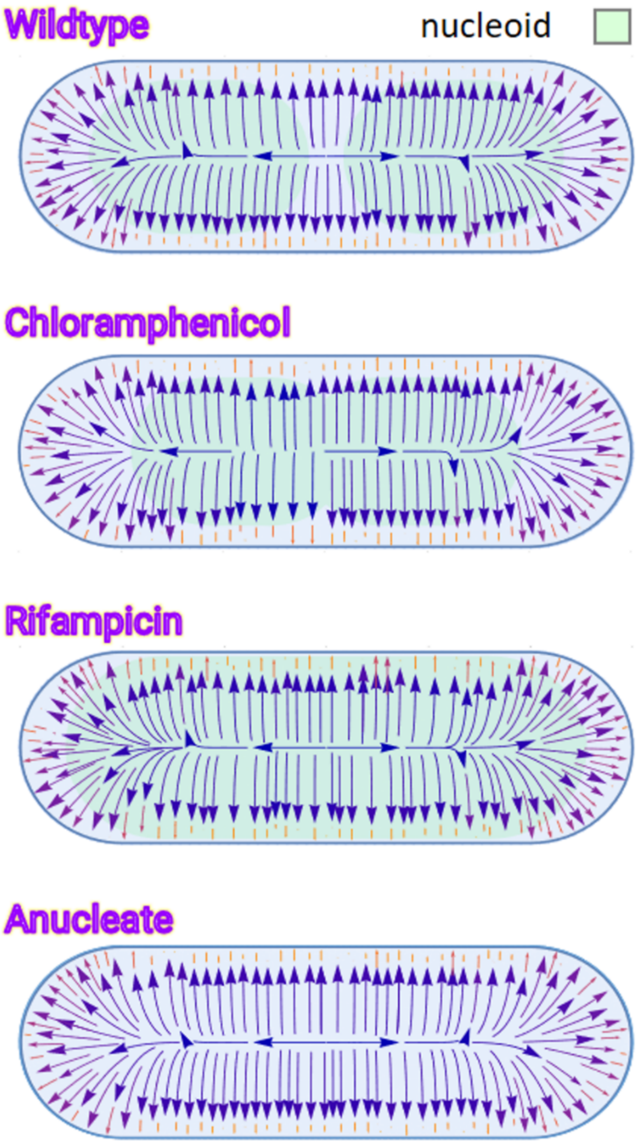


FIG. S14 The configurations of E-field and nucleoid for wildtype, chloramphenicol-treated, rifampicin-treated and anucleate cells.

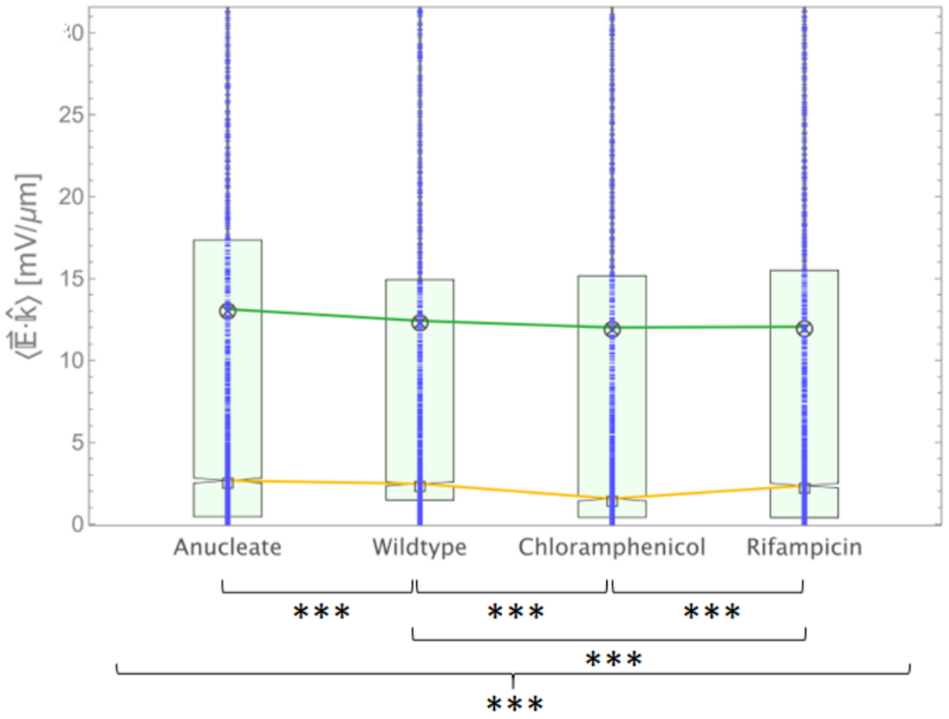


FIG. S15 The distributions of E-field strength and statistical analysis for wildtype, chloramphenicol- and rifampicin-treated and anucleate bacteria. Symbols of triple stars indicate significant difference of $p<0.001$ by either mutually statistical tests (medians) using non-parametric Mann-Whitney method or statistical tests among four distributions using both Krustal-Wallis method (medians) and K-samples T-test method (means).

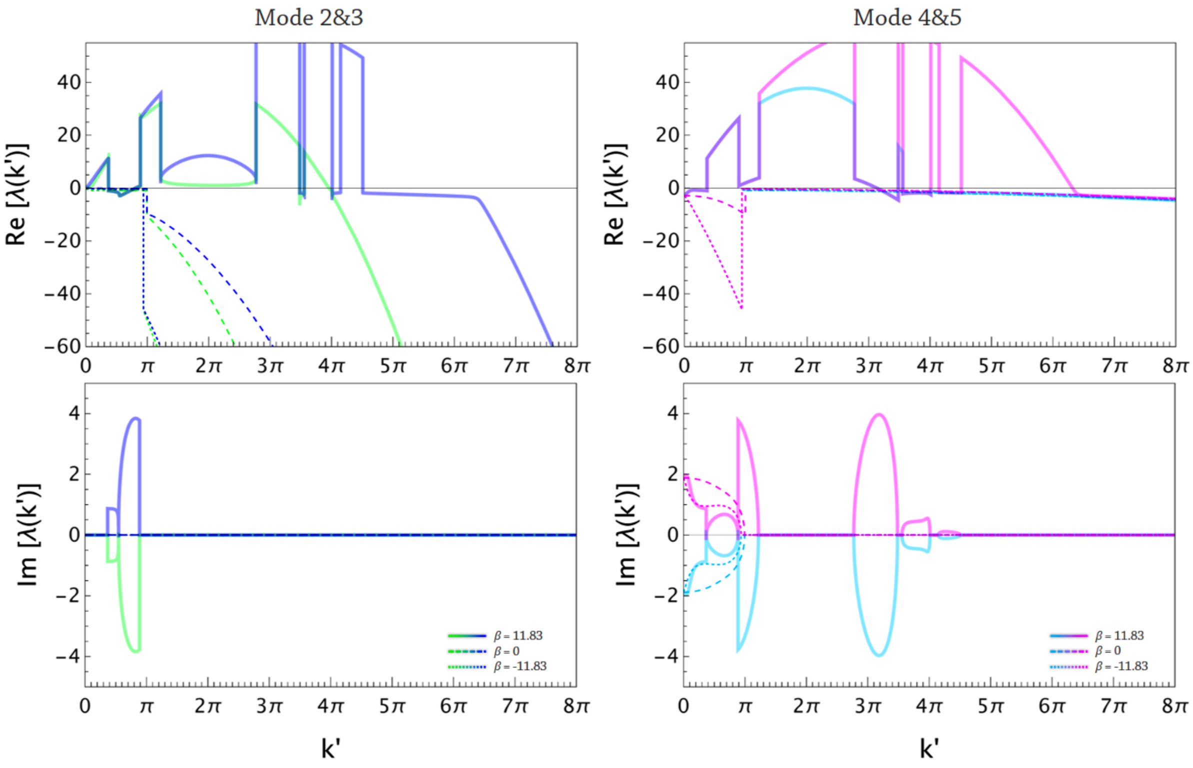


FIG. S16 The dispersion diagrams for Wave Mode 2&3 (left panels) and Wave Mode 4&5 (right panels). The real parts for all modes are in the upper panel and the imaginary parts for all modes are in the lower panels.

### A. Materials and Methods

#### Bacterial strains and growth conditions

*E. coli* strain YLS1 (*2*) transformed with plasmid pDE4-9b were used in differential expression of mCherry-MinD and MinE-YFP. The same strain transformed with single-copy mini-F plasmid pZC320-CDE was used to mimic the native expression level of MinD and MinE under the control of *minB* promoter in the single-copy *minB* cassette under MC1000/*min*⁻ background. In the experiments of differential expression, bacteria grown in Luria-Bertani medium (1% wt tryptone, 0.5% wt yeast extract, and 1% wt NaCl, BD Difco, USA) supplemented with 0.4% glucose and 100μM ampicillin were cultured overnight at 37°C. In each case, a 1/20 dilution of overnight culture was grown in the same medium at 37°C until O.D. ca. 0.6 – 0.8. After 3 times washes of M9 medium, the cultures were resuspended in M9 medium supplemented with 400 μM isopropyl β-D-1-thiogalactopyranoside (IPTG) and antibiotics for induction of protein expressions and grown at 20°C for 40, 80, 120, 160, 200 and 240 minutes depending on expression levels. Finally, the cultures were ready for in vivo time-lapse fluorescence imaging after the removal of inducer by washes of M9 medium supplemented with 0.4% glucose. All experiments were conducted at 25°C.

#### Plasmid constructions

All plasmids were constructed by In-Fusion Cloning method (Clontech Laboratories, Inc., USA). The method seamlessly and directionally fuses two or multiple linearized DNA fragments with 15bp overlap by in-fusion enzymes. The linearized DNA fragments can be derived from any source by PCR. All the PCR fragments were synthesized by PrimStar MAX DNA polymerase (Takara Bio Inc., Japan) to ensure their accuracy and fidelity up to 10kb.

Plasmid pDE4-9b was constructed from plasmid pSOT37 via sequential constructions of intermediate plasmids. The detailed process has been described in prior work (*3*). Plasmid pZC320-CDE was constructed by fusing fragment [$P_{lacI}$::*lacI t1-rrnB*::*cfp*::P*tet* $P_{lac}$:: *mcherry-l16d-minD minE-l16c-yfp*] (from plasmid pDE4-9b) and [*oriT* ::*repE incC* $P_{sop}$::*sopA sopB sopC* $t_{\Omega}$ *bla*::$P_{aadA}$] (from plasmid pZC320) by primers 1124-1127.

Table S1 Primers.

| Primer | Sequence (5' →3' ) |
|---|---|
| 1124 | gccttgatgttacccgagag |
| 1125 | gtccagaaccttgaccgaac |
| 1126 | cgggtaacatcaaggcgagctcagaggccctttcgtcttcaa |
| 1127 | ggtcaaggttctggactccgcgtttccagactttac |

**Microscopy**

Fluorescent Min proteins were visualized in live bacteria via wide-field time-lapse imaging on an Olympus IX71 inverted microscope (Olympus Corp., Japan) with a 100x PlanAPO oil objective (N.A. 1.4). YFP and mCherry fluorescent proteins were excited with a 488 nm and 561 nm lasers (DPSS Laser, TWC Opto Corp., Taiwan) through appropriate neutral-density filters (Edmund Optics, USA) and a filter cube (Olympus Corporation) housing multiband filter set (LF405/488/561/635-A-000; Semrock Inc., USA). CFP, msfGFP and mCherry were excited using a 175 W Xenon lamp housed in the wavelength switcher Lambda DG-4 (Sutter Instrument, USA) with appropriated band-pass filters and visualized through a filter cube housing multiband filter set (CFP/YFP/HcRed-3X-A-000; Semrock Inc., USA). An electron-multiplying charged-coupled device camera (C9100, Hamamatsu Photonics) was employed in image acquisitions of Min oscillation at a frame rate of 0.5 Hz.

**Image and Data Analysis**

MATLAB (MathWorks Inc.) -based programs were custom-coded for image and data analysis. To acquire cell boundaries, bright-field images or the average of all fluorescence stacks for a bacterial cell were analyzed with PSICIC (*4*). In order to estimate the image background for the stack-averaged image, the intensity of pixels enclosed by each cell boundary was filled with the value of averaged intensity over the cell boundary obtained from the stack-averaged image to get an approximated background image. Then we may obtain the image background by fitting the approximated background image to a 2D Gaussian distribution since the later profile of laser illumination projected to sample plane is a 2D Gaussian function. After background correction, the expression level for each bacterial cell was then determined by the averaged intensity retrieved from all fluorescence stacks over the interest of region defined by the cell boundary. The cell length was measured based on the longitudinal midline determined by self-coded Matlab algorithm and/or PSICIC. Besides, we measured the fluorescence intensity of one polar zone over time to acquire the time course of fluorescence intensity. The method of empirical mode decomposition (EMD) used in Hilbert-Huang Transformation (*5*) was adopted to remove high frequency noises and variations. The period of Min oscillation was estimated by the averaged interval between intensity peaks calculated from the EMD-processed time course of fluorescence intensity. Peak-to-peak variation was calculated by the standard deviation of the mean period from the processed time course of fluorescence intensity.

**Nucleocytoplasmic Ratio Approximation**

To visualize the nucleoid for size approximation, bacterial cells were cultured to experimental conditions and subject to 5 minutes 1μM staining by 4', 6-diamidino-2-phenylindole (DAPI). Stained cells were then washed three time by M9 medium and resuspended in the same medium for later fluorescence imaging as described in above, but using appropriated band-pass filter (FF01-356/30-25, Semrock Inc., USA) in the wavelength switcher Lambda DG-4 (Sutter Instrument, USA) for excitation. The acquired DAPI-stained image stacks were segmented using the open-source image analysis software Oufti (*6*). Nucleocytoplasmic ratio for single cells was approximated by dividing the nucleoid area by the cell area for each cell in according to the reference (*7*). The nucleoid and cell areas were calculated from the segmented area through the images analysis of DAPI-stained and phase contrast images, respectively.

**Estimation of Nucleoid Charge Density**

In chloramphenicol-treated cells, the nucleoid morphology is compressed and the NC ratio is ~ 0.47 in our simulations (Fig. 4J). Even though the charge density of nucleoid in chloramphenicol-treated cells appears to be higher than that of wildtype cells, incomplete blockade of translations by 30 min treatment of chloramphenicol allows further synthesis of nucleoid-associated proteins up to two-fold amount of that in the growth phase as their transition into stationary phase (see Table 1 in (*8*)). Therefore, the charge amounts carried by the nucleoid in chloramphenicol-treated cells would instead increase owing to the increase of nucleoid-associated proteins (see Appendix). The charge density of nucleoid in these cells would eventually increase up to about 1.17 folds in simulations. At last, in rifampicin-treated cells, the nucleoid expands to the NC ratio ~ 0.8 (Fig. 4A and 4D). Though the charge density of nucleoid is supposed to reduce due to the expansion of nucleoid, the total charges carried by the expanded nucleoid include more DNA, RNA and proteins. Earlier studies report that about 80% of DNA, 10% of RNA and 10% proteins are included in the normal nucleoid (*9, 10*) such that the charge amounts might increase about 4–5 folds. Therefore, the resulting condition about the increase of charge density up to 2.5 folds for the expanded nucleoid will be considered in evaluation of E-field strength.

#### B. Discrepancies of Electroneutrality in Real Cellular Context

Electroneutrality is a basic assumption in classical electrolyte theory, shaping the key concepts in electrochemistry and electrophysiology (*11*). It suggests that charge separation or excess charge does not occur within the solution, ensuring that neutrality is maintained throughout. Coulomb forces between ions in an electrolyte solution work to preserve macroscopic electroneutrality, as any spatial charge imbalances in the bulk are immediately neutralized by ion movement to minimize the electrochemical potential. At charged interfaces, the electrolyte compensates for surface charges by forming electric double layers (EDLs), which are not electroneutral. In electrostatic equilibrium, these double layers prevent potential differences between the interfaces and prevent the formation of a macroscopic E-field in the bulk (Fig. 1A). However, this physical understanding may be challenged when the assumptions underlying electroneutrality are invalid, especially considering that an E-field in an electrolyte can only exist due to net charge separation, creating an inherent paradox in the principle (*12*). For dynamic problems, such as ion channel-mediated current around the cell membrane (*13, 14*) and non-stationary liquid junction potential during the continued dialysis (*15*), emergence of excess charge at the interface generates a local E-field and induces extension of diffuse layer away from the interface. Alternatively, for steady-state problems such as electrochemical cells with liquid electrolytes, 1-2 nm range of EDL around the cell membrane and electroneutral bulk are derived by Poisson–Boltzmann (PB) equation based on the Gouy–Chapman theory when electroneutrality assumption is maintained at infinity and cytosolic cationic/anionic charges are equal in the bulk (*11*). Whereas, the validity of both conditions may be questioned for micron-sized bacterial cells.

First, the effect of intrinsic E-field generated by solvated ions in the immediate vicinity exhibits substantial variation among distinct solvated ionic species with identical charges. For monoatomic ions like $K^+$, which is the most abundant cation in bacterial cytosol, the hydration shell is thinner, less structured, and more tightly bound compared to atomically smaller (e.g., $Na^{+}$) or multivalent ions (e.g., $Ca^{2+}$) (*16*). By contrast, for cell-impermeable anions (e.g., proteins, phosphates, sulfates and other organic macromolecules) like glutamate ion , which is the most abundant anion in bacterial cytosol, carboxylate and amino groups form multiple hydrogen-bonding with surrounding water molecules and the hydration shell is thick when concentration is less than 300 mM (*17*). High hydration water number revealed in experiments and simulations suggest that all first-shell water molecules are retarded compared to bulk water and even the second hydration shell is dynamically affected. Exceptionally, monoatomic ions with thick hydration shell behave like electrically neutral particles (*16*). This might not be true for small macromolecule ions, but still implies the intrinsic field effect generated by glutamate ions in the immediate vicinity is significantly reduced compared to monoatomic ions with thinner hydration shell such as $K^{+}$ and $Cl^{-}$ (*18*). Further, distinct size of solvated equal-valent ionic species would induce different charge density distribution, hence, complex water-ion and water-water interactions in the hydration shell (*16*) which leads to differential screening effect over ionic field.

Notably, the concentrations of key intracellular electrolytes of *Escherichia coli* during its growth phase have been experimentally determined, including 217-257 mM $K^{+}$ (*19, 20*), 5 mM $Na^{+}$ (*21*), 8-33 mM $Cl^{-}$ (*20*), 1-10 mM Pi (*22, 23*), 90 nM $Ca^{+2}$ (*24*), 1-2 mM $Mg^{+2}$ (*25, 26*), 30-60 μM $Fe^{+2}$/ $Fe^{+3}$ (*27*) and 0.2 mM $Zn^{2+}$ (*26*). Some anionic metabolites in the cytosol are maintained with millimolar concentration, such as 96 mM glutamate, 17 mM glutathione and 4.2 mM aspartate (*28*). Even though the total concentration of all metabolites is reported as much as 230 mM, ca. 350-500 mM intracellular $K^{+}$ concentration can be accumulated in the cytosol for high metabolically active bacteria (*29*), indicating the intracellular concentration of $K^{+}$ constantly surpasses that of other cations, as well as that of anions like $Cl^{-}$, Pi and metabolites. Therefore, not only the total effective cationic/anionic concentration, but also the total effective charges carried by cations/anions are substantially imbalanced in bacterial cytosol, leading to a significant deviation from bulk electroneutrality as this charge-imbalanced electrolyte solution is enclosed in a finite, subcellular space invalidating unlimited supplies of counterions to balance the excess charge in the bulk (Fig. 1B). Further, those negatively-charged macromolecules allocated to the nucleoid, such as DNA, RNA, and assorted metabolites (nucleotides and nucleoid-associated proteins), are much less mobile compared to electrolytic background, at least to the time scale for electrolyte redistribution to achieve equilibrium, acting like an immobile negatively charged region over the central part of bacterial cell (Fig. 1C). Taken these contexts together, the physical conditions to support the electroneutrality assumption are not evidently justified in the cytosolic bulk of micron-sized bacteria, implicating the possibility of intracellular E-field generated by excess charge in the bulk and asymmetric charge distribution in EDL. The latter may also bring an extended diffuse layer around the cell membrane much longer than typical Debye length (*14*).

To better address these discrepancies, we employed the Poisson-Nernst-Planck (PNP) equations which, when solved numerically, can provide a more accurate representation of ionic behavior in biological systems without relying on a priori assumption of electroneutrality (*30, 31*). While still as a continuum approach, the PNP model has proven to be computationally efficient and widely used in biophysics (*30, 31*). It has been applied to study phenomena when electroneutrality is not held such as the electrodiffusion of neurotransmitters across neuronal membranes (*11, 32, 33*), the formation of liquid junction potentials in extracellular spaces (*11, 34-38*), and ion gating, selectivity, and transport in ion channels (*39, 40*). Theoretical models suggest that ionic influx across the cell membrane can lead to the development of a local electric potential, extending the diffuse layer up to 100 nm around the cell membrane (*14*). For a micron-sized bacterium, the extended diffuse layer around cell boundary can significantly impact the mobility and organization of subcellular biomolecules. Recent studies reveal that, in addition to the mechanism of entropic-driven demixing stemming from cytosolic polydispersity, geometrical confinement and nucleoid structure, electrostatic mechanism also play an important role in charge-dependent protein mobility, segregation of subcellular biomolecules into specific regions and formation of protein-ribosome coacervate compartment in bacteria (*41-45*). While these studies focus on local protein-ribosome surface charge interactions, they do not rule out the role of the cell's pervading electric field. Generated by the extended diffuse layer near the boundary, this intracellular E-field could electrostatically organize charged biomolecules, driving their spatial localization (*45*) or maintaining equilibrium.

#### C. Models for Intracellular Electric Potential and Chemical Wave – Electric Field Interaction

##### Modeling Intracellular Electric Potential in Bacteria

Taking *Escherichia coli* during its growth phase as an example, the concentrations of key intracellular electrolytes are well-documented, including 217-257 mM $K^+$ (*19, 20*), 5 mM $Na^+$ (*21*), 8-33 mM $Cl^-$ (*20*), 1-10 mM Pi (*22, 23*), 90 nM $Ca^{+2}$ (*24*), 1-2 mM $Mg^{+2}$ (*25, 26*), 30-60 μM $Fe^{+2}$/ $Fe^{+3}$ (*27*) and 0.2 mM $Zn^{2+}$ (*26*). These values reveal an apparent imbalance in charge distribution. It is well acknowledged that water molecule is permanently dipolar and its peculiar microscopic network structure constantly undergoes topological reformation through hydrogen bonding (*46*). Given that the concentration of all metabolites is reported to be as much as 230 mM in *E .coli* (*28*) and even the net charge carried by many of them is not nil, direct interactions between ionic metabolites and surrounding water solvent facilitate these molecules readily accommodated into the hydrogen-bonding network of hierarchical water structure. The resulting structure and bonding thereby restrict intermolecular interactions in the immediate vicinity of surrounding water molecules or other ionic molecules. For example, glutamate ion as the most abundant anions in *E. coli* also acts as a permanent dipolar molecule at pH 7 and as such the structure and bonding of proximal glutamate-water, glutamate-glutamate and glutamate-cation interactions have been illustrated in a series of studies (*17, 47, 48*). Structural information extracted by neutron diffraction experiments depicts that each carboxylate oxygen/amine hydrogen atom in glutamate ion forms an average of three/one hydrogen bonds with water molecules (*48*); while clusters of glutamate dimers (*47*) and formation of aqueous $Na^+$-Glutamate ion pairs (*17*) are characterized by spectral experiments and molecular simulations. But contrary to the constraint between glutamate ions and surrounding water molecules via hydrogen bonding, solvation of $K^+$, due to its high ionic field strength, is thought to be a structure-breaker that aligns water dipoles radially in the first hydration shell, but causes the breaking of the bridging hydrogen bonds forming between the first and second shells, wherein water structure network is collapsed and water dynamics is accelerated (*16, 18, 49*). All these ion-water and water-water interacting events have been experimentally shown to repeat continually in the time scale of 10 to 100 picoseconds. By incorporating these microscopic findings of ion-water interactions with mesoscopic ion-ion interactions in the cellular context, an electrolyte such as $K^+$ experiences an ensemble average of dielectric frictions along with hydrodynamic friction in the solution, and as such the ensemble ion-water interactions considered on the basis of continuum model can be simply dictated by the macroscopic properties of water, i.e. dielectric polarization and fluidic viscosity. By contrast, a dipolar molecule such as glutamate ion is repeatedly constrained by the continually making and breaking of hydrogen-bonding and accommodated into water structure network. Accordingly, dipolar molecules in the cellular context, owing to their charged residues continually restricted in formation of hydrogen bonds with surrounding water molecules and screened by thick hydration shell, can be counted electroneutral-like in the mesoscopic scale, even these dipolar metabolites and water molecules are always in diffusion. Concisely, we mesoscopically view monoatomic ions as fast-moving, point-like electric sources and ionic metabolites as moving, electroneutral, ultrafast-shifting but constrained water-metabolite complex.

Next, as the intracellular content of monovalent cation $K^+$ surpasses that of $Na^+$ and other divalent cations, as well as that of anions $Cl^-$ and Pi, imbalanced charge distribution and thereof deviation from electroneutrality in subcellular space are very likely to build up a substantial level of electric potential. But for those negative charge carriers allocated to the nucleoid, such as DNA, RNA, an eclectic mix of metabolites such as nucleotides and nucleoid-associated proteins, exhibit much less mobile in comparison with electrolytic background, and at least to the time scale for ionic redistribution to achieve equilibrium, act much akin to an immobile, negative charged region over the central part of bacterial cell. Finally, taken these physical contexts together, insomuch as electric potential could be generated to a substantial level by intracellular ionic media, long-range electrostatic interaction may not be a negligible variable in the studies of biomolecular transport and related electrochemical phenomena in bacteria.

Considering that monovalent cation $K^+$ and anions $Cl^-$ and Pi are dominant in *E. coli* cytoplasm, as well as in which mesoscopic electroneutrality of abundant glutamate and aspartate anions is caused by hydrogen-bonded ionic metabolite-water structures, we construct the model for intracellular electric potential generated by ionic medium comprised of these electrolytes. The model based on PNP equations are described by

$$\frac{\partial C^+}{\partial t} = D_{C^+}\left[\nabla^2 C^+ + \frac{z^+F}{RT}\nabla\cdot(C^+\nabla\phi)\right] \tag{1}$$

$$\frac{\partial C^-}{\partial t} = D_{C^-}\left[\nabla^2 C^- + \frac{z^-F}{RT}\nabla\cdot(C^-\nabla\phi)\right] \tag{2}$$

$$\nabla^2\Phi + \frac{4\pi F}{\epsilon_s\epsilon_0}\sum(z^+C^+ + z^-C^-) \tag{3}$$

, where $C$, $D$, $z$ and $\Phi$ are the concentration, diffusion coefficient, valence and electric potential for the dominant monovalent ions and the electric potential set up by these ions in the system and superscripts + and – denotes cation and anion respectively. $F$, $R$, $T$, $\epsilon_0$ and $\epsilon_s$ have their usual meanings of Faraday constant, gas constant, absolute temperature, vacuum and relative permittivity. Equations (1) and (2) are the continuity, or Nernst-Plank equations, and relate to Poisson equation by summation of ionic charge densities derived from the concentrations of dominant ions as shown in (3). Due to rotational symmetry around the longitudinal axis of a rod-shaped cell, the dimension is reduced to two in the model (Fig. 1D). Equations (1)-(3) can be further transformed to their dimensionless forms by defining

$$u = \frac{C^+}{C^0_{K^+}}, v = \frac{C^-}{C^0_{K^+}} \ and \ w = \frac{F}{RT}\Phi \ . \tag{D1}$$

$C^0_{K^+}$ is the total concentration of $K^+$. The diffusion coefficients are normalized to the diffusion coefficient of $K^+$:

$$d_i = \frac{D_i}{D_{K^+}} \ . \tag{D2}$$

and dimensionless time and space coordinates (original unit: nm) are defined as:

$$X = \kappa \cdot x/1000 \ \ and \ \ Y = \kappa \cdot y/1000 \ . \tag{D3}$$

$$\tau = \kappa^2 \cdot D_{K^+} \cdot t \tag{D4}$$

where

$$\kappa = \frac{F^2 C^0_{K^+}}{RT\epsilon_s\epsilon_0} \tag{D5}$$

Then the PNP equations are obtained in dimensionless coordinates:

$$\hat{\nabla}^2 u + z^+ \hat{\nabla}\left(u \hat{\nabla} w\right) - \frac{1}{d_u}\frac{\partial u}{\partial \tau} = 0 \tag{4'}$$

$$\hat{\nabla}^2 v + z^- \hat{\nabla}\left(v \hat{\nabla} w\right) - \frac{1}{d_v}\frac{\partial v}{\partial \tau} = 0 \tag{5'}$$

$$\hat{\nabla}^2 \mathrm{w} + 4\pi(z^+ u + z^- v) = 0 \tag{6'}$$

where $\hat{\nabla}$ and $\hat{\nabla}^2$ are the operators working in X-Y coordinates.

To solve Equations. (1) to (3) applied to the 2-dimensional projected geometry of a rod-shaped cell, numerical solutions corresponding to assorted physical configurations are implemented in *Mathematica* (Wolfram, 2024) and by finite element method with Dirichlet boundary condition $\Phi_{membrane} =$ -50 mV at the cell boundary (*50*) and Neumann boundary condition $\hat{n} \cdot \nabla C^{\pm} = 0$, wherein $\hat{n}$ is the unit vector normal to the geometric border and the conditions depict no ionic flux may cross the cellular periphery and a constant surface potential is postulated. To overcome the difficulty in solving elliptic partial differential equations (PDE) in most solvers, such as Poisson's equation in (3), the method of false transients is adopted in the efficient solver for initial value problem (IVP) system of ordinary differential equations (ODE) or differential algebraic equations (DAEs) through arbitrarily introducing a pseudo time derivative to modify the elliptic PDE to a parabolic PDE, such as to allow the solution to be determined by marching in pseudo time to a steady state condition (*51*). The physical scheme stated above is exploited to figure out the alterations in the extension of diffuse layer due to bounded cellular space. Given imbalanced electrolytes as the solely cellular content, the nucleoid is modeled, to the time scale for ionic redistribution to achieve equilibrium, as the region with immobile, unform distribution of negatively-charged carriers allocated to the central part of the cell. Since information about the charge number carried by the nucleoid is lacking but speculated, the charge density, or total charge number of the nucleoid, is the parameter to be investigated in the simulations.

##### The comparison of BZ system and the Min-protein oscillation

Next to the understanding of the intracellular buildup of electric potential in the wake of imbalanced electrolyte distribution, how protein gradients (chemical waves) generated in bacterial cells by reaction-diffusion mechanism respond to the electric field is in particular an interesting topic to study for the emerging second instability induced by the coupling between the permitted wave mode and the electric field. Here, electrolytes are no longer spectators, but set up the electric field through the composition gradient of chemical waves. On the subject of chemical wave–electric field interactions, yet chemical systems such as BZ reaction present a strong resemblance to its counterparts in biological systems, direct applications of the theoretic results from BZ reaction (*52, 53*) to the bacterial analogue could be overshadowed by the mechanistic dissimilarity between them. First, all the reactions for BZ system are proceeded in the same ionic medium; while for some reacting systems in bacteria, reactions are carried out in two interfaced media: one is the cytoplasm, an ionic medium, and another is the cell membrane, a 2-dimensional liquid. Notably, the difference of diffusion coefficient for proteins in these two media is up to 3-4 orders. Moreover, the size of bacteria is more than 2 orders smaller than the characteristic dimension of BZ system and its varieties. Therefore, if a chemical wave/pattern with characteristic diffusion coefficient $\bar{D}$ could be self-organized only in the cytoplasm of bacterial cell, the diffusion length $(\bar{D}T_c)^{1/2}$ in an oscillation period $T_c$ of the chemical wave/pattern has to be much shorter than the characteristic length $\bar{L}$ of the phenomena of interest to enable instabilities. Otherwise, the reactant gradient could not be formed in bacterial cells due to fast diffusion of reactants in the system because the diffusion coefficient for reactants in bacterial cytoplasm is in the order of few to tens of $\mu m^2/s$ and bacterial size is in the order of few $\mu m$ such that $\left(\frac{\bar{D}T_c}{\bar{L}^2}\right) \gg 1$ for the chemical instabilities with period $T_c$ of few to tens of seconds. Alternatively, the reduction of characteristic diffusion coefficient $\bar{D}$ to the order of $0.01\,\mu m^2/s$ or smaller would suffice to form the composition gradient of chemical instabilities due to $\left(\frac{\bar{D}T_c}{\bar{L}^2}\right) \ll 1$, that is, some reactions must be carried out on the cell membrane and as such, the apparent diffusion coefficient of chemical instabilities shall be 3-4 orders smaller in comparison with that in bacterial cytoplasm.

#### The Phase Equation for Chemical Wave – Electric Field Interactions (Ortoleva's Method)

To get a better idea how a chemical wave interacts with the cross-gradient electric field in a bacterial cell as $\left(\frac{\bar{D}T_c}{\bar{L}^2}\right) \ll 1$, the dynamics of chemical wave–electric field interactions can be described by the coupling of the reaction-diffusion equations and Poisson's equation, that is, for an N-reactant species system

$$\frac{\partial \mathbb{C}_i}{\partial t} = -\nabla \cdot \vec{\boldsymbol{J}}_i + R_i(\mathbb{C}) \tag{7}$$

$$\vec{\boldsymbol{J}}_i = -D_i \nabla \mathbb{C}_i - M_i \mathbb{C}_i \nabla \Phi \tag{8}$$

$$\nabla^2 \Phi + \frac{4\pi F}{\epsilon_s \epsilon_0} \sum_{i=1}^{N} z_i \mathbb{C}_i = 0 \tag{9}$$

where $\mathbb{C}_i$ is the concentration for reactant species $i$; $\vec{\boldsymbol{J}}_i$ is the flux; and $R_i(\mathbb{C})$ is the net rate of reaction owing to all processes affecting species $i$. $D_i$ and $z_i$ are the $i$-diffusion coefficient and valence; and $M_i$ is $z_i F$ times the mobility of species $i$. $R_i$ may depend on all concentrations $\mathbb{C}_i = \{\mathbb{C}_1, \mathbb{C}_2, \cdots, \mathbb{C}_N\}$. The electric potential $\Phi$ is built up due to imbalanced charge distribution and traversed across a bacterial cell. Here, we simplify the analysis to a 1-dimensional system and, for plane waves of velocity $\mathfrak{v}$ traveling in the $x$-axis, Equations (7)-(9) can be further rearranged into the equations of motion in the wave-fixed coordinate system $\psi = x - \mathfrak{v}t$ (*53*). Then a closed integro-differential is obtained for steady electro-chemical wave propagation by dropping the subscripts and yields

$$\frac{d}{d\psi}\left[\boldsymbol{D}\frac{d\mathbb{C}}{d\psi} - \left(E_0 + \frac{4\pi F}{\epsilon_s \epsilon_0}\int_{\psi_0}^{\psi} d\psi\, \boldsymbol{z} * \mathbb{C}\right)\boldsymbol{M}\mathbb{C}\right] + \mathfrak{v}\frac{d\mathbb{C}}{d\psi} + \boldsymbol{R}(\mathbb{C}) = 0 \tag{10}$$

where $\vec{\boldsymbol{E}} = -\nabla\Phi$ is the electric field and let $E_0$ be $E(\psi_0)$ for a reference point $\psi_0$ and $\boldsymbol{z} * \mathbb{C} = \sum_{i=1}^{N} z_i \mathbb{C}_i$. Provided that $(\epsilon_s \epsilon_0 RT / 4\pi F^2 \bar{\mathbb{C}} \bar{\psi}^2)^{1/2} \ll 1$, i.e., the Debye length due to reactants is much smaller than the characteristic reaction-diffusion length $\bar{\psi}$, further approximation (*53*) of (10) by perturbation analysis leads to

$$\frac{d}{d\psi}\left[\boldsymbol{D}\frac{d\mathbb{C}}{d\psi}-\tilde{E}\boldsymbol{M}\mathbb{C}\right]+\mathfrak{v}\frac{d\mathbb{C}}{d\psi}+\boldsymbol{R}(\mathbb{C})=0 \tag{11}$$

$$\tilde{E}=\frac{I+\boldsymbol{z}\cdot\boldsymbol{D}\frac{d\mathbb{C}}{d\psi}}{\boldsymbol{z}\cdot\boldsymbol{M}\mathbb{C}} \tag{12}$$

where $\tilde{E}$ contains the Planck electric field due to the charge density $\boldsymbol{z}\cdot\mathbb{C}$ arising from the concentration gradient and the ohmic term $I/\boldsymbol{z}\cdot\boldsymbol{M}\mathbb{C}$. Specifically, a Planck potential is induced to neutralize the charge separation caused by the instability of reaction-diffusion mechanism and propagates along with an electrochemical wave. When the through current $I$ that flows across the system is imposed externally or by the buildup of electric potential in a bacterial cell, the electric field is directly coupled with the chemical instability by (12).

The spatiotemporally oscillatory pattern in bacteria is primarily generated by electrochemical instability described by Equations (11) and (12). Aside from subcellular waves or patterns, the instability may emerge by destabilization of phase wave either through phase diffusion or the coupling of wave mode and electric field. Let $\Psi(t)$ be the limit-cycle solution to this system, the trajectory $\mathbb{C}\approx\Psi(t+\phi)$ moves around the limit cycle, i.e., $d\Psi(t)/dt=R(\Psi)$, wherein the effect of diffusion gradually diminishes in the configuration $\mathbb{C}$-space as $\left(\frac{\bar{D}T_c}{\bar{L}}\right)\ll 1$ and the local phase $\phi$ of oscillation varies slowly in space and time. With the aid of multiple-scale analysis (*52*), the perturbation over the local phase about a limit cycle solution of (11) gives the phase equation

$$\frac{\partial\phi}{\partial t}=D_p\nabla^2\phi+\Delta|\nabla\phi|^2-\mu\vec{\boldsymbol{E}}\cdot\nabla\phi \tag{13}$$

$$\begin{Bmatrix}D_p\\ \Delta\end{Bmatrix}=\int_0^1\mathbf{f}\boldsymbol{D}\begin{Bmatrix}1\\ \Omega\end{Bmatrix}\frac{\partial\Psi}{\partial t_0}dt_0\,,\ \text{for}\int_0^1\mathbf{f}\frac{\partial\Psi}{\partial t_0}dt_0\equiv 1 \tag{14}$$

$$\mu\equiv\int_0^1\mathbf{f}\boldsymbol{M}\frac{\partial\Psi}{\partial t_0}dt_0 \tag{15}$$

where $D_p$ is the diffusion coefficient for phase $\phi$; $\Delta$ is the frequency renormalization factor ascribed to diffusion and $\Omega = \partial R(\mathbb{C})/\partial\mathbb{C}$ in relation to the reactions; $\mathbf{f}$ is the adjoint function to $d\Psi/dt$ and satisfies $\int_0^1 \mathbf{f}\frac{\partial\Psi}{\partial t_0}dt_0 \equiv 1$ for the linearized dynamics about the limit cycle. Similar to $D_p$ denoting phase diffusion, the parameter $\mu$ relates mobility $\boldsymbol{M}$ of charge-carrying reactants via (15) and denotes an electric mobility of the phase.

For periodic plane waves along the $x$-axis, upon responding to an electric field $\vec{\boldsymbol{E}}$ across the chemical gradient, then the phase shows

$$\phi = \vec{\boldsymbol{k}} \cdot \vec{\boldsymbol{x}} + \nu t, \tag{16}$$

$$\nu = \Delta k^2 - \mu\vec{\boldsymbol{E}} \cdot \vec{\boldsymbol{k}}. \tag{17}$$

Therefore, for frequency $\omega_c$ of the limit cycle, the frequency $\omega(k)$ of chemical waves/patterns with wavelength $2\pi/k$ in the length scale of cell length $L$, is obtained by $\omega_c\big(1+\nu(k)\big)$. Accordingly, the dispersion relation, when $k$ is small due to its equivalence to $\pi$ over the inverse of cell length, is given by

$$\omega(k) \sim \omega_c\big(1 - \mu\vec{\boldsymbol{E}} \cdot \vec{\boldsymbol{k}} + \Delta k^2\big) \tag{18}$$

Remarkably, the shift of the frequency for small $k$ wave is proportional to the inner product of wave vector $\vec{\boldsymbol{k}}$ and the electric field $\vec{\boldsymbol{E}}$, or the wavenumber $k$ and the strength of the field, as well as the crossing angle between $\vec{\boldsymbol{k}}$ and $\vec{\boldsymbol{E}}$. For the systems with field-free condition or the perpendicular direction of wave propagation relative to the cross-gradient field, the shift of phase diffusion is attributed to the parameter $\Delta$, that is, the effect arising from reactant diffusion and the rate of reactions. It is thereby indicated by the quadratic term of the dispersion relation (18) that rapid diffusion and/or high reaction rates would promote local instability of the phase, furthering the shift of frequency.

#### Diffusive Wave Mode Caused by Nucleotide Exchange in the Min-Protein Waves

As indicated above that apparent diffusion coefficient $\bar{D}$ must be in the order of $0.01\ \mu m^2/s$ and satisfies $\left(\frac{\bar{D}T_c}{\bar{L}}\right) \ll 1$ to enable instability in micron-sized bacterial cells, chemical waves/patterns emerging in such a cellular dimension require membrane-associated reactions. One such an example in *E. coli* is Min-protein oscillator (*54-57*), which is characterized by the pole-to-pole oscillations of MinD and MinE proteins. Both MinD and MinE possess cytoplasmic free-diffusing state and membrane-associated states. The difference in diffusion coefficient for these two states is up to 3-4 orders. Membrane-associated MinD is in an ATP-bound "active" state that can be activated by membrane-associated MinE to switch to an "inactive", free-diffusing ADP-bound state in the cytoplasm once they form MinDE complex for ATP hydrolysis (Fig. S12) (*54, 58*). Membrane-associated MinD can also induce conformational switches in MinE from free-diffusing "latent" state to amphipathic "active" state for membrane association and MinDE complexation (*59*). Nucleotide exchange between free-diffusing ADP-bound and ATP-bound MinD is the only reaction proceeding and influenced by the electric field self-sustained in the cytoplasm; while other reactions such as membrane associations, complexation and dissociations are carried out on the cell membrane (*1, 60*), wherein no membrane-associated reactions will experience any difference in electric potential. We will investigate and verify the plane-wave approximation to the dispersion relation (18) owing to chemical wave–electric field interaction by experiments using Min-protein oscillator in *E. coli* and its mutants.

The values for parameters $\mu$, $\bar{E}$ and $\Delta$ in the dispersion relation (15) are missing due to the lack of experimental measurement or simulation inference. First, in the prior section, we have discussed that the reactants as charge carriers cross the whole cell and experience nonuniform wave–field interactions. Though field strength varies over the whole cell, but for chemical wave traversing through the cell, we may expect the reactants interact with a mean electric field $\bar{E}$ over an oscillation period. Thereby, for L = 3 μm, we may use the estimated value at $|\rho_{nc}|/|\rho_{C^+}| \approx \frac{1}{16}$ , i.e., 13.82 $mV/\mu m$ as the expected magnitude of electric field $\bar{E}$ to represent the averaged field strength that reactants constantly interact when traversing across the cell over an oscillation period.

Next, the frequency normalization factor $\Delta$ relates to the diffusion coefficient and the reaction terms, or $\Omega = \partial R(\mathbb{C})/\partial\mathbb{C}$ (see (14)) . Because the electric field is built up in the bounded cellular space, the membrane-associated reactions do not interact with the field. The only reaction undergoing in the cytoplasm is the nucleotide exchange between free-diffusing MinD-ADP and MinD-ATP, which is the first order reaction with the kinetic constant $\sigma_{ADP\to ATP} \sim 1 - 6/\mathrm{s}$ (*61*). Since the diffusion coefficient for all membrane-associated species are 3-4 order smaller compared to free-diffusing species, the products for these membrane-associated species contribute very little to $\boldsymbol{D}\,\partial R(\mathbb{C})/\partial\mathbb{C}$ unless the reactions are concentration-dependent. However, the range of Min-protein concentration is within 2-order difference in accordance with our experimental results (Fig. S13). One may consider the reaction linked to free-diffusing species in computation of the frequency normalization factor $\Delta$, that is, $\Omega = \partial R(\mathbb{C})/\partial\mathbb{C} = \sigma_{ADP\to ATP}$. The diffusion coefficient for free-diffusing MinD has been measured by fluorescence correlation spectroscopy and reported about 16-17 $\mu m^2/s$ (*62*). Based on these data and $T_c = 59.01\,s$ extracted from the mean value of our experimental results (Fig. S13), the renormalization factor $\Delta$ is calculated to be 1.73 when $\sigma_{ADP\to ATP} = 6/s$ is adopted.

At last, we need to figure out the electric mobility of free-diffusing MinD over an oscillation period. Unfortunately, it is unlikely to acquire the electric mobility directly from experiments using living cells, but it's acknowledged that about 70% / 38% of all proteins in living systems have isoelectric points (pI) below pH 7 / pH 6, respectively (*63*). Accordingly, the proteins of interest here may be regarded to carry a negative net charge at physiological pH. General practices such as protein electrophoresis (*64*) and capillary zone electrophoresis (*65*) might indirectly provide some reference values that could be applied to the calculation. Especially capillary zone electrophoresis is extensively applied to proteomics and large amounts of data from a plethora of proteins are thereby measured. Since the concentration of electrolytes in bacteria is in the range of few hundreds mM, the electric mobilities of proteomic samples measured under the condition of high ionic strength (*65*) meet the requirement for protein transport under the intracellular electric field $(10^3 - 10^4\, V/m)$ in bacteria. From the data reported in the proteomic study by capillary zone electrophoresis (*65*), the distribution of electric mobilities cover the ranges between 5.5 – 26.5 $\mu m^2 \cdot mV^{-1} \cdot s^{-1}$. We postulate that the electric mobilities of all the proteins in the proteomic sample follows the normal distribution and in according to empirical (or 68–95–99.7) rule of normal distribution, $\pm 3$ standard deviations shall cover 99.7% of data points drawn from a normal distribution. We may thereby obtain the mean of electric mobility as 16 $\mu m^2 \cdot mV^{-1} \cdot s^{-1}$ and the standard deviation is ca. 3.5 $\mu m^2 \cdot mV^{-1} \cdot s^{-1}$. Inserting these values to (15), we may obtain $\bar{\mu} = 0.271 \pm 0.059\, \mu m^2 \cdot mV^{-1}$.

#### D. Dispersion Relation for Min-protein Waves in a 2D Rectangular Cell

The Min-protein oscillator comprises MinC, MinD and MinE proteins. MinC is considered as a passive effector distributed by membrane-associated MinD. Under the scheme of activator-inhibitor model in the framework of reaction-diffusion dynamics, membrane-associated MinD and MinE are thought as activator and inhibitor respectively. The interaction network of Min-protein oscillator illustrated in Fig. S12 is facilitated by the biochemical reactions between cytoplasmic MinD / MinE and membrane-associated MinD and MinDE. Briefly, MinD becomes an active ATPase as its binding to ATP and thereby presents hydrophobicity due to the conformation change from MinD-ADP to MinD-ATP. Once MinD-ATP associates on the membrane (step 1 in Fig. S12), it actively and cooperatively acquires more MinD-ATP in the cytoplasm to be membrane-associated (step 2). Membrane-associated MinD also recruits cytoplasmic MinE to associate with the membrane and forms MinDE complex. The complexation of MinD-ATP and MinE in turn triggers ATP hydrolysis in MinD-ATP (step 3) and inhibits their membrane-associated state owing to the hydrophilicity of freshly formed MinD-ADP after ATP hydrolysis, and as such to relieve the membrane-associated state of MinD-ADP and MinE , allowing them to enter a free-diffusing state in the cytoplasm (step 4). MinD-ADP may resume MinD-ATP through the reaction of nucleotide exchange during their diffusion in the cytoplasm (step 5). The collective cycling over the reaction-and-diffusion of MinD and MinE proteins switching between the membrane-associated and free-diffusing states accomplish the self-emergence of Min-protein waves.

To comprehend the insights of dispersion relation for Min-protein waves traversing across a bacterial cell, one need a set of reaction-diffusion (RD) equations to model the biochemical reactions illustrated in Fig. S12 for simplicity but without loss of generality. The minimal model that faithfully recapitulates the oscillatory dynamics is adopted to implement the derivation of the dispersion relation for Min-protein waves in a rectangular cell. Instead of dealing with a rod-like geometry, theoretical analysis upon a rectangular geometry is much easier to handle in mathematics. Since Min-protein oscillations have been observed in the rectangular cells that are resculpted by microfluidic chambers to demonstrate the resemblant wave dynamics displayed in rod-like cells (*66*), the qualitative results based on the analytic studies delved into a rectangular cell would reflect the most physical insights revealed in rod-like cells. Further, due to the longitudinal symmetry in a rod-like and rectangular cell, Min-protein waves are modeled in a 2D rectangular cell enclosed by the reactive membrane and assumed that MinD and MinE proteins are allowed to undergo free diffusion in the enclosed space and interact with each other when they associate with the reactive boundary. For cytoplasmic MinD-ATP, MinD-ADP, MinE and membrane-associated MinD and MinDE, the concentration and the surface density in a cell are $C_{D^*}$, $C_D$ , $C_E$ and $\hat{C}_d$ and $\hat{C}_{de}$, respectively. The total concentration of MinD and MinE are denoted as $C_D^0$ and $C_E^0$. The average expression level of total MinD concentration from in vivo experiments is denoted as $C_0$ . Then for any expression level of MinD and MinE in a cell, the concentration ratios of MinD over the average expression level of MinD and that of MinD over MinE are set to be $\alpha = C_D^0/C_0$ and $\gamma_0 = C_E^0/C_D^0$ in a cell. Therefore, owing to the law of mass conservation, the total amounts of MinD and MinE in a cell are given by

$$\begin{aligned}\int_\Omega C_D^0\, d\Omega &= \alpha \int_\Omega C_0 \; d\Omega \\ &= \int_\Omega C_{D^*}\, d\Omega + \int_\Omega C_D \; d\Omega + \int_{\partial\Omega} C_d \; d\partial\Omega + \int_{\partial\Omega} C_{de}\, d\partial\Omega\,,\end{aligned} \tag{1}$$

$$\int_\Omega C_E^0\, d\Omega = \int_\Omega C_E \; d\Omega + \int_{\partial\Omega} C_{de}\, d\partial\Omega. \tag{2}$$

That is,

$$\frac{C_D^0}{C_0} = \alpha = \frac{\left(\frac{\int_\Omega C_D^0 \, d\Omega}{\Omega}\right)}{\left(\frac{\int_\Omega C_0 \, d\Omega}{\Omega}\right)}$$

$$= \frac{\left(\frac{\int_\Omega C_{D^*} \, d\Omega}{\Omega}\right)}{\left(\frac{\int_\Omega C_0 \, d\Omega}{\Omega}\right)} + \frac{\left(\frac{\int_\Omega C_D \, d\Omega}{\Omega}\right)}{\left(\frac{\int_\Omega C_0 \, d\Omega}{\Omega}\right)} + \frac{\left(\frac{\Omega'}{\Omega}\right)\left(\frac{\int_{\partial\Omega} C_d \, d\partial\Omega}{\Omega'}\right)}{\left(\frac{\int_\Omega C_0 \, d\Omega}{\Omega}\right)}$$ (3)

$$+ \frac{\left(\frac{\Omega'}{\Omega}\right)\left(\frac{\int_{\partial\Omega} C_{de} \, d\partial\Omega}{\Omega'}\right)}{\left(\frac{\int_\Omega C_0 \, d\Omega}{\Omega}\right)},$$

and

$$\frac{C_E^0}{C_0} = \gamma = \alpha \cdot \gamma_0 = \frac{\left(\frac{\int_\Omega C_E^0 \, d\Omega}{\Omega}\right)}{\left(\frac{\int_\Omega C_0 \, d\Omega}{\Omega}\right)} = \frac{\left(\frac{\int_\Omega C_E \, d\Omega}{\Omega}\right)}{\left(\frac{\int_\Omega C_0 \, d\Omega}{\Omega}\right)} + \frac{\left(\frac{\Omega'}{\Omega}\right)\left(\frac{\int_{\partial\Omega} C_{de} \, d\partial\Omega}{\Omega'}\right)}{\left(\frac{\int_\Omega C_0 \, d\Omega}{\Omega}\right)},$$ (4)

where $\Omega$ is the volume of the cell and $\Omega' = \partial\Omega \cdot \xi$ is the volume of the cell membrane in the associated area $\partial\Omega$ and thickness $\xi$. The surface density $\hat{C}_d$ and $\hat{C}_{de}$ of membrane-associated MinD and MinDE follow the relations:

$$C_d = \frac{\int_{\partial\Omega} C_d \, d\partial\Omega}{\Omega'} = \frac{\int_{\partial\Omega} C_d \, d\partial\Omega}{\partial\Omega \cdot \xi} = \frac{\hat{C}_d}{\xi},$$ (5)

and

$$C_{de} = \frac{\int_{\partial\Omega} C_{de} \, d\partial\Omega}{\Omega'} = \frac{\int_{\partial\Omega} C_{de} \, d\partial\Omega}{\partial\Omega \cdot \xi} = \frac{\hat{C}_{de}}{\xi}$$ (6)

The skeleton model offers a minimal RD framework for describing the Min-protein waves, by which the reaction processes in both the cytoplasm and the cell membrane are integrated into the following PDE set. Besides, intracellular electric field (E-field) is introduced in the 1st order derivative, or advection term, in the partial differential equations to incorporate the Min-protein dynamics in the cytoplasm, constituting the framework of the reaction-diffusion system under the action of advection induced by E-field, or the RDA system. Namely, the dynamics of MinD-ATP, MinD-ADP and MinE in the cytoplasm is governed by

$$\frac{\partial C_{D^*}}{\partial t} = D_{D^*}\nabla^2 C_{D^*} - \vec{\boldsymbol{E}} \cdot M\nabla C_{D^*} + \sigma^* C_D \,, \tag{7.1}$$

$$\frac{\partial C_D}{\partial t} = D_D\nabla^2 C_D - \vec{\boldsymbol{E}} \cdot M\nabla C_D - \sigma^* C_D \,, \tag{7.2}$$

$$\frac{\partial C_E}{\partial t} = D_E\nabla^2 C_E - \vec{\boldsymbol{E}} \cdot M\nabla C_E \,, \tag{7.3}$$

with the boundary conditions

$$D_{D^*}\nabla_n C_{D^*} = -\left(\sigma_D C_{D^*} + \sigma_{dD}\hat{C}_d C_{D^*}\right) = -\left(\sigma_D + \sigma_{dD}\hat{C}_d\right)C_{D^*} \,, \tag{7.4}$$

$$D_D\nabla_n C_D = \sigma_{de}\hat{C}_{de}, \tag{7.5}$$

$$D_E\nabla_n C_E = \sigma_{de}\hat{C}_{de} - \sigma_{dE}\hat{C}_d C_E \,, \tag{7.6}$$

where $\nabla_n$ denotes the gradient outwardly normal to the interface, or the membrane, and the – and + signs denote the flux onto and off the interface. $D_{D^*}$, $D_D$ and $D_E$ are the diffusion coefficients for cytoplasmic MinD-ATP, MinD-ADP and MinE. $M$ is the electric mobility. $\sigma^*, \sigma_D$, $\sigma_{dD}$, $\sigma_{de}$ and $\sigma_{dE}$ are rate constants for the reaction terms.

While upon the interfaces (or the cell membrane), the dynamics of MinD and MinDE is governed by

$$\frac{\partial \hat{C}_d}{\partial t} = D_d \nabla^2 \hat{C}_d + \left(\sigma_D + \sigma_{dD} \hat{C}_d\right) C_{D^*} - \sigma_{dE} \hat{C}_d C_E, \tag{8.1}$$

$$\frac{\partial \hat{C}_{de}}{\partial t} = D_{de} \nabla^2 \hat{C}_{de} + \sigma_{dE} \hat{C}_d C_E - \sigma_{de} \hat{C}_{de}\,, \tag{8.2}$$

where $D_d$ and $D_{de}$ are the diffusion coefficients for membrane-associated MinD and MinE within the interface.

Since the rod-like geometry of bacteria hampers the analytic solutions for the RDA system (7.1-5) and (8.1-2), to understand the effect of intracellular E-field upon the Min-protein waves, a 2D rectangular geometry is instead applied to solve the dynamics of cytoplasmic MinD; and linear stability analysis is employed to derive the dispersion relation for characterizing the permitted wave modes influenced by intracellular E-field. Under the circumstance without interactions of Min proteins between the cytoplasm and the membrane, the behavior of MinE in the cytoplasm without intracellular E-field (7.3) is solely dominated by diffusion; while beyond that, the nucleotide exchange between MinD-ADP and MinD-ATP is the major reaction process to build up the density profile of MinD-ATP and MinD-ADP. *Halatek J.* and *Fery E.* provided the solution to MinD-ATP, MinD-ADP and MinE in the bulk when only the bottom side of a 2D rectangular geometry is considered as the reactive boundary condition (*67*). Consequently, the buildup of stationary density profiles for MinD-ATP, MinD-ADP and MinE is found distributed along the direction normal to the reactive membrane. Likewise, assuming that the dimension of a 2D rectangular cell is L x H (4μm x 1μm in x-z coordinates), the solution of Equations (7.1-6) without intracellular E-field can be decoupled to the functions in x and z coordinates respectively. Similar stationary density profiles for MinD-ATP, MinD-ADP and MinE can be found by considering 2D rectangular geometry with reactive boundary condition over four peripheral sides. The stationary density profiles in the cytoplasm are then given as

$$\tilde{c}_D(z) = \tilde{c}_D^*(\tilde{z}) \frac{cosh((2z - H)/2\ell)}{cosh(H/2\ell)}, \tag{9.1}$$

$$\tilde{c}_{D^*}(z) = \tilde{c}^*_{D^*}(\tilde{z}) + \tilde{c}^*_D(\tilde{z})\left(1 - \frac{cosh\big(\big((2z-H)\big)/2\ell\big)}{cosh(H/2\ell)}\right), \tag{9.2}$$

$$\tilde{c}_E(z) = \tilde{c}^*_E\,, \tag{9.3}$$

where $\tilde{z} = 0\ or\ H$ and $c_i^*$ denote the homogeneous stationary cytoplasmic densities at the membrane in z-direction and $\ell = \sqrt{D_{D^*}/\sigma^*}$ is the penetration depth into the cytoplasm. 2z and $2\ell$ in hyper cosine function reflect the symmetry of density profiles in z-direction due to symmetric boundary conditions. One may apply the stationary density profiles (9.1-3) to the reactive boundary conditions (7.4-6) and obtain

$$\ell \cdot \tilde{c}^*_D(\tilde{z}) \cdot tanh(H/2\ell) = \frac{\sigma_{de}\hat{C}_{de}}{\sigma^*}, \tag{10.1}$$

$$\ell \cdot \tilde{c}^*_D(\tilde{z}) \cdot tanh(H/2\ell) = \frac{\big(\sigma_D + \sigma_{dD}\hat{C}_d\big)\tilde{c}^*_{D^*}(\tilde{z})}{\sigma^*}, \tag{10.2}$$

$$0 = \sigma_{de}\hat{C}_{de} - \sigma_{dE}\hat{C}_d\tilde{c}^*_E(\tilde{z})\,, \tag{10.3}$$

On account of the symmetry in the reactive membranes along x-direction, the density profiles of MinD-ADP, MinD-ATP and MinE in the absence of intracellular E-field deem to behave oscillatory and encompass a linear combination of assorted cosine functions in wave number k. Whereas, the advection induced by intracellular E-field would break the symmetry in wave dynamics along x-direction. In real cellular context, one direct effect of intracellular E-field ascribing to the extended diffuse layer caused by imbalanced amounts of free-moving cations and anions in the cytosol, is creation of the pH gradient away from the cell membrane because hydrogen/hydronium ions are also monoatomic/small-molecular cations. The intracellular E-field, together with pH gradient, gives rise to the natural phenomenon, known as isoelectric focusing (*68-71*), where intracellular proteins, as amphoteric molecules, migrate within an internal pH gradient under intracellular E-field until they reach their specific isoelectric point (pI), where their net charge is zero, effectively concentrating them and aiding cellular organization, movement, and function, much like the lab isoelectric focusing separates proteins in a gel. Therefore, direct transport of cytosolic proteins owing to the advection induced by intracellular E-field is prone to reach equilibrium state in the wake of isoelectric focusing. One may consider that, for any concentration disturbance of interest around the equilibrium state in real cellular context, the action of intracellular E-field induced advection upon the concentration disturbance would become purely imaginary and lead to oscillatory, coupled behaviors, i.e., instability or sustained waves, in ionic solutions, where the real part of advection is almost damped.

To apprehend how Min-protein waves respond to the advection term in linear stability analysis, one may remove the advection term in (7.1-3) and as such to incorporate the action of intracellular E-field upon concentration gradient in the following mode analysis. Let $C_D\,(x,z,t) = e^{\lambda t}\hat{C}_D\,(x,z)$ and $C_{D^*}(x,z,t) = e^{\lambda t}\hat{C}_{D^*}(x,z)$, as well as $D_D = D_{D^*} = \mathcal{D}$. Equations (7.1-2) lead to

$$\frac{\partial \hat{C}_D}{\partial t} = \mathcal{D}\nabla^2\hat{C}_D - i\beta\left(\frac{\partial \hat{C}_D}{\partial x}\right) - \sigma^*\hat{C}_D\,, \tag{11.1}$$

$$\frac{\partial \hat{C}_{D^*}}{\partial t} = \mathcal{D}\nabla^2\hat{C}_{D^*} - i\beta\left(\frac{\partial \hat{C}_{D^*}}{\partial x}\right) + \sigma^*\hat{C}_D, \tag{11.2}$$

where $\langle \vec{\boldsymbol{E}}\cdot M\nabla\rangle := \langle i\vec{\beta}\cdot\nabla\rangle$, with $\beta = \langle\vec{\beta}\cdot\hat{x}\rangle$; and $\langle\cdot\rangle$ refers to spatial average, and $\hat{x}$ is the unit vector along the gradient direction. Then Equation (11.1-2) give

$$\mathcal{D}\nabla^2\hat{C}_D - i\beta\frac{\partial \hat{C}_D}{\partial x} - (\lambda + \sigma^*\,)\hat{C}_D = 0, \tag{12.1}$$

$$\mathcal{D}\nabla^2\hat{C}_{D^*} - i\beta\frac{\partial \hat{C}_{D^*}}{\partial x} + \sigma^*\hat{C}_D - \lambda\hat{C}_{D^*} = 0\,. \tag{12.2}$$

Next, let $\hat{C}_D\,(x,z) = e^{i\left(\frac{\beta x}{2\mathcal{D}}\right)}\,\phi(x,z)$ and $\hat{C}_{D^*}(x,z) = ei^{\left(\frac{\beta x}{2\mathcal{D}}\right)}\,\psi(x,z)$. Then Equation (12.1) leads to

$$\left(\frac{\partial^2\phi}{\partial x^2}+\frac{\partial^2\phi}{\partial z^2}\right)+\left(\frac{\beta^2}{4\mathcal{D}^2}-\frac{\lambda+\sigma^*}{\mathcal{D}}\right)\phi=0 \quad \Rightarrow \quad \nabla^2\phi+k_\phi^2\phi=0, \tag{13}$$

where $k_\phi^2=\frac{\beta^2}{4\mathcal{D}^2}-\frac{\lambda+\sigma^*}{\mathcal{D}}$. Likewise, Equation (12.2) is given as

$$\nabla^2\psi+k_\psi^2\psi=-\frac{\sigma^*}{\mathcal{D}}\phi, \tag{14}$$

where $k_\psi^2=\frac{\beta^2}{4\mathcal{D}^2}-\frac{\lambda}{\mathcal{D}}$. By using separation of variables method, one defines $\phi(x,z)=\mathrm{X}_\phi(x)\cdot\mathrm{Z}_\phi(z)$ and finds

$$\frac{\mathrm{X}_\phi{}''}{\mathrm{X}_\phi}+\frac{\mathrm{Z}_\phi{}''}{\mathrm{Z}_\phi}=k_\phi^2 \quad \Rightarrow \quad \frac{\mathrm{X}_\phi{}''}{\mathrm{X}_\phi}-k_x^2=\frac{\mathrm{Z}_\phi{}''}{\mathrm{Z}_\phi}+k_z^2=k_{\phi_z}^2, \tag{14.1}$$

$$\mathrm{X}_\phi{}''-k_x^2\,\mathrm{X}_\phi=0, \tag{14.2}$$

$$\mathrm{Z}_\phi{}''+k_z^2\,\mathrm{Z}_\phi=0, \tag{14.3}$$

where $k_x^2+k_z^2=k_\phi^2$. One may have linear stability analysis proceeding by studying the time evolution of small perturbations around the stationary solutions, written as $\tilde{c}_i(x,z,t)=\tilde{c}_i^*(x,z)+\delta\tilde{c}_i(x,z,t)$, subject to the reactive boundary conditions (7.4–6). Also, similar to Equations (9.1-3) and (10.1-3), the ansatz for the perturbations are subsequently decomposed into Fourier modes:

$$\delta\tilde{c}_D(x,z,t)=e^{\Delta x-\Lambda t}\cdot\sum_k e^{\lambda(k)t}\cdot cos(kx)\cdot\left(\frac{cosh\big((2z-H)/2\ell_k(\sigma^*+\lambda(k))\big)}{cosh\big(H/2\ell_k(\sigma^*+\lambda(k))\big)}\right)\cdot\delta\tilde{c}_D^k, \tag{15.1}$$

$$\begin{aligned}\delta\tilde{c}_{D^*}(x,z,t)=e^{\Delta x-\Lambda t}\cdot\sum_k e^{\lambda(k)t}\cdot cos(kx)\\ \cdot\left[\left(\frac{cosh\big((2z-H)/2\ell_k(\lambda(k))\big)}{cosh\big(H/2\ell_k(\lambda(k))\big)}\right)\cdot\left(\delta\tilde{c}_{D^*}^k+\delta\tilde{c}_D^k\right)\right.\\ \left.-\left(\frac{cosh\big((2z-H)/2\ell_k(\sigma^*+\lambda(k))\big)}{cosh\big(L/2\ell_k(\sigma^*+\lambda(k))\big)}\right)\cdot\delta\tilde{c}_D^k\right],\end{aligned} \tag{15.2}$$

$$\delta\tilde{c}_E(x,z,t)=e^{\Delta x-\Lambda t}\cdot\sum_k e^{\lambda(k)t}\cdot\ cos(kx)\cdot\left(\frac{cosh\big((2z-H)/2\ell_k(\lambda(k))\big)}{cosh\big(H/2\ell_k(\lambda(k))\big)}\right)\cdot\delta\tilde{c}_E^k, \tag{15.3}$$

$$\delta\tilde{c}_d(x,z,t) = \sum_k e^{\lambda(k)t} \cdot \cos(kx) \cdot \delta\tilde{c}_d^k\,, \tag{15.4}$$

$$\delta\tilde{c}_{de}(x,z,t) = \sum_k e^{\lambda(k)t} \cdot \cos(kx) \cdot \delta\tilde{c}_{de}^k\,, \tag{15.5}$$

where $\Delta = \frac{i\beta}{2\mathcal{D}}$ and $\Lambda = \sigma^* - \frac{\beta^2}{4\mathcal{D}}$; and the generalized penetration depth with respect to $\tilde{c}_i$ is obtained by substituting the ansatz into Equations (7.1-3) and (8.1-2) and defined as

$$\ell_k(\chi) = \sqrt{\frac{D_i}{\chi + D_i k^2}}\,. \tag{16}$$

Equations (7.4-6) show that the coupling between the membrane concentrations and the cytoplasmic density profiles can be expressed as

$$\begin{aligned} \Lambda_k^{D_i}(\chi) &= \frac{D_i}{\ell_k(\chi)}\, tanh\left(\frac{H}{2\ell_k(\chi)}\right) \cong \frac{D_i \cdot H}{2\ell_k(\chi)^2} \\ &= \frac{H}{2} \cdot (\chi + D_i k^2). \qquad \text{(for small } k\text{)} \end{aligned} \tag{17}$$

Before preceding to linear stability analysis, Equations (7.1-6) and (8.1-2) casted into the analysis of dimensionless variables are useful in revealing fundamental relationships, generalizing solutions across scales, and identifying and highlighting key parameters that govern physical behavior. Therefore, a set of dimensionless (primed) quantities are introduced by defining characteristic values (indicated with a bar)

$$\begin{aligned} &C = \bar{C}\, C', \qquad D = \bar{D}\, D', \qquad M = \bar{M}\, M', \\ &x_j = \bar{x}\, x_j', \qquad t = \bar{t}\, t', \qquad R(C) = \bar{C}\, R'(C)/\bar{t}, \\ &\bar{x}^2 = \bar{D}\, \bar{t}, \qquad \bar{M} = F\bar{D}/R_g T, \end{aligned} \tag{18}$$

where subscript $j$ denotes $x$ and $z$ coordinate; $F$, $R_g$ and $T$ are Faraday's constant, gas constant and absolute temperature respectively. $R(C)$ denotes the reaction term in (7.1-6) and

(8.1-2) . The last equation connects $\bar{M}$ and $\bar{D}$ via Einstein's relation. Let $\bar{x}$ and $\bar{D}$ be the cell length L and $D_{D^*}$, then one may get $\bar{t} = L^2/D_{D^*}$. Further, also let $\epsilon = \Omega'/\Omega$ and

$$u \equiv \frac{C_{D^*}}{C_0} = \frac{\left(\frac{\int_\Omega C_{D^*}\, d\Omega}{\Omega}\right)}{\left(\frac{\int_\Omega C_0\, d\Omega}{\Omega}\right)}, \tag{19.1}$$

$$\epsilon v \equiv \epsilon \cdot \frac{C_d}{C_0} = \frac{\epsilon}{\xi} \cdot \frac{\hat{C}_d}{C_0} = \bar{v} = \epsilon \cdot \frac{\left(\frac{\int_{\partial\Omega} C_d\, d\partial\Omega}{\Omega'}\right)}{\left(\frac{\int_\Omega C_0\, d\Omega}{\Omega}\right)}, \tag{19.2}$$

$$\epsilon w \equiv \epsilon \cdot \frac{C_{de}}{C_0} = \frac{\epsilon}{\xi} \cdot \frac{\hat{C}_{de}}{C_0} = \bar{w} = \epsilon \cdot \frac{\left(\frac{\int_{\partial\Omega} C_{de}\, d\partial\Omega}{\Omega'}\right)}{\left(\frac{\int_\Omega C_0\, d\Omega}{\Omega}\right)}. \tag{19.3}$$

In according to Equation (1)-(4), we also have

$$\frac{C_D}{C_0} = \alpha - u - \bar{v} - \bar{w}\,, \tag{19.4}$$

$$\frac{C_E}{C_0} = \alpha\gamma_0 - \bar{w} = \gamma - \bar{w}\,. \tag{19.5}$$

By casting these dimensionless quantities and dropping the primes, Equations (7.1-6) and (8.1-2) are converted to

$$\frac{\partial u}{\partial t} = \nabla^2 u - \vec{\beta} \cdot \nabla \mathrm{u} + \left[\mathrm{a}\alpha - (\mathrm{a}+\mathrm{b})u - \mathrm{a}\bar{v} - \mathrm{a}\bar{w} - \mathrm{c}u\left(\frac{\xi}{\epsilon}\bar{v}\right)\right], \tag{20.1}$$

$$\frac{\partial \bar{v}}{\partial t} = \delta\nabla^2 \bar{v} - \vec{\beta} \cdot \nabla \bar{v} + \left[\left(\frac{\epsilon \mathrm{b}}{\xi}u\right) - \gamma\bar{v} + \mathrm{c}u\bar{v} + \mathrm{h}\bar{v}\bar{w}\right], \tag{20.2}$$

$$\frac{\partial \bar{w}}{\partial t} = \delta\nabla^2 \bar{w} - \vec{\beta} \cdot \nabla \bar{w} + [\gamma\bar{v} - \mathrm{h}\bar{v}\bar{w} - \mathrm{e}\bar{w}]\,, \tag{20.3}$$

$$\frac{\partial}{\partial t}(\gamma - \bar{w}) = d\nabla^2(\gamma - \bar{w}) - \vec{\beta} \cdot \nabla(\gamma - \bar{w}) + [-\gamma\bar{v} + \mathrm{h}\bar{v}\bar{w} + \mathrm{e}\bar{w}] \,, \tag{20.4}$$

$$\begin{aligned} \frac{\partial}{\partial t}(\alpha - u - \bar{v} - \bar{w}) \\ &= \nabla^2(\alpha - u - \bar{v} - \bar{w}) - \vec{\beta} \cdot \nabla(\alpha - u - \bar{v} - \bar{w}) \\ &+ \left[-\mathrm{a}\alpha + \mathrm{a}u + \mathrm{a}\bar{v} + \left(\mathrm{a} + \frac{\xi \mathrm{e}}{\epsilon}\right)\bar{w}\right]. \end{aligned} \tag{20.5}$$

The intracellular E-field is modulated by the charge density induced by the negatively-charged chromosomes and associated biomolecules in the nucleoid, and as such to bolster the sway of advection over charged biomolecules in wave dispersion. The parameters d and δ are dimensionless diffusion coefficients $D_D/D_{D^*}$ and $D_d/D_{D^*}$, where $D_D = D_{D^*}$ and $D_d = D_{de}$ are assumed. Also, the parameters a, b, c, e and h are set to stand for the rate constant $\sigma^*, \sigma_D$, $\sigma_{dD}$, $\sigma_{de}$ and $\sigma_{dE}$. Notably, owing to the mass conservation (19.4-5), the five-component RDA system in (7.1-3) and (8.1-2) is reduced to three-component RDA system (20.1-5). The reduction benefits the numerical solvation of homogeneous equilibrium states, where all time and spatial derivative terms are zero. .

To carry out linear stability analysis and derive dispersion relation, one may analyze how small perturbations from homogeneous steady-state solutions evolve over time by substituting the ansatz

$$(u, \bar{v}, \bar{w})^T = (u^*, \bar{v}^*, \bar{w}^*)^T + (\delta u, \delta\bar{v}, \delta\bar{w})^T e^{ikx+\lambda t} + c.c., \tag{21}$$

into Equations (17), where $k$ is the spatial wave number and λ is the temporal eigenvalue, respectively, as well as $(u^*, \bar{v}^*, \bar{w}^*)$ homogeneous equilibrium and $\|\delta u, \delta\bar{v}, \delta\bar{w}\| \ll 1$. Consequently, by solving the corresponding eigenvalue problem, the characteristic matrix $\mathbb{M}$ is typically expressed as

$$det[\mathbb{M}] = det\left[\mathbb{J}\left(R';\ (u^*, \bar{v}^*, \bar{w}^*)\right) + k\mathbb{L} - k^2\mathbb{D} - \lambda(k)\mathbb{I}\right] = det[\mathbb{J}_{\mathbf{0}} + \mathbb{J}_{\mathbf{1}}] = 0\,, \tag{22}$$

where $\mathbb{J}_{\mathbf{0}} = \mathbb{J}(R'(u^*, \bar{v}^*, \bar{w}^*))$ is the Jacobian matrix of the reactions $R'(C')$ at the homogeneous equilibrium $(u^*, \bar{v}^*, \bar{w}^*)$; $\mathbb{L} = (\beta/2L,\ 0,\ 0,\ \beta/2L,\ \beta/2L) \cdot \mathbb{I}$ is the first-order derivative term; $\mathbb{D} = (1,\ \delta,\ \delta,\ d,\ 1) \cdot \mathbb{I}$ is the diffusion matrix; and $\mathbb{I}$ is the identity matrix. The dispersion relation is then given by solving the characteristic equation (22). Here, one may incorporate the original Jacobian matrix $\mathbb{J}_{\mathbf{0}}$ derived from Equations (20.1-5) with the $\mathbb{J}_{\mathbf{1}}$ matrix, which includes the modification of Jacobian matrix due to the boundary conditions, the influence of the first-order advection term, the diffusion term and the eigenvalue term. That is,

$$\mathbb{J}_{\mathbf{0}} = \begin{bmatrix} -B & -\left(\frac{\xi \cdot c \cdot u^*}{\epsilon}\right) & 0 & 0 & \left(B + \frac{\xi \cdot c \cdot u^*}{\epsilon}\right) \\ \left(\frac{\epsilon \cdot b}{\xi} + c \cdot \bar{v}^*\right) & (-\Gamma + h \cdot \bar{v}^*) & h \cdot \bar{v}^* & -h \cdot \bar{v}^* & \left(-\frac{\epsilon \cdot b}{\xi} - c \cdot \bar{v}^* + \Gamma - c \cdot u^* - h \cdot \bar{v}^*\right) \\ 0 & \Gamma & -E & E & (-\Gamma + E) \\ 0 & -\Gamma & E & -E & (\Gamma - E) \\ 0 & 0 & e & -e & -e \end{bmatrix} \tag{23.1}$$

$$\mathbb{J}_1 = \begin{bmatrix} \left(-\frac{a \cdot H}{2L} + \frac{\beta \cdot k}{2L} - \Lambda_k^1(\lambda)\right) & -\left(\frac{a \cdot H}{2L}\right) & -\left(\frac{a \cdot H}{2L}\right) & \left(\frac{a \cdot H}{2L}\right) & \left(\frac{3a \cdot H}{2L}\right) \\ 0 & -\Lambda_k^\delta(\lambda) & 0 & 0 & 0 \\ 0 & 0 & -\Lambda_k^\delta(\lambda) & 0 & 0 \\ 0 & 0 & 0 & \left(\frac{\beta \cdot k}{2L} - \Lambda_k^d(\lambda)\right) & 0 \\ \left(\frac{a \cdot H}{2L}\right) & \left(\frac{a \cdot H}{2L}\right) & \left(\frac{a \cdot H}{2L}\right) & -\left(\frac{a \cdot H}{2L}\right) & \left(-2\left(\frac{a \cdot H}{2L}\right) + \frac{\beta \cdot k}{2L} - \Lambda_k^1(\lambda + a)\right) \end{bmatrix} \tag{23.2}$$

where $B = b + c \cdot \bar{v}^* \cdot \xi/\epsilon$; $\Gamma = \alpha * \gamma_0 - h \cdot \bar{w}^*$; $E = e + h \cdot \bar{\mathrm{v}}^*$; and $\Lambda_k^{D_i}(\lambda)$ is from equation (17). The characteristic equation (23) is then numerically computed and thereof the dispersion relation is obtained.

At last, on account of its real and/or imaginary part, the advection term, $\vec{\beta} \cdot \nabla$, induced by intracellular E-field plays a crucial role in determination of the wave modes. The advection term, which is proportional to $ik$, fundamentally contributes terms that are odd in $k$. The real advection typically relates to direct transport; while the imaginary advection contributes to the emergence of instability and wave modes. As the effect of isoelectric focusing in the cytosol incurs a balanced state of transport, the advection induced by intracellular E-field becomes purely imaginary. For a general linear stability analysis of the dynamic system, the growth/decay rate for a wave mode $e^{ikx+\lambda t}$ is contributed by the real part of $Re[\lambda(k)]$. The imaginary part $Im[\lambda(k)]$ is known to delineate the phase dynamics, leading to the determination of wave modes.

### E. Statistical Analysis

The datasets ef1xyt to ef4xyt in the demonstrated code represent the inner products of electrical field and wave vector (i.e. $\vec{E}\cdot\vec{k}$), calculated from anucleate, wildtype, chloramphenicol-treated and rifampicin-treated cells, respectively. These data are skewed and apparently not normally distributed such that parametric testes are not appropriate to use for verifying if the true location parameters of the populations are equal. Instead, non-parametric Mann-Whitney test is used to test significant difference between the *medians* from two datasets. Further, non-parametric Kruskal-Wallis test is used to test significant difference among the *medians* from all datasets. Finally, K-sample t-test is used to test significant difference among the *means* from all datasets. For those test results with p-value < 0.05 indicate the statistical significance of rejection in the null hypothesis $H_0$. The testes were run under Mathematica (Wolfram Research, Inc) and the functions/results are shown below.

```
In[267]:= MannWhitneyTest[{ef1xt, ef2xt}, Automatic, {"TestDataTable"}]
          MannWhitneyTest[{ef2xt, ef3xt}, Automatic, {"TestDataTable"}]
          MannWhitneyTest[{ef2xt, ef4xt}, Automatic, {"TestDataTable"}]
          MannWhitneyTest[{ef3xt, ef4xt}, Automatic, {"TestDataTable"}]
          ℋ = LocationEquivalenceTest[{ef1xt, ef2xt, ef3xt, ef4xt}, "HypothesisTestData"];
          ℋ["TestDataTable", {"KSampleT", "KruskalWallis"}]
```

Out[267]=

| | Statistic | P-Value |
|---|---|---|
| Mann-Whitney | $1.04453\times10^{6}$ | 0.000556266 |

Out[268]=

| | Statistic | P-Value |
|---|---|---|
| Mann-Whitney | $1.3163\times10^{6}$ | $1.31373\times10^{-15}$ |

Out[269]=

| | Statistic | P-Value |
|---|---|---|
| Mann-Whitney | $1.24954\times10^{6}$ | $2.19635\times10^{-7}$ |

Out[270]=

| | Statistic | P-Value |
|---|---|---|
| Mann-Whitney | $1.07866\times10^{6}$ | 0.0439253 |

Out[272]=

| | Statistic | P-Value |
|---|---|---|
| K-Sample T | 1.52244 | 0.206481 |
| Kruskal-Wallis | 60.8022 | $3.44376\times10^{-13}$ |

### F. Capacity-Limited Surface Reaction Kinetics

Heterogeneous reactions at solid–liquid and solid–gas interfaces are governed by constraints that do not arise in homogeneous bulk phases. In a homogeneous medium, the reaction rate generally changes continuously with reactant concentration and is often represented by integer- or fractional-order power laws. At an interface, in contrast, reaction is limited by a finite and discrete population of active surface sites, imposing a capacity constraint on the kinetics.

When the reactant concentration in the liquid phase is very low, most surface sites remain vacant and the overall rate is controlled primarily by delivery of reactant to the interface, producing the familiar first-order limit. As the concentration increases, site occupancy rises until the available sites become saturated; the rate then loses its dependence on bulk concentration and approaches zero-order behavior. Between these limiting cases, partial coverage produces a nonlinear response that can be expressed as an effective fractional kinetic order. This framework applies to direct unimolecular surface reactions as well as competitive and autocatalytic pathways. The following analysis develops the corresponding rate expressions and half-life relations across the first-order-to-zero-order transition.

#### Surface Capacity and the Origin of Kinetic-Order Transitions

A mathematical description of a capacity-limited surface begins by specifying how reactant molecules populate a finite set of adsorption sites. We adopt the Langmuir adsorption isotherm, based on three assumptions:

1. The solid contains a fixed total number of distinct active sites, denoted by S.
2. Each active site accommodates at most one adsorbed molecule, so adsorption forms a single molecular layer.
3. All sites have the same adsorption energy, and adsorbed molecules do not interact laterally with neighboring molecules.

Define $\theta_A$ as the fractional surface coverage of reactant A, and let $\theta_V$ denote the fraction of vacant sites. The site balance is therefore:

$$\theta_A + \theta_V = 1 \Rightarrow \theta_V = 1 - \theta_A \quad (1)$$

A molecule of A adsorbs when it encounters an unoccupied site S, whereas desorption corresponds to rupture of the bond between A and the surface:

$$A + S \rightleftharpoons A \cdot S \quad (2)$$

At adsorption equilibrium, the forward adsorption flux and the reverse desorption flux are equal:

$$k_a[A]\theta_V \ = \ k_d\theta_A \tag{3}$$

Inserting the site-balance relation into the equilibrium condition gives:

$$k_a[A](1 \ - \ \theta_A) \ = \ k_d\theta_A \tag{4}$$

Introducing $K_ς = k_a/k_d$ and collecting terms gives the standard Langmuir isotherm:

$$\theta_A \ = \ \frac{K_ς[A]}{1 \ + \ K_ς[A]} \tag{5}$$

**The Continuous Shift in Kinetic Order**

Within the Langmuir–Hinshelwood mechanism (*72-74*), the macroscopic reaction rate R is proportional to the fraction of surface sites occupied by the reacting species. The proportionality constant *k* represents the intrinsic rate constant of the surface reaction:

$$R \ = \ -\frac{d[A]}{dt} = \frac{kK_ς[A]}{1 \ + \ K_ς[A]} \tag{6}$$

The concentration-dependent kinetic order can be defined through the logarithmic sensitivity of the rate to the bulk reactant concentration:

$$n_{app} \ = \ \frac{d \ ln \ R}{d \ ln \ [A]} = \frac{[A]}{R} \cdot \frac{dR}{d[A]} \tag{7}$$

Differentiating the rate expression with respect to concentration using the quotient rule gives:

$$\frac{dR}{d[A]} \ = \ \frac{kK_ς}{(1 \ + \ K_A[A])^2} \tag{8}$$

Substitution into the definition above yields the resulting expression for the apparent kinetic order:

$$n_{app} \ = \ \frac{1}{1 \ + \ K_ς[A]} \tag{9}$$

#### Case I — Direct Binding with a Unimolecular Surface Reaction

Consider a direct unimolecular surface process in which an adsorbed reactant undergoes conversion to a liquid-phase product without competition from another adsorbate. The resulting kinetics follow a mixed-order differential equation:

$$-\frac{d[A]}{dt} = \frac{kK_\varsigma[A]}{1 + K_\varsigma[A]} \tag{10}$$

**The Integrated Rate Law**

To obtain the concentration as a function of time, separate the concentration-dependent and time-dependent terms:

$$\frac{1 + K_\varsigma[A]}{kK_\varsigma[A]} d[A] = -dt \tag{11}$$

Decomposing the concentration factor on the left-hand side gives:

$$(\frac{1}{kK_\varsigma} \cdot \frac{1}{[A]} + \frac{1}{k}) d[A] = -dt \tag{12}$$

Integrating from the initial state at $t = 0$ to the remaining concentration at time $t$ gives the integrated rate law:

$$t = \frac{1}{kK_\varsigma} ln\frac{[A]_0}{[A]_t} + \frac{1}{k}([A]_0 - [A]_t) \tag{13}$$

**Half-Life in the Intermediate Regime**

Applying the half-life condition, namely that the remaining reactant concentration equals one-half of its initial value, gives the explicit intermediate half-life:

$$t_{1/2} = \frac{\ln(2)}{kK_\varsigma} + \frac{[A]_0}{2k} \tag{14}$$

#### Case II — Competitive Autocatalysis on a Capacity-Limited Surface

We next consider a surface-bound autocatalytic pathway in which liquid-phase reactant is converted to product only in the presence of pre-existing product acting as a catalyst. Reactant and product compete for the same finite population of adsorption sites.

$$\theta_A = \frac{K_\varsigma[A]}{1 + K_\varsigma[A] + K_P[P]} \tag{15}$$

Because the surface reaction requires both adsorbed species, its rate is proportional to their respective surface coverages, leading to the competitive rate law:

$$-\frac{d[A]}{dt} = \frac{kK_\varsigma K_P[A][P]}{(1 + K_\varsigma[A] + K_P[P])^2} \tag{16}$$

Mass Conservation and Dimensionless Parameter Grouping

For a closed system, mass conservation gives $[P] = C - [A]$. For compact notation, define the parameters $\alpha = 1 + K_p C$, $\beta = K_\varsigma - K_p$, and $\gamma = k K_\varsigma K_p$. The governing differential equation then takes the form:

$$-\frac{d[A]}{dt} = \frac{\gamma[A](C - [A])}{(\alpha + \beta[A])^2} \tag{17}$$

Explicit Half-Life for the Autocatalytic Reaction

Term-by-term integration using partial fractions, followed by application of the half-life condition, produces the explicit half-life expression for the intermediate site-limited autocatalytic regime:

$$t_{1/2} = \frac{1}{kK_\varsigma K_P}[-\frac{1}{2}(K_\varsigma - K_P)^2[A]_0 + \frac{(1+K_P C)^2}{C} ln(2) + \frac{(1+K_\varsigma C)^2}{C} ln(1 + \frac{[A]_0}{2[P]_0})] \tag{18}$$

#### Asymptotic Behavior of the Autocatalytic Half-Life

To determine how the processing time changes with increasing initial reactant concentration, examine the asymptotic limit of the explicit half-life expression as the initial concentration becomes very large, $[A]_0 \rightarrow \infty$.

In this limit, the initial catalyst or product concentration becomes small compared with the reactant pool, $[P]_0 \ll [A]_0$, so the total concentration constant approaches the initial reactant concentration, $C \approx [A]_0$. The scaling of the individual terms can then be evaluated with respect to $[A]_0$:

Linear Contribution:

$$-\frac{1}{2}(K_\varsigma - K_P)^2[A]_0 \propto \mathcal{O}([A]_0) \tag{19}$$

Reactant Adsorption Contribution:

After replacing $C$ by $[A]_0$,

$$\frac{(1+K_P C)^2}{C} ln(2) \approx \frac{{K_P}^2{[A]_0}^2}{[A]_0} ln(2) \propto \mathcal{O}([A]_0) \tag{20}$$

Product Competitive-Adsorption Contribution:

Then replacing $C$ by $[A]_0$,

$$\frac{(1+\ K_{\varsigma}C)^2}{C}\ ln(1+\frac{[A]_0}{2[P]_0})\ \approx\ \frac{{K_P}^2{[A]_0}^2}{[A]_0}\ ln(\frac{[A]_0}{2[P]_0})\ \propto\ \mathcal{O}([A]_0\ ln([A]_0)) \tag{21}$$

Because $\mathcal{O}(x \cdot \ln(x))$ grows more rapidly than $\mathcal{O}(x)$ at large x, the former ultimately dominates, giving:

$$t_{1/2}\ \approx\ \frac{K_{\varsigma}}{kK_P}\ [A]_0\ ln([A]_0)\ \rightarrow\ \infty \tag{22}$$

This asymptotic result has an important physical implication: sufficiently large reactant concentrations ultimately increase the half-life because the finite surface becomes strongly site-poisoned. Product generated during the reaction accumulates on the interface faster than it can desorb, creating a bottleneck that eventually dominates over the accelerating effect expected from homogeneous autocatalysis.

At very low concentrations, a modest increase in reactant can initially reduce the half-life by facilitating autocatalytic product formation, consistent with the shallow minimum in the corresponding curve. This benefit does not persist, however. Once the surface approaches substantial occupancy, the finite site capacity becomes the controlling constraint.

As product accumulates, incomplete desorption leaves fewer sites available for incoming reactant, intensifying competition between the two species. The reaction therefore becomes increasingly self-limiting: the density-driven buildup of product produces surface poisoning that ultimately overwhelms the acceleration associated with increasing bulk reactant concentration.

#### Summary

Capacity-limited surfaces impose kinetic constraints that fundamentally distinguish interfacial reactions from homogeneous reactions. As reactant concentration increases, the system progresses through three regimes: a low-coverage first-order limit, an intermediate partially saturated regime with a concentration-dependent apparent order, and a high-coverage zero-order limit. For direct binding, saturation changes both the rate and the concentration dependence of the half-life. For the competitive autocatalytic pathway, increasing concentration can eventually lengthen the half-life because product accumulation and site competition create a surface bottleneck.

| **Kinetic feature** | **First-order (Low [A])** | **Mixed-order (Intermediate)** | **Zero-order (High [A])** |
|---|---|---|---|
| **Surface state** | Predominantly vacant | Partially saturated | Fully saturated |
| **Direct Binding Rate:** $-\frac{d[A]}{dt}$ | $\approx kK_{\varsigma}[A]$ | $\frac{kK_{\varsigma}[A]}{1 + K_{\varsigma}[A]}$ | $\approx k$ |
| **Direct Binding Half-Life:** $t_{1/2}$ | $\frac{\ln(2)}{kK_{\varsigma}}$ | $\frac{\ln(2)}{kK_{\varsigma}} + \frac{[A]_0}{2k}$ | $\frac{[A]_0}{2k}$ |
| **Autocatalytic half-life** | Shortens through autocatalytic acceleration | Lengthens through site competition | Increases through $\mathcal{O}([A]_0\, ln([A]_0)$ site poisoning |

### APPENDIX: Key Numbers and Parameters

**Compositions in an *E. coli* cell**

| Composition | Amount/Cell | Concentration | Copies/Cell | Remark |
|---|---|---|---|---|
| DNA | 0.017 pg | 11 -18 mg/ml (*75, 76*) | 5.75x10$^6$ bp | 4.6x10$^6$ bp (genome size) |
| RNA | 0.200 pg | 75 – 120 mg/ml (*75, 76*) | 3.382x10$^7$ nt | |
| Protein | 0.200 pg | 200 – 320 mg/ml (*75, 76*) | 2.35x10$^6$ copies | |

**Major nucleoid-associated proteins (NAPs) of *E. coli***

| Growth Phase | Stationary Phase |
|---|---|
| 153,000 copies (*8*) | 280,000 copies (*8*) |

**Compositions in the nucleoid of *E. coli***

| Composition | Abundance in a cell | Copies/Cell | Remark | |
|---|---|---|---|---|
| DNA | 80% (*9, 10*) | 4.6x10$^6$ bp | | |
| RNA | 10% (*9, 10*) | 3.382x10$^6$ nt | | |
| Protein | 10% (*9, 10*) | 2.35x10$^5$ copies | Growth phase | 1.66x10$^5$ copies |
| | | | Stationary Phase | 3.04x10$^5$ copies |

The protein abundance in the cytosol and the nucleoid of E. coli is about 66.7%(*77*). Since the protein abundance in the nucleoid is known to be 10% (*9, 10*), 56.7% of proteins reside in the cytosol.

**Charges carried by the nucleoid of *E. coli***

| Composition | Charge number | | Remark |
|---|---|---|---|
| DNA | 9.2x10$^6$ e$^-$ | | 2 e$^-$/bp |
| RNA | 3.382x10$^6$ e$^-$ | | 1 e$^-$/nt |
| Protein | Growth Phase | Stationary Phase | 20 e$^-$/protein<br>Average copy number = 2.35x10$^5$<br>NAPs$^{Stationary}$/ NAPs$^{Growth}$ = 1.83 (*8*) |
| | 2.49x10$^6$ e$^-$ | 4.56x10$^6$ e$^-$ | |
| Total | 15.90x10$^6$ e$^-$ | 18.66x10$^6$ e$^-$ | |

**Other parameters in *E. coli* cells**

| | | |
|---|---|---|
| $C_{K^+}^0$ | 231.25 mM | 1.392x10$^8$ #/μm$^3$; initial $K^+$ concentration |
| $C_{Cl^-}^0$ | 20.65 mM | 1.243x10$^7$ #/μm$^3$; initial $Cl^-$ concentration |
| $D_{K^+}$ | 1950 μm$^2$/s | Diffusivity of potassium ions |
| $D_{Na^+}$ | 1330 μm$^2$/s | Diffusivity of sodium ions |

| | | |
|---|---|---|
| $D_{Cl^-}$ | 2020 $\mu m^2/s$ | Diffusivity of chloride ions |
| $D_{X^-}$ | 2.56 $\mu m^2/s$ | Diffusivity of other anions |
| $\epsilon_0$ | 8.85 x$10^{-12}$ F/m ($C \cdot V^{-1} \cdot m^{-1}$) | Vacuum permittivity |
| $\epsilon_s$ | 78 | Relative permittivity |
| F | 96485 s·A/mol | Faraday constant |
| $\kappa$ | 1.21x$10^6$ /$\mu m^2$ | $\kappa^2 = \frac{F^2}{\epsilon_s \epsilon_0 RT} C_{K^+}^0 = \frac{1}{{\lambda_D}^2}$ <br> $\lambda_D$: Debye screening length |
| $\lambda_D$ | 0.909 nm | |
| NC ratio | 0.52 (*7*) | Nucleocytoplasmic ratio |

#### References


1. D. Fange, J. Elf, Noise-induced Min phenotypes in E. coli. *PLoS Comput Biol* **2**, e80 (2006).
2. Y. L. Shih, I. Kawagishi, L. Rothfield, The MreB and Min cytoskeletal-like systems play independent roles in prokaryotic polar differentiation. *Mol. Microbiol.* **58**, 917-928 (2005).
3. J. P. Shen, Y. R. Chang, C. F. Chou, Frequency modulation of the Min-protein oscillator by nucleoid-associated factors in Escherichia coli. *Biochem. Biophys. Res. Commun.* **525**, 857-862 (2020).
4. J. M. Guberman, A. Fay, J. Dworkin, N. S. Wingreen, Z. Gitai, PSICIC: noise and asymmetry in bacterial division revealed by computational image analysis at sub-pixel resolution. *PLoS Comput Biol* **4**, e1000233 (2008).
5. N. E. Huang *et al.*, The empirical mode decomposition and the Hilbert spectrum for nonlinear and non-stationary time series analysis. *Proc. R. Soc. A* **454**, 903-995 (1998).
6. A. Paintdakhi *et al.*, Oufti: an integrated software package for high-accuracy, high-throughput quantitative microscopy analysis. *Mol. Microbiol.* **99**, 767-777 (2016).
7. W. T. Gray *et al.*, Nucleoid Size Scaling and Intracellular Organization of Translation across Bacteria. *Cell* **177**, 1632-1648 e1620 (2019).
8. S. C. Verma, Z. Qian, S. L. Adhya, Architecture of the Escherichia coli nucleoid. *PLoS Genet.* **15**, e1008456 (2019).
9. O. G. Stonington, D. E. Pettijohn, The folded genome of Escherichia coli isolated in a protein-DNA-RNA complex. *Proc Natl Acad Sci U S A* **68**, 6-9 (1971).
10. A. Worcel, E. Burgi, On the structure of the folded chromosome of Escherichia coli. *J. Mol. Biol.* **71**, 127-147 (1972).
11. L. P. Savtchenko, M. M. Poo, D. A. Rusakov, Electrodiffusion phenomena in neuroscience: a neglected companion. *Nat. Rev. Neurosci.* **18**, 598-612 (2017).
12. E. J. F. Dickinson, J. G. Limon-Petersen, R. G. Compton, The electroneutrality approximation in electrochemistry. *J. Solid State Electrochem.* **15**, 1335-1345 (2011).
13. K. R. Ward, E. J. F. Dickinson, R. G. Compton, How Far Do Membrane Potentials Extend in Space Beyond the Membrane Itself? *Int. J. Electrochem. Sci.* **5**, 1527-1534 (2010).
14. M. Marhl, M. Brumen, R. Glaser, R. Heinrich, Diffusion layer caused by local ionic transmembrane fluxes. *Pflugers Arch* **431**, R259-260 (1996).
15. E. J. Dickinson, L. Freitag, R. G. Compton, Dynamic theory of liquid junction potentials. *J. Phys. Chem. B* **114**, 187-197 (2010).
16. R. Shi, A. J. Cooper, H. Tanaka, Impact of hierarchical water dipole orderings on the dynamics of aqueous salt solutions. *Nat Commun* **14**, 4616 (2023).

17. S. Friesen, M. V. Fedotova, S. E. Kruchinin, R. Buchner, Hydration and dynamics of L-glutamate ion in aqueous solution. *Phys. Chem. Chem. Phys.* **23**, 1590-1600 (2021).
18. R. Mancinelli, A. Botti, F. Bruni, M. A. Ricci, A. K. Soper, Hydration of sodium, potassium, and chloride ions in solution and the concept of structure maker/breaker. *J. Phys. Chem. B* **111**, 13570-13577 (2007).
19. S. G. Schultz, A. K. Solomon, Cation transport in Escherichia coli. I. Intracellular Na and K concentrations and net cation movement. *J. Gen. Physiol.* **45**, 355-369 (1961).
20. S. G. Schultz, N. L. Wilson, W. Epstein, Cation transport in Escherichia coli. II. Intracellular chloride concentration. *J. Gen. Physiol.* **46**, 159-166 (1962).
21. L. Shabala *et al.*, Ion transport and osmotic adjustment in Escherichia coli in response to ionic and non-ionic osmotica. *Environ. Microbiol.* **11**, 137-148 (2009).
22. N. Amin, A. Peterkofsky, A dual mechanism for regulating cAMP levels in Escherichia coli. *J. Biol. Chem.* **270**, 11803-11805 (1995).
23. K. B. Xavier, M. Kossmann, H. Santos, W. Boos, Kinetic analysis by in vivo 31P nuclear magnetic resonance of internal Pi during the uptake of sn-glycerol-3-phosphate by the pho regulon-dependent Ugp system and the glp regulon-dependent GlpT system. *J. Bacteriol.* **177**, 699-704 (1995).
24. P. Gangola, B. P. Rosen, Maintenance of intracellular calcium in Escherichia coli. *J. Biol. Chem.* **262**, 12570-12574 (1987).
25. F. X. Theillet *et al.*, Physicochemical properties of cells and their effects on intrinsically disordered proteins (IDPs). *Chem Rev* **114**, 6661-6714 (2014).
26. C. E. Outten, T. V. O'Halloran, Femtomolar sensitivity of metalloregulatory proteins controlling zinc homeostasis. *Science* **292**, 2488-2492 (2001).
27. A. N. Woodmansee, J. A. Imlay, Reduced flavins promote oxidative DNA damage in non-respiring Escherichia coli by delivering electrons to intracellular free iron. *J. Biol. Chem.* **277**, 34055-34066 (2002).
28. B. D. Bennett *et al.*, Absolute metabolite concentrations and implied enzyme active site occupancy in Escherichia coli. *Nat. Chem. Biol.* **5**, 593-599 (2009).
29. J. Stautz *et al.*, Molecular Mechanisms for Bacterial Potassium Homeostasis. *J. Mol. Biol.* **433**, 166968 (2021).
30. B. Eisenberg, Interacting ions in biophysics: real is not ideal. *Biophys. J.* **104**, 1849-1866 (2013).
31. J. J. Jasielec, Electrodiffusion Phenomena in Neuroscience and the Nernst–Planck–Poisson Equations. *Electrochem* **2**, 197-215 (2021).
32. L. P. Savtchenko, N. Kulahin, S. M. Korogod, D. A. Rusakov, Electric fields of synaptic currents could influence diffusion of charged neurotransmitter molecules. *Synapse* **51**, 270-278 (2004).

33. C. L. Lopreore *et al.*, Computational modeling of three-dimensional electrodiffusion in biological systems: application to the node of Ranvier. *Biophys. J.* **95**, 2624-2635 (2008).
34. J. J. Jasielec *et al.*, Computer simulations of electrodiffusion problems based on Nernst–Planck and Poisson equations. *Comput. Mater. Sci.* **63**, 75-90 (2012).
35. K. Zhurov, E. J. Dickinson, R. G. Compton, Dynamics of ion transfer potentials at liquid-liquid interfaces. *J. Phys. Chem. B* **115**, 6909-6921 (2011).
36. G. Halnes *et al.*, Effect of Ionic Diffusion on Extracellular Potentials in Neural Tissue. *PLoS Comput Biol* **12**, e1005193 (2016).
37. K. R. Ward, E. J. F. Dickinson, R. G. Compton, Dynamic Theory of Membrane Potentials. *J. Phys. Chem. B* **114**, 10763-10773 (2010).
38. K. R. Ward, E. J. Dickinson, R. G. Compton, Dynamic theory of type 3 liquid junction potentials: formation of multilayer liquid junctions. *J. Phys. Chem. B* **114**, 4521-4528 (2010).
39. Z. Schuss, B. Nadler, R. S. Eisenberg, Derivation of Poisson and Nernst-Planck equations in a bath and channel from a molecular model. *Phys Rev E Stat Nonlin Soft Matter Phys* **64**, 036116 (2001).
40. C. Maffeo, S. Bhattacharya, J. Yoo, D. Wells, A. Aksimentiev, Modeling and simulation of ion channels. *Chem Rev* **112**, 6250-6284 (2012).
41. S. Bakshi, H. Choi, J. C. Weisshaar, The spatial biology of transcription and translation in rapidly growing Escherichia coli. *Front Microbiol* **6**, 636 (2015).
42. P. E. Schavemaker, W. M. Smigiel, B. Poolman, Ribosome surface properties may impose limits on the nature of the cytoplasmic proteome. *Elife* **6**, e30084 (2017).
43. V. Yeong, E. G. Werth, L. M. Brown, A. C. Obermeyer, Formation of Biomolecular Condensates in Bacteria by Tuning Protein Electrostatics. *ACS Cent Sci* **6**, 2301-2310 (2020).
44. W. M. Smigiel *et al.*, Protein diffusion in Escherichia coli cytoplasm scales with the mass of the complexes and is location dependent. *Sci Adv* **8**, eabo5387 (2022).
45. D. Valverde-Mendez *et al.*, Macromolecular interactions and geometrical confinement determine the 3D diffusion of ribosome-sized particles in live Escherichia coli cells. *Proc Natl Acad Sci U S A* **122**, e2406340121 (2025).
46. G. Gonella *et al.*, Water at charged interfaces. *Nat Rev Chem* **5**, 466-485 (2021).
47. A. B. Collis, P. R. Tulip, S. P. Bates, Structure and bonding of aqueous glutamic acid from classical molecular dynamics simulations. *Phys. Chem. Chem. Phys.* **12**, 5341-5352 (2010).
48. S. E. McLain, A. K. Soper, A. Watts, Structural studies on the hydration of L-glutamic acid in solution. *J Phys Chem B* **110**, 21251-21258 (2006).

49. I. Waluyo *et al.*, A different view of structure-making and structure-breaking in alkali halide aqueous solutions through x-ray absorption spectroscopy. *J. Chem. Phys.* **140**, 244506 (2014).
50. H. Ohshima, S. Ohki, Donnan potential and surface potential of a charged membrane. *Biophys. J.* **47**, 673-678 (1985).
51. P. W. C. Northrop, P. A. Ramachandran, W. E. Schiesser, V. R. Subramanian, A robust false transient method of lines for elliptic partial differential equations. *Chem. Eng. Sci.* **90**, 32-39 (2013).
52. P. Ortoleva, S. L. Schmidt, in *Oscillations and Traveling Waves in Chemical Systems,* R. J. Field, M. Burger, Eds. (1985), chap. 10, pp. 333-418.
53. P. Ortoleva, Chemical wave-electrical field interaction phenomena. *Physica D* **26**, 67-84 (1987).
54. L. L. Lackner, D. M. Raskin, P. A. de Boer, ATP-dependent interactions between Escherichia coli Min proteins and the phospholipid membrane in vitro. *J. Bacteriol.* **185**, 735-749 (2003).
55. Z. Hu, A. Mukherjee, S. Pichoff, J. Lutkenhaus, The MinC component of the division site selection system in Escherichia coli interacts with FtsZ to prevent polymerization. *Proc Natl Acad Sci U S A* **96**, 14819-14824 (1999).
56. D. M. Raskin, P. A. de Boer, MinDE-dependent pole-to-pole oscillation of division inhibitor MinC in Escherichia coli. *J. Bacteriol.* **181**, 6419-6424 (1999).
57. D. M. Raskin, P. A. de Boer, Rapid pole-to-pole oscillation of a protein required for directing division to the middle of Escherichia coli. *Proc Natl Acad Sci U S A* **96**, 4971-4976 (1999).
58. Z. Hu, E. P. Gogol, J. Lutkenhaus, Dynamic assembly of MinD on phospholipid vesicles regulated by ATP and MinE. *Proc Natl Acad Sci U S A* **99**, 6761-6766 (2002).
59. K. T. Park *et al.*, The Min oscillator uses MinD-dependent conformational changes in MinE to spatially regulate cytokinesis. *Cell* **146**, 396-407 (2011).
60. K. C. Huang, Y. Meir, N. S. Wingreen, Dynamic structures in Escherichia coli: spontaneous formation of MinE rings and MinD polar zones. *Proc Natl Acad Sci U S A* **100**, 12724-12728 (2003).
61. J. Halatek, E. Frey, Highly canalized MinD transfer and MinE sequestration explain the origin of robust MinCDE-protein dynamics. *Cell Rep* **1**, 741-752 (2012).
62. G. Meacci *et al.*, Mobility of Min-proteins in Escherichia coli measured by fluorescence correlation spectroscopy. *Phys Biol* **3**, 255-263 (2006).
63. E. Gianazza, P. Giorgio Righetti, Size and charge distribution of macromolecules in living systems. *J. Chromatogr. A* **193**, 1-8 (1980).

64. M.-S. Chun, I. Lee, Rigorous estimation of effective protein charge from experimental electrophoretic mobilities for proteomics analysis using microchip electrophoresis. *Colloids Surf. Physicochem. Eng. Aspects* **318**, 191-198 (2008).
65. D. Chen *et al.*, Predicting Electrophoretic Mobility of Proteoforms for Large-Scale Top-Down Proteomics. *Anal. Chem.* **92**, 3503-3507 (2020).
66. F. Wu, B. G. van Schie, J. E. Keymer, C. Dekker, Symmetry and scale orient Min protein patterns in shaped bacterial sculptures. *Nat Nanotechnol* **10**, 719-726 (2015).
67. J. Halatek, E. Frey, Rethinking pattern formation in reaction–diffusion systems. *Nature Physics* **14**, 507-514 (2018).
68. A. A. Tokmakov, A. Kurotani, K. I. Sato, Protein pI and Intracellular Localization. *Front Mol Biosci* **8**, 775736 (2021).
69. J. Slavik, A. Kotyk, Intracellular pH distribution and transmembrane pH profile of yeast cells. *Biochim Biophys Acta* **766**, 679-684 (1984).
70. J. Flegr, Does a cell perform isoelectric focusing? *Biosystems* **24**, 127-133 (1990).
71. E. M. Baskin, S. Bukshpan, G. V. Zilberstein, pH-induced intracellular protein transport. *Phys Biol* **3**, 101-106 (2006).
72. H. S. Fogler, *Elements of Chemical Reaction Engineering*. (Prentice Hall, ed. 5th, 2016).
73. O. A. Hougen, K. M. Watson, *Chemical Process Principles: Kinetics and Catalysis*. (John Wiley & Sons, 1947), vol. 3.
74. G. A. Somorjai, Y. Li, *Introduction to Surface Chemistry and Catalysis*. (John Wiley & Sons, ed. 2nd, 2010).
75. S. Cayley, B. A. Lewis, H. J. Guttman, M. T. Record, Characterization of the cytoplasm of Escherichia coli K-12 as a function of external osmolarity. *J. Mol. Biol.* **222**, 281-300 (1991).
76. S. B. Zimmerman, S. O. Trach, Estimation of macromolecule concentrations and excluded volume effects for the cytoplasm of Escherichia coli. *J. Mol. Biol.* **222**, 599-620 (1991).
77. Y. Ishihama *et al.*, Protein abundance profiling of the Escherichia coli cytosol. *BMC Genomics* **9**, 102 (2008).
